# High-avidity TCR signaling drives an NKG2A-positive CD8 T cell exhaustion state in human tumors

**DOI:** 10.64898/2026.05.29.728765

**Authors:** Riley D.Z. Mullins, Jesse M. Zaretsky, Thomas F. Barrett, Emily Stoller, Porter Bischoff, Ann Marie Egloff, Cody Huffman, Salma Ramadan, Amna Ramadan, Jessica Ley, Katherine Schwetye, Daniel Miller, Jennifer De Los Santos, Anthony J. Apicelli, Nikhil Rammohan, Wade Thorstad, Peter Oppelt, Brendan J. Knapp, Christine Auberle, Guangyong Peng, Craig Buchman, Andres Bur, Ben Wahle, Jason T. Rich, Ryan S. Jackson, Patrik Pipkorn, Paul A. Zolkind, Richard A. Harbison, Takeshi Egawa, Nathan Singh, Robi D. Mitra, Sana D. Karam, Douglas R. Adkins, Ravindra Uppaluri, Sidharth V. Puram

## Abstract

T cells adopt diverse states in human disease, with tumor-specific CD8 T cells serving as the quintessential example. These cells are critical for immunotherapy response, yet acquire hypofunctional states of exhaustion that impair treatment efficacy. While significant heterogeneity is observed among exhausted CD8 T cells (T_EX_), the mechanisms directing their unique phenotypes remain unresolved, limiting our efforts to augment T_EX_ cell function. Here, we profile T_EX_ cells in human head and neck squamous cell carcinoma (HNSCC) tumors and define a major mechanism regulating their cell state. With single-cell RNA and T cell receptor (TCR) sequencing of 106,667 tumor-infiltrating CD8 T cells, we define and validate three fundamental T_EX_ subsets, each harboring dominant clonotypes: a progenitor subset (T_PEX_) and two *intra*-tumoral populations marked by intermediate (T_EX-int_) or high expression of inhibitory receptors, including killer cell lectin-like receptors, such as *KLRC1*/NKG2A (T_EX-KLR_). Using an *in vitro* model, we show that TCR avidity-dependent NFAT signaling governs an axis between the T_EX-int_ and T_EX-KLR_ states across human cancers, with high-avidity signaling driving the T_EX-KLR_ state. Additionally, we uncover that interleukin-12 signaling selectively antagonizes high-avidity T_EX-KLR_-specific genes, despite broadly potentiating activation and cytotoxicity. Finally, we find that T_EX-int_ and T_EX-KLR_ cells are associated with anti-PD-1 response in HNSCC, with both of these cell types enriched near viable cancer cells following treatment. Together, these findings reveal that TCR avidity orchestrates tumor-infiltrating T_EX_ cell states through NFAT signaling – a tunable axis that may be targeted to improve T_EX_ cell function.

## INTRODUCTION

Although substantial progress has been made in surgery, radiation, and systemic therapy for solid tumors, immunotherapy arguably represents the most significant treatment advance in decades. Despite the promise of cancer immunotherapeutics, such as immune checkpoint blockade (ICB), durable tumor responses are only achieved in a minority of patients for most cancer types. For example, in head and neck squamous cell carcinoma (HNSCC), the objective response rate to anti-programmed death-1 (anti-PD-1) monotherapy is 20% in the recurrent/metastatic setting^1^ and 40–50% in the neoadjuvant setting^2–6^. Tumor-specific CD8 T cells are critical mediators of the anti-tumor immune response elicited by ICB^7–12^. However, due to chronic exposure to tumor-associated antigens, these cells adopt a hypofunctional state, termed exhaustion, that limits immunotherapy efficacy^13^. Single-cell sequencing of tumor-infiltrating lymphocytes (TILs) has revealed pronounced heterogeneity among exhausted CD8 T cells (T_EX_), yet T_EX_ subpopulations have been variously named across studies, including with numbers corresponding to clusters or descriptive names, such as “cytotoxic,” “dysfunctional,” or “extreme exhausted,” based on their relative expression of inhibitory receptors^2,11,13–16^. Furthermore, the mechanisms that direct T_EX_ cells into their unique cell states remain poorly defined. Thus, a comprehensive understanding of the flux between T_EX_ phenotypes observed across solid tumors may inform therapies that improve T_EX_ function and immunotherapy responses.

Here, we define a major mechanism controlling the inter-conversion of cell states in tumor-infiltrating T_EX_ cells using HNSCC as a model disease. By combining immunophenotyping with single-cell RNA sequencing (scRNA-seq) and single-cell T cell receptor (TCR) sequencing (scTCR-seq) of 106,667 tumor-infiltrating CD8 T cells, we define and validate three transcriptionally distinct T_EX_ subsets, each harboring dominant clonotypes: a progenitor subset (T_PEX_)^17,18^ and two *intra*-tumoral subsets marked by intermediate (T_EX-int_) or high expression of inhibitory receptors, including killer cell lectin-like receptors (KLR), such as *KLRC1*/NKG2A (T_EX-KLR_). With *in vitro* modeling of TCR avidity, we uncover that high-avidity TCR signaling regulates a reciprocal axis between the T_EX-int_ and T_EX-KLR_ states through nuclear factor of activated T cells (NFAT), in which high-avidity signaling drives the T_EX-KLR_ program. Furthermore, we reveal that interleukin-12 (IL-12) broadly potentiates activation but specifically inhibits the T_EX-KLR_-specific gene program induced by high-avidity TCR signaling. Finally, in HNSCC, we show that *intra*-tumoral infiltration of T_EX-int_ and T_EX-KLR_ cells is associated with anti-PD-1 response, with increases in these cells near viable cancer cells following therapy. Taken together, we demonstrate that TCR avidity directs tumor-infiltrating T_EX_ phenotypes via NFAT – a tunable axis that may be targeted to augment T_EX_ effector functions and potentially improve therapeutic responses.

## RESULTS

### HNSCC tumors contain three distinct subsets of exhausted CD8 T cells

We initially analyzed scRNA-seq and scTCR-seq data of CD8^+^ TILs from 32 treatment-naïve HNSCC patient tumors, including new data from 3 human papillomavirus-negative (HPV-) and 18 HPV-positive (HPV+) tumors (**Supplementary Table 1**), as well as 11 HPV-samples previously published by our group^19^. Among 106,667 CD8 T cells and 3,247 CD56^+^ CD8^-^ natural killer (NK) cells, we identified 14 scRNA-seq clusters, including memory and T_EX_ CD8 T cell populations, which were consistent with prior datasets (**Fig. 1a**, **Extended Data Fig. 1**, and **Supplementary Table 2**)^20^. Specifically, all T_EX_ cells expressed inhibitory receptors (*PDCD1*, *LAG3*, and *HAVCR2*), other classic exhaustion markers (*CXCL13*, *GZMB*, and *ENTPD1*), and tumor-reactive signatures (**Fig. 1b, c**; **Extended Data Fig. 1**; and **Supplementary Tables 3** and **4**)^7–12,21^. Among the total T_EX_ population, we identified three subtypes of exhaustion: T_PEX_ marked by *CCR6*, *LEF1*, and *TCF7*^17,18^; T_EX-int_ marked by intermediate expression of exhaustion genes and a subset of memory-associated genes (*EOMES, RYR2*, *GZMK*, and *GZMH*); and T_EX-KLR_ marked by high expression of exhaustion genes, KLR genes (*KLRC1*, *KLRC2*, and *KLRD1*), and other genes unique to this cluster, such as *KRT81*, *MYO1E*, and *CSF1* (**Fig. 1b** and **Supplementary Table 3**). Altogether, we define three transcriptionally distinct subsets of exhaustion in HNSCC tumors, each marked by gene programs beyond canonical exhaustion-associated genes.

**Figure 1.**
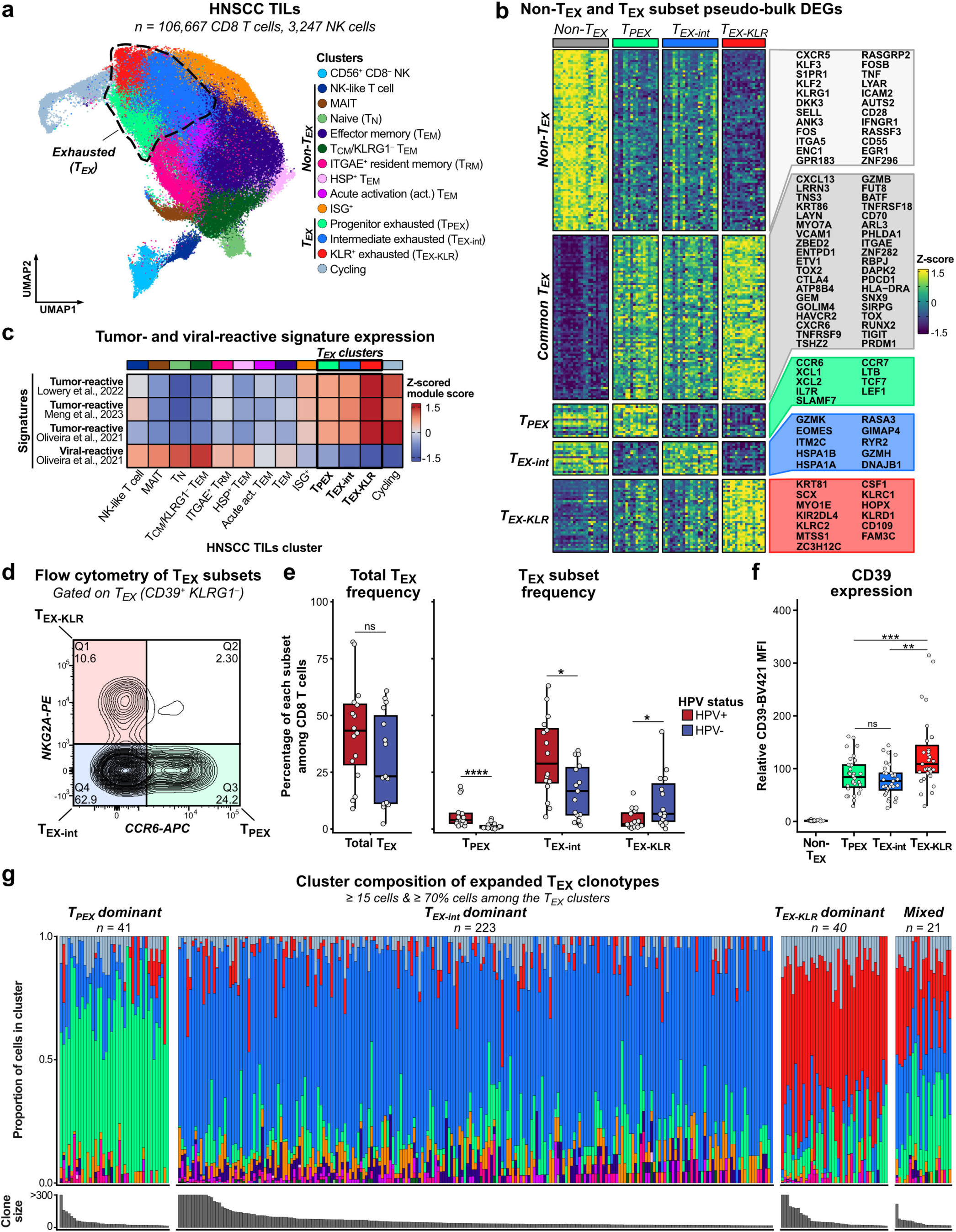
HNSCC tumors contain three distinct subsets of exhausted CD8 T cells. **a**, UMAP plot of the HNSCC TILs dataset. **b**, Z-scored pseudo-bulk RNA expression for DEGs between non-T_EX_ and T_EX_ subsets grouped by signature (see **Methods**). **c**, Mean z-scored module scores for tumor- and viral-reactive signatures in each CD8 T cell cluster. **d**, Representative flow cytometry plot of NKG2A and CCR6 expression on T_EX_ cells (CD8^+^ CD39^+^ KLRG1^-^). **e**, Percentage of total T_EX_ cells (*left*) and each T_EX_ subset (*right*) among total CD8 T cells in the flow cytometry data (*n* = 16 HPV+ and 15 HPV-tumors). **f**, CD39 mean fluorescence intensity (MFI) of the T_EX_ subsets relative to non-T_EX_ cells (*n* = 31 tumors). **g**, scRNA-seq cluster composition of expanded T_EX_ clonotypes grouped by their dominant T_EX_ cluster and ordered by descending cell number. *P* values from a Wilcoxon rank-sum test with Benjamini–Hochberg correction are shown in (**e**, **f**).

We next validated these T_EX_ subsets by flow cytometry in an independent cohort of 16 HPV+ HNSCC samples and 15 HPV-HNSCC samples (**Fig. 1d–f** and **Supplementary Table 1**). The total T_EX_ population, which was marked by CD39 (*ENTPD1*), was separated into three exclusive subsets by NKG2A (*KLRC1*) and CCR6: T_PEX_ (NKG2A^-^ CCR6^+^), T_EX-int_ (NKG2A^-^ CCR6^-^), and T_EX-KLR_ (NKG2A^+^ CCR6^-^; **Fig. 1d** and **Extended Data Fig. 2a**). While all three T_EX_ subsets were detected across samples, HPV+ tumors had a higher fraction of T_PEX_ and T_EX-int_ cells than HPV-tumors, which instead had a higher proportion of T_EX-KLR_ cells among CD8 T cells (**Fig. 1e**). Additionally, HPV+ tumors tended to have a higher fraction of CD8 T cells among all T cells, while HPV-tumors were relatively enriched for CD4 T cells (**Extended Data Fig. 2b, c**). Next, to confirm these populations were functionally exhausted, we stimulated T_EX-int_ and T_EX-KLR_ cells sorted from freshly dissociated tumors with anti-CD3 *in vitro* and found that both populations exhibited a lack of proliferation and activation, cardinal features of exhaustion; there were insufficient T_PEX_ cells for analysis (**Extended Data Fig. 3**). The flow cytometry T_EX-KLR_ population also had the highest expression of CD39, matching the scRNA-seq T_EX-KLR_ cluster (**Fig. 1f**). In addition to their distinct transcriptional profiles and surface protein markers, these T_EX_ subsets harbored unique, dominant clonotypes (**Fig. 1g**). Among 325 expanded T_EX_ clonotypes, 304 were dominant in a single T_EX_ subset (93.5%), defined as at least half of a clonotype’s cells in a single T_EX_ cluster, suggesting a TCR-intrinsic bias in phenotype (**Fig. 1g** and **Extended Data Fig. 4**). Most clonotypes were predominantly in the T_EX-int_ subset (*n* = 223, 68.6%), followed by the T_PEX_ (*n* = 41, 12.6%) and T_EX-KLR_ (*n* = 40, 12.3%) subsets (**Fig. 1g** and **Extended Data Fig. 4**). In summary, we identify and validate three transcriptionally and clonally distinct T_EX_ subsets distinguishable by CD39, NKG2A, and CCR6 in HPV- and HPV+ HNSCC.

### Exhausted CD8 T cell subsets are uniquely localized in tumors

Given the distinct transcriptional signatures of the T_EX_ subsets, we asked whether these populations were uniquely localized in tumors. To investigate this, we performed multiplexed immunofluorescence staining on 11 treatment-naïve HNSCC tumors (**Supplementary Table 1**), in which we identified cancer cells, B cells, NK cells, macrophages, and detailed CD4 and CD8 T cell subsets (**Fig. 2a, b** and **Extended Data Fig. 5**). Among CD8 T cells, T_PEX_ cells were marked by expression of TOX, TCF-1 (*TCF7*), and PD-1, as classically defined, whereas T_EX-int_ and T_EX-KLR_ cells were captured as TOX^+^ or PD-1^+^ and TCF-1^-^, with T_EX-KLR_ cells additionally expressing NKG2A (**Fig. 2b–d** and **Extended Data Fig. 5**). The CD8 T cell naïve (T_N_; TCF-1^+^ CD45RO^-^), central memory (T_CM_; TCF-1^+^ CD45RO^+^), effector memory (T_EM_; GZMB^-^ GZMK-high), and effector memory-intermediate (T_EM-int_; GZMB^+^ GZMK-high) populations were grouped as non-T_EX_ cells. We quantified the relative localization of each immune cell type in the tumor as an odds ratio of its frequency among all cells in the *intra*-tumoral region relative to the *peri*-tumoral region, the latter defined as the 200 μm of stroma extending outward from the tumor–stroma interface (**Fig. 2e**). We found that only T_EX-KLR_ cells preferentially localized *intra*-tumorally, and T_EX-int_ cells were approximately equally distributed across both regions, while T_PEX_ cells and all other T cell types were predominantly in the *peri*-tumoral region (**Fig. 2e**). As a proportion of total CD8 T cells, T_EX-int_ cells were the most frequent T_EX_ subset and T_PEX_ cells the least frequent in both regions (**Fig. 2f**). Although T_EX-KLR_ cells were infrequent *peri*-tumorally, they were relatively enriched *intra*-tumorally in each patient, constituting a median of 8% of the *intra*-tumoral CD8 T cell population, ranging from 0.5% to 27.5% (**Fig. 2f**). Consistent with these distributions, T_EX-KLR_ cells neighbored the most cancer cells of any T_EX_ subset, suggesting that their greater expression of exhaustion genes could reflect TCR-mediated activation through interaction with target cancer cells (**Fig. 1b** and **Fig. 2g**). Notably, T_EX-int_ and T_EX-KLR_ cells were often adjacent in the *intra*-tumoral region, implying that local environmental signals are unlikely to fully account for induction of these states (**Fig. 2c, d**).

**Figure 2.**
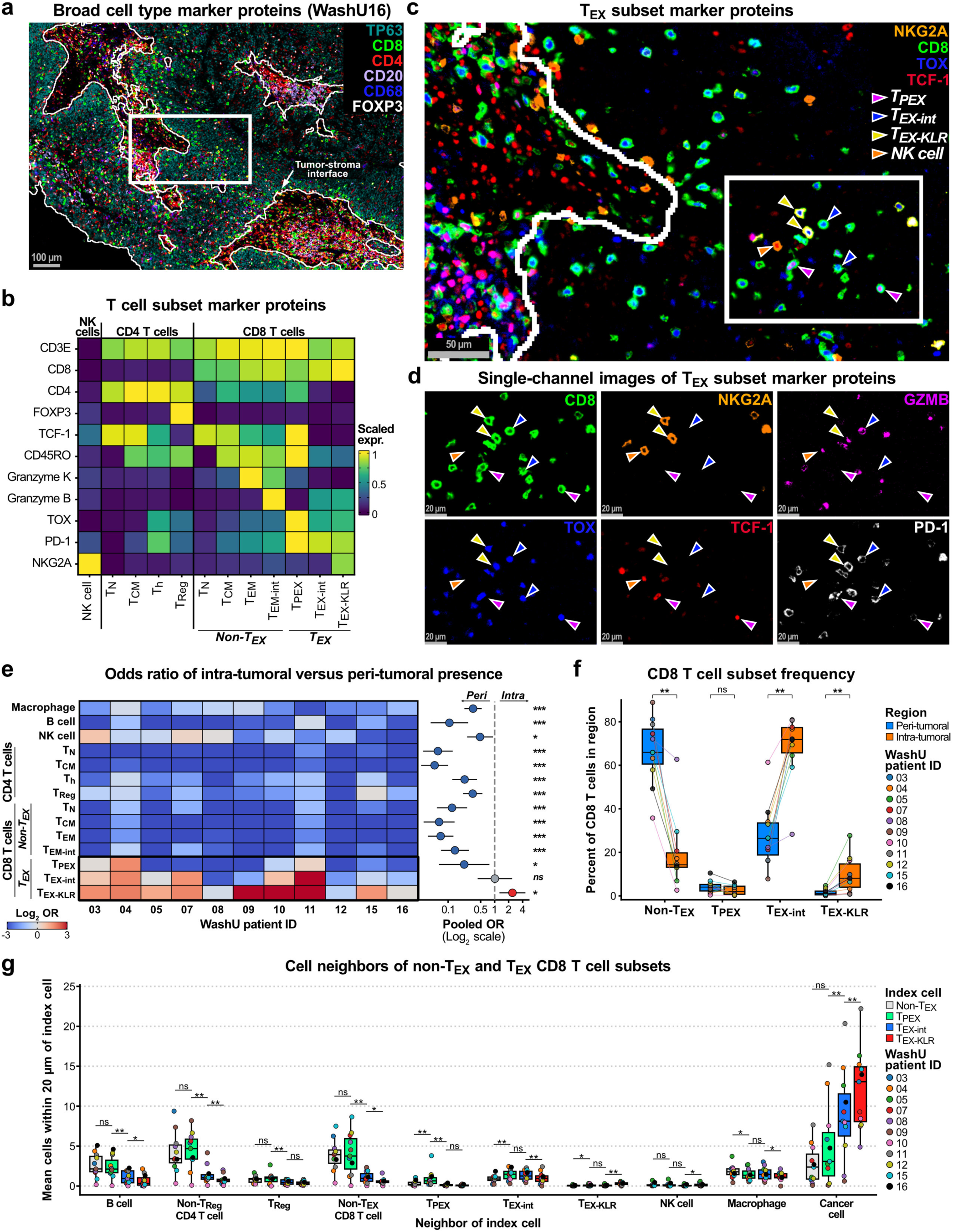
Exhausted CD8 T cell subsets are uniquely localized in tumors. **a**, Representative immunofluorescence image showing the tumor–stroma interface and TP63, marking malignant cells, and broad immune cell type marker proteins, including CD8, CD4, CD20, CD68, and FOXP3. **b**, Expression of T cell marker proteins in CD4 and CD8 T cell subsets and NK cells. **c**, Boxed region of (**a**) showing T_EX_ subset marker proteins, including NKG2A, CD8, TOX, and TCF-1, with selected T_PEX_, T_EX-int_, T_EX-KLR_, and NK cells indicated by colored arrows. **d**, Boxed region of (**c**) showing single-channel images of T_EX_ subset marker proteins. **e**, Log_2_ odds ratio (OR) for the frequency of each cell type among all cells in the intra-tumoral relative to the peri-tumoral region per patient (*left*) and the pooled OR with its 95% confidence interval (*right*). **f**, Percentage of each CD8 T cell subset among all CD8 T cells within the peri- and intra-tumoral regions. **g**, Mean count of each cell type neighboring non-T_EX_, T_PEX_, T_EX-int_, and T_EX-KLR_ CD8 T cells within 20 μm. *P* values from a Wald test of the pooled OR (**e**) and a Wilcoxon signed-rank test (**f**, **g**) with Benjamini–Hochberg correction are shown (n = 11 tumors each).

### High-avidity TCR signaling promotes NKG2A expression

Given the *intra*-tumoral enrichment of T_EX-KLR_ cells, their dominant clonotypes, and their greater expression of exhaustion genes, we hypothesized that high-avidity TCR signaling drives the T_EX-KLR_ state. To test this, we developed an *in vitro* system to model activation elicited by different TCR avidities and measured NKG2A as a marker of the T_EX-KLR_ state (**Fig. 3a**). This system was based on co-culture of an HPV16-positive HNSCC cell line, UM-SCC47, with primary peripheral human CD8 T cells transduced with a TCR that binds the HPV16 E7_11–20_ peptide (YMLDLQPETT) presented by HLA-A2^22^. We selected the UM-SCC47 cell line because it naturally expresses the HPV16 E7_11–20_ target peptide but does not express HLA-A2, allowing us to modify both components of avidity: surface antigen load and the TCR-peptide interaction. As negative controls, we used empty vector transduction with or without E7_11–20_ peptide pulsing. To model natural presentation of the E7_11–20_ peptide, we overexpressed HLA-A2, also with or without supplemental peptide pulsing. For constitutively high surface antigen presentation, we created and overexpressed a single-chain trimer (SCT) made of HLA-A2 fused to β2-microglobulin and the E7_11–20_ peptide, similar to previously reported SCT designs^23^. Lastly, to perturb the TCR-peptide interaction directly, we generated three mutants of the wildtype (WT) HPV16 E7_11–20_ peptide in the SCT that each differed by only a single charge-altering amino acid substitution: glutamine to glutamate (Q6E), glutamate to arginine (E8R), or glutamate to lysine (E8K). Following 48-hour co-culture, the CD8 T cells transduced with the E7_11–20_-specific TCR (E7 TCR^+^ CD8 T cells) were collected and flow cytometry was performed to measure their expression of CD3, CD137, and NKG2A (**Extended Data Fig. 6**).

**Figure 3.**
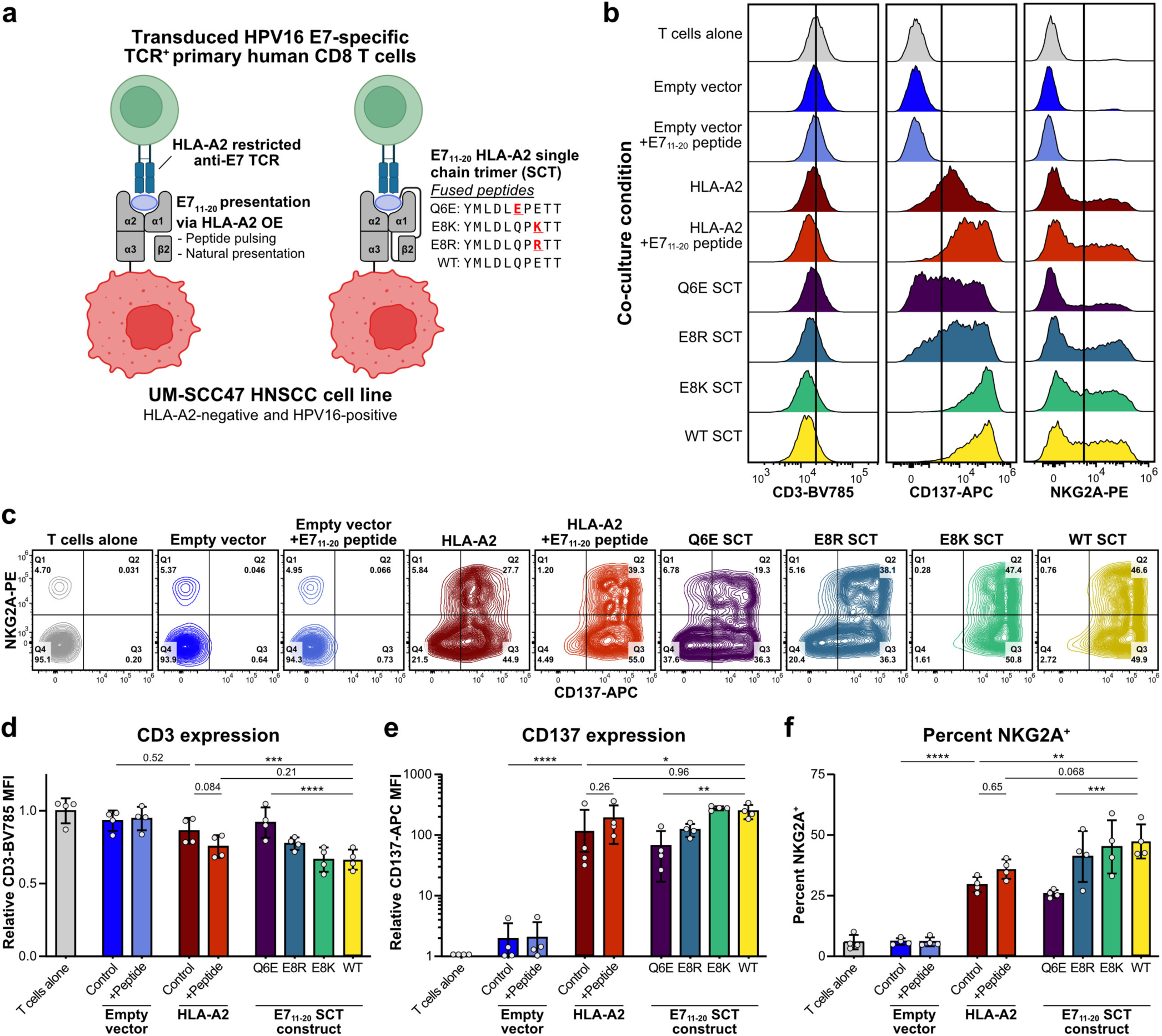
High-avidity TCR signaling promotes NKG2A expression. **a**, Schematic of UM-SCC47 target cell lines, including HLA-A2 overexpression (OE; *left*) with or without peptide pulsing and the WT and mutant E7_11–20_ SCT cell lines (*right*). **b–c**, Representative flow cytometry (**b**) histograms of CD3, CD137, and NKG2A expression normalized to the mode and (**c**) plots of NKG2A and CD137 expression on E7 TCR^+^ CD8 T cells following 48 h co-culture in each condition. Data for E7 TCR^+^ CD8 T cells generated from donor LRS071724 are shown. **d–f**, Quantification of (**d**) CD3 and (**e**) CD137 mean fluorescence intensity (MFI) relative to E7 TCR^+^ CD8 T cells cultured alone and (**f**) percentage of NKG2A^+^ E7 TCR^+^ CD8 T cells (*n* = 4 donors; E7 TCR^+^ CD8 T cells generated from donors LRS050924, LRS071724, LRS081224, and LRS081524 were used). *P* values from a repeated-measures one-way ANOVA with Sidak’s correction are shown in (**d–f**).

The E7 TCR^+^ CD8 T cells exhibited avidity-dependent expression of NKG2A, CD3, and CD137, the latter two serving as activation markers where higher CD137 and lower CD3 indicate greater activation (**Fig. 3b, c**). When co-cultured with HLA-A2 UM-SCC47 cells, the E7 TCR^+^ CD8 T cells moderately upregulated CD137 and NKG2A while decreasing CD3, with 28–34% NKG2A^+^ cells, an increase from the baseline 3–11% NKG2A^+^ fraction (**Fig. 3d–f**). Supplemental peptide pulsing, which increases the amount of presented surface antigen, modestly raised the NKG2A^+^ fraction to 31–38% (**Fig. 3f**). Co-culture with UM-SCC47 cells expressing the WT SCT, which confers a greater presented surface antigen load than HLA-A2 overexpression, further increased the NKG2A^+^ fraction to 42–58% alongside an increase in CD137 and decrease in CD3 (**Fig. 3d–f**). Direct perturbation of the TCR-peptide interaction with the Q6E SCT mutant lowered activation (**Fig. 3d, e**) and reduced the NKG2A^+^ fraction to 23–28% (**Fig. 3f**). Surface expression of the Q6E SCT and WT SCT was approximately equal, supporting that the observed effects were due to disruption of the TCR-peptide interaction by the Q6E mutation rather than the level of presented surface antigen (**Extended Data Fig. 6b**). The E8K mutation did not substantially alter these markers, but the E8R mutation moderately reduced activation and NKG2A expression relative to the WT SCT; however, surface expression of these mutants was lower than that of the WT SCT (**Fig. 3d–f** and **Extended Data Fig. 6b**). We next repeated these experiments in a second, non-malignant cell line – human embryonic kidney 293T (HEK293T) – that naturally expresses HLA-A2 but is not infected with HPV16 (**Extended Data Fig. 7**). We obtained similar results in this cell line, indicating that the avidity-dependent induction of NKG2A is robust to cellular context: the WT SCT induced higher NKG2A expression than any other condition, and the Q6E substitution attenuated NKG2A expression and activation (**Extended Data Fig. 7d–f**). Collectively, these data indicate that NKG2A expression on activated CD8 T cells is directly related to TCR avidity, specifically the abundance of presented surface antigen and the TCR-peptide interaction.

### NFAT-dependent high-avidity TCR signaling induces NKG2A

We next sought to identify the signaling pathway mediating the avidity-dependent NKG2A expression and to determine whether that pathway also drives the broader T_EX-KLR_ gene program observed *in vivo*. IL-12 has been reported to potentiate T cell activation and induce NKG2A in CD8 T cells^24–26^, an effect that may arise from its signaling through AP-1 and STAT4^26–28^. However, the concentration of IL-12p70, the active form of IL-12, was negligible in the media following a 48-hour co-culture (< 4 pg mL^−1^), ruling out IL-12 as responsible for NKG2A induction in our model (**Extended Data Fig. 6c**). This suggested that high-avidity TCR signaling is sufficient to drive NKG2A expression through an IL-12-independent mechanism. Given that NFAT activity increases with TCR avidity^29^, we hypothesized that NFAT regulates NKG2A and the broader *in vivo* T_EX-KLR_ state.

To directly compare IL-12 and NFAT signaling, we performed flow cytometry and bulk RNA-seq on E7 TCR^+^ CD8 T cells co-cultured for 48 hours with Q6E SCT, HLA-A2, or WT SCT UM-SCC47 target cells under treatment with FK506 to inhibit calcineurin–NFAT signaling^30^ or IL-12p70 to activate IL-12 signaling (**Fig. 4a**). We also tested anti-CD3 and anti-CD28 stimulation by flow cytometry but omitted this condition from the bulk RNA-seq because it did not lead to induction of NKG2A, suggesting that TCR-mediated activation on a target cell may have distinct signaling that is absent from antibody-based activation (**Extended Data Fig. 8a, b**). Consistent with prior literature, IL-12 increased the fraction of NKG2A^+^ cells in addition to modestly reducing CD137 expression compared with the vehicle control in co-cultures with each target cell line (**Fig. 4b, c** and **Extended Data Fig. 8a**). FK506 blocked avidity-related NKG2A induction, indicating that calcineurin–NFAT signaling positively regulates NKG2A, and this was accompanied by a large reduction in the percentage of CD137^+^ cells in all co-culture conditions (**Fig. 4b, c**). Surprisingly, IL-12 rescued NKG2A expression in an NFAT-*independent* manner, inducing it even with concomitant FK506 treatment (**Fig. 4b, c**). In addition to promoting NKG2A expression, IL-12 increased cytotoxicity independent of NFAT, overcoming the deficit in cytotoxicity caused by NFAT inhibition (**Extended Data Fig. 8c**). Overall, these data demonstrate that NFAT and IL-12 signaling upregulate NKG2A through independent mechanisms, each associated with functional impacts on cytotoxicity.

**Figure 4.**
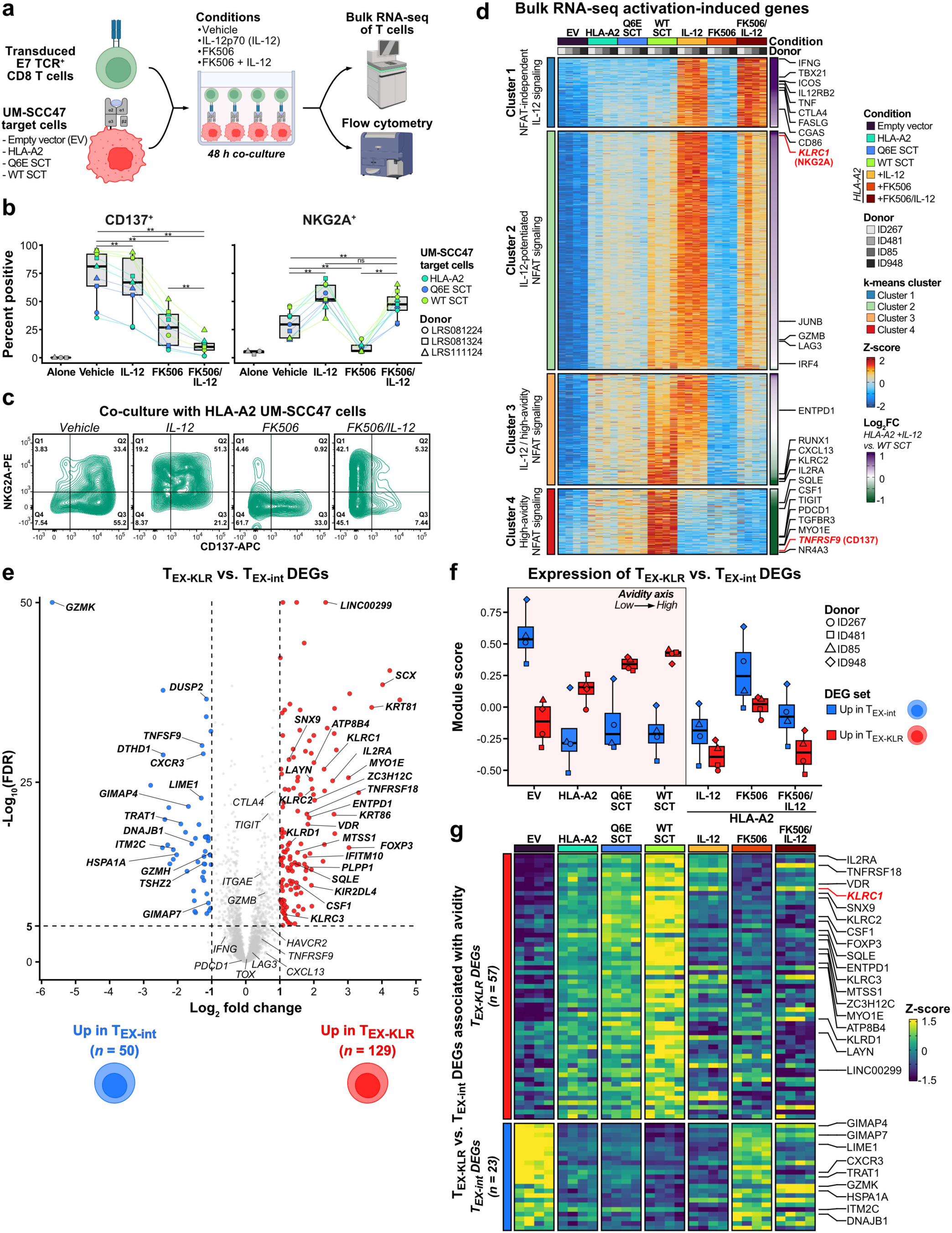
NFAT-dependent high-avidity TCR signaling induces NKG2A and the T_EX-KLR_ state. **a**, Schematic of experimental design. E7 TCR^+^ CD8 T cells were co-cultured for 48 h with UM-SCC47 target cells under treatment with vehicle, IL-12p70 (IL-12), FK506, or FK506 and IL-12, followed by flow cytometry and bulk RNA-seq (*n* = 3 donors for flow cytometry and 4 donors for bulk RNA-seq). **b**, Percentage of CD137^+^ (*left*) and NKG2A^+^ (*right*) E7 TCR^+^ CD8 T cells following 48 h co-culture in each condition (*n* = 3 donors in each of 3 co-culture conditions). *P* values from a Wilcoxon signed-rank test using all 9 data points in each treatment condition with Benjamini–Hochberg correction are shown. **c**, Representative flow cytometry plots of NKG2A and CD137 expression on E7 TCR^+^ CD8 T cells following 48 h co-culture. Data for E7 TCR^+^ CD8 T cells generated from donor LRS081324 are shown. **d**, Z-scored RNA expression of activation-induced genes, k-means clustered and ordered by descending log_2_FC between co-culture with HLA-A2 target cells plus IL-12 and co-culture with WT SCT target cells. **e**, Pseudo-bulk DEGs between the T_EX-KLR_ and T_EX-int_ scRNA-seq clusters (*n* = 17 patients represented in both clusters, each contributing a pseudo-bulk profile). **f**, Module scores for T_EX-int_ versus T_EX-KLR_ DEGs in the bulk RNA-seq samples. The light red overlay indicates the avidity-axis samples. **g**, Z-scored RNA expression of avidity-associated T_EX-int_ and T_EX-KLR_ DEGs in the bulk RNA-seq samples.

### TCR avidity, IL-12, and NFAT signaling activate both shared and distinct gene expression programs

To define the gene programs regulated by TCR avidity, IL-12, and NFAT signaling, we performed k-means clustering on activation-induced genes, defined as those upregulated in E7 TCR^+^ CD8 T cells co-cultured with HLA-A2 target cells plus IL-12 or with WT SCT target cells (log_2_ fold change [log_2_FC] > 1 and adjusted *P* < 0.01 versus the empty vector control; **Fig. 4d** and **Supplementary Table 5**). We defined four k-means clusters varying in their regulation by TCR avidity, IL-12, and NFAT (**Fig. 4d** and **Supplementary Table 6**). Cluster 1 genes (*n* = 246) were induced by NFAT-independent IL-12 signaling and included canonical markers of IL-12-enhanced TCR signaling, such as *IFNG*, *IL12RB2*, *TBX21*, and *TNF*^31^. Cluster 2 genes (*n* = 845) were potentiated by IL-12 in an NFAT-sensitive manner, indicating additive effects of NFAT and IL-12 signaling, and included *KLRC1*, *GZMB*, and *JUNB*. Cluster 3 genes (*n* = 394), such as *CXCL13* and *ENTPD1*, were regulated similarly to those of Cluster 2 but displayed greater induction by high-avidity TCR signaling and higher sensitivity to NFAT inhibition. Lastly, Cluster 4 genes (*n* = 232) were increased by NFAT-dependent high-avidity TCR signaling yet *suppressed* by IL-12. Notably, this cluster included genes marking the T_EX-KLR_ scRNA-seq subset, such as *CSF1*, *MYO1E*, and *TNFRSF9*, suggesting that high-avidity TCR signaling may promote the T_EX-KLR_ state. Together, these data demonstrate that although IL-12 and NFAT signaling both induce NKG2A (*KLRC1*), they act on shared and distinct gene expression programs, with genes of the T_EX-KLR_ program induced by NFAT and inhibited by IL-12.

### NFAT-dependent high-avidity TCR signaling drives the T_EX-KLR_ state

We next formally assessed how the T_EX-KLR_ signature seen in human TILs was affected by TCR avidity, IL-12, and NFAT signaling. Specifically, we created gene sets from the differentially expressed genes (DEGs) between the T_EX-KLR_ and T_EX-int_ scRNA-seq clusters (|log_2_FC| > 1 and adjusted *P* < 1 × 10^−5^) and applied them to the bulk RNA-seq dataset as module scores (**Fig. 4e** and **Supplementary Table 7**). We found that the T_EX-int_ DEG module score (*n* = 50 genes) was suppressed by *in vitro* activation and rescued by NFAT inhibition (**Fig. 4f**). Of these genes, 23 (46%) were negatively correlated with avidity *in vitro* (**Fig. 4g** and **Supplementary Table 7**). These included *GZMK*, *ITM2C*, *GIMAP4,* and other genes expressed in both the memory and the T_EX-int_ scRNA-seq clusters, which suggests that NFAT substantially represses memory-like genes expressed in T_EX_ cells upon activation (**Fig. 1b** and **Extended Data Fig. 1d**). Conversely, the T_EX-KLR_ DEG module score (*n* = 129 genes) was promoted in an avidity- and NFAT-dependent manner, suggesting that NFAT-dependent high-avidity TCR signaling drives the T_EX-KLR_ state (**Fig. 4f** and **Supplementary Table 7**). Of the T_EX-KLR_ DEGs, 57 (44%) were positively correlated with avidity *in vitro*, including *KLRC1*, *KLRC2*, *KLRD1*, and *CSF1* (**Fig. 4g** and **Supplementary Table 7**). The T_EX-int_ and T_EX-KLR_ DEGs that were not associated with avidity *in vitro* likely reflect additional *in vivo*-specific signals. Strikingly, while IL-12 signaling upregulated many activation-induced genes (**Fig. 4d**), it selectively suppressed the T_EX-KLR_ signature to an even greater extent than NFAT inhibition via FK506 treatment (**Fig. 4f, g**). We therefore directly compared the effects of IL-12 and high-avidity TCR signaling on activation by defining genes upregulated in E7 TCR^+^ CD8 T cells co-cultured with HLA-A2 target cells plus IL-12 or with WT SCT target cells, each relative to the HLA-A2 plus vehicle condition as the baseline-activation reference (log_2_FC > 0.5 and adjusted *P* < 0.05 in either contrast). These genes were labeled as IL-12- or avidity-dominant if the difference in log_2_FC between these contrasts exceeded 0.75 or as shared genes if the log_2_FC was greater than 0.5 in each contrast (**Extended Data Fig. 9a**; see **Methods**). We identified 117 IL-12-dominant, 151 avidity-dominant, and 164 shared genes (**Supplementary Table 8**). Of the avidity-dominant genes, IL-12 significantly suppressed 61 (40%; log_2_FC < −0.5 and adjusted *P* < 0.05), including *NR4A1*, *NR4A3*, *MYO1E*, *SLA, TIGIT,* and *TNFRSF9*, which suggests that IL-12 antagonizes a high-avidity, T_EX-KLR_-specific gene program. Indeed, the scRNA-seq T_EX-KLR_ cluster was marked by expression of the avidity-dominant genes, but not the IL-12-dominant or shared genes, supporting that high-avidity TCR signaling promotes the T_EX-KLR_ state (**Extended Data Fig. 9b–e**). Overall, these data demonstrate that NFAT-dependent high-avidity TCR signaling induces the T_EX-KLR_ signature and represses memory-like genes associated with the T_EX-int_ subset.

### Exhausted clonotypes occupy a reciprocal axis between T_EX-int_ and T_EX-KLR_ states across human tumors

We next applied the T_EX-int_ and T_EX-KLR_ DEGs that were associated with avidity *in vitro* (defined in **Fig. 4g**) as module scores to T_EX_ cells from 13 human cancer types in the scRNA-seq human Antigen Receptor database (huARdb)^32^ (**Fig. 5a**). We found that distinct T_EX_ subclusters variably expressed the avidity-associated T_EX-int_ and T_EX-KLR_ DEG signatures in a reciprocal, anti-correlated fashion (**Fig. 5b**). Additionally, in the huARdb and our HNSCC TILs dataset, T_EX_ clonotypes were stratified along an axis of avidity-associated T_EX-int_ and T_EX-KLR_ DEG signature expression, with decreased expression of one associated with gain of the other, implying a clonal bias in cell state related to avidity-dependent NFAT signaling (**Fig. 5c** and **Extended Data Fig. 10**). Because the dynamics of avidity require a TCR-peptide interaction, we hypothesized that the variable expression of the T_EX-int_ and T_EX-KLR_ DEG signatures across clonotypes may be partially explained by target cell contact. To investigate this, we analyzed a dataset of melanoma CD8^+^ TILs sorted as singlets or as doublets engaged with an antigen-presenting cell (APC) or cancer cell (**Fig. 5d**)^33^. Consistent with the T_EX-KLR_ state reflecting TCR signaling, CD8 T cells engaged with an APC or cancer cell had higher relative expression of the avidity-associated T_EX-KLR_ DEG signature compared with singlets (**Fig. 5e, f**). This was also apparent per clonotype; cells of the same clonotype that were sorted as doublets had, on average, higher relative expression of the T_EX-KLR_ signature than their singlet counterparts (**Fig. 5g**). Together, these data support that TCR avidity-dependent NFAT signaling regulates an axis of T_EX-int_ and T_EX-KLR_ states in tumor-infiltrating T_EX_ cells.

**Figure 5.**
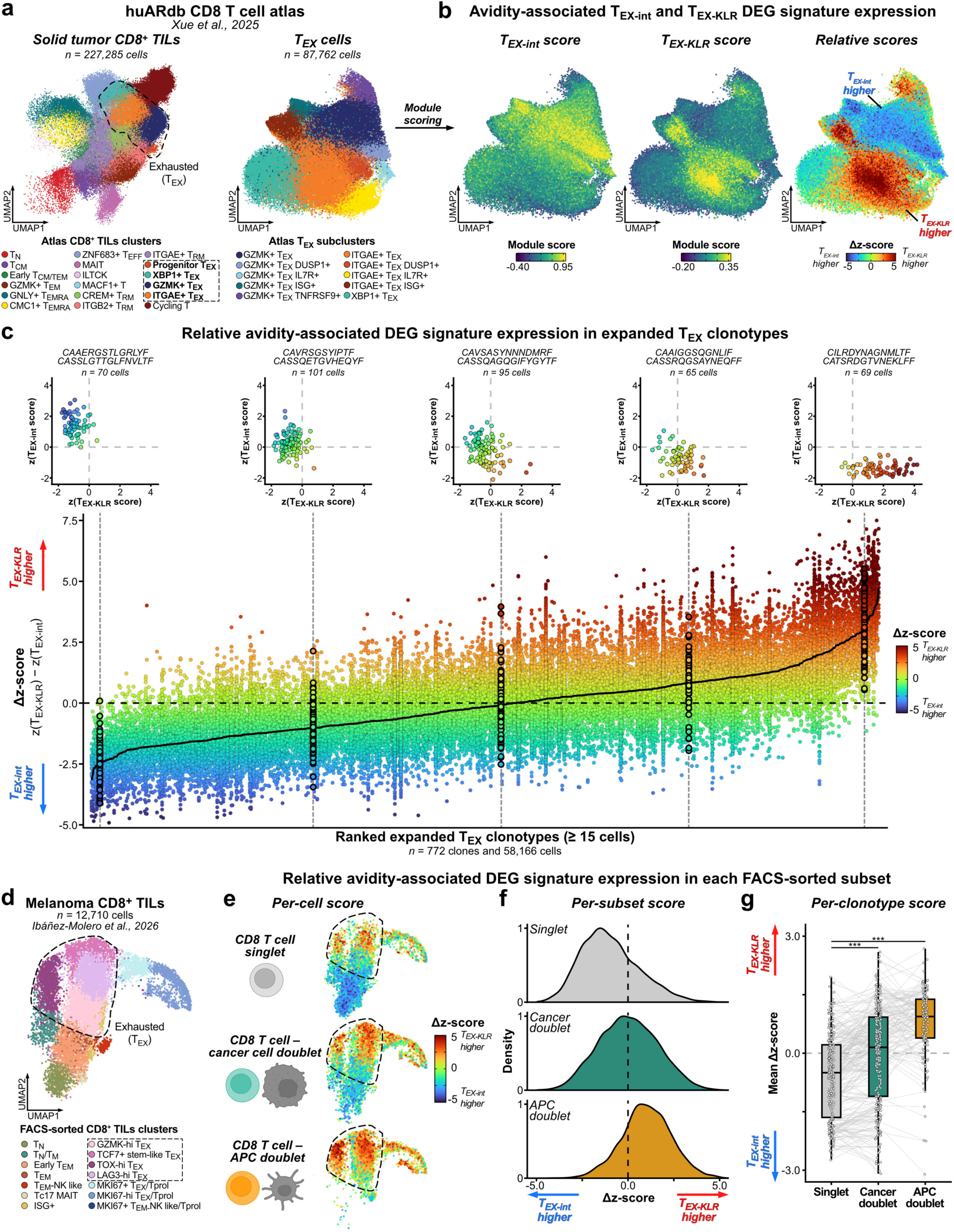
Exhausted clonotypes occupy a reciprocal axis between T_EX-int_ and T_EX-KLR_ states across human tumors. **a**, UMAP plot of solid tumor CD8^+^ TILs for all clusters (*left*) and T_EX_ subclusters (*right*) from the huARdb dataset; the published UMAP coordinates and clusters are shown (Xue *et al*., 2025). **b**, Module scores for the avidity-associated T_EX-int_ (*left*) and T_EX-KLR_ (*middle*) DEG signatures (from Fig. 4g) and the relative module scores represented as the Δz-score (*right*; Δz-score = z-scored T_EX-KLR_ module score minus z-scored T_EX-int_ module score). **c**, Relative avidity-associated DEG signature module scores in expanded T_EX_ clonotypes, where cells of each clonotype are arranged into columns and filled by their Δz-score, and clonotypes are ordered by their mean Δz-score. Cells of highlighted clonotypes are shown on a scatter plot. **d**, UMAP plot of analyzed melanoma CD8^+^ TILs (Ibáñez-Molero *et al*., 2026), including cells from clonotypes with at least three cells in each of at least two FACS-sorted subsets; the published UMAP coordinates and clusters are shown. **e–g**, Relative avidity-associated DEG signature module scores represented as the (**e**) per-cell, (**f**) per-subset, and (**g**) per-clonotype Δz-score, where lines in (**g**) connect the same clonotype across different subsets (*n* = 242 paired clonotypes in singlet and cancer cell doublet subsets and *n* = 82 in singlet and APC doublet subsets). *P* values from a Wilcoxon signed-rank test with Benjamini–Hochberg correction are shown.

### T_EX-int_ and T_EX-KLR_ cells are associated with anti-PD-1 response and increase in areas of viable tumor post-treatment

Given the association between the T_EX-KLR_ state and high-avidity TCR signaling, we asked whether T_EX-KLR_ cells are associated with anti-PD-1 response. We investigated this in HNSCC patients who received neoadjuvant anti-PD-1 therapy: one dose of pembrolizumab 2–3 weeks prior to surgery (Cohort 1) or two doses over 5–6 weeks (Cohort 2; **Fig. 6a**)^2,34^. The surgical specimen was assessed for pathologic tumor response (pTR), where pTR2 corresponds to ≥ 50% tumor regression, pTR1 to 10–49%, and pTR0 to < 10%^2^. For Cohort 1, we performed multiplexed immunofluorescence staining on a tissue microarray of pre- and post-treatment samples (*n* = 16 pTR0, 6 pTR1, and 5 pTR2 patient tumor samples). For Cohort 2, which was previously assessed for broad cell types^19^, we performed new comprehensive T cell immunophenotyping on whole-tissue sections from two non-responders (pTR0) and two responders (pTR2). In both cohorts, broad cell types and detailed CD8 T cell subsets were defined by the same criteria used before (**Fig. 2** and **Extended Data Fig. 5**).

**Figure 6.**
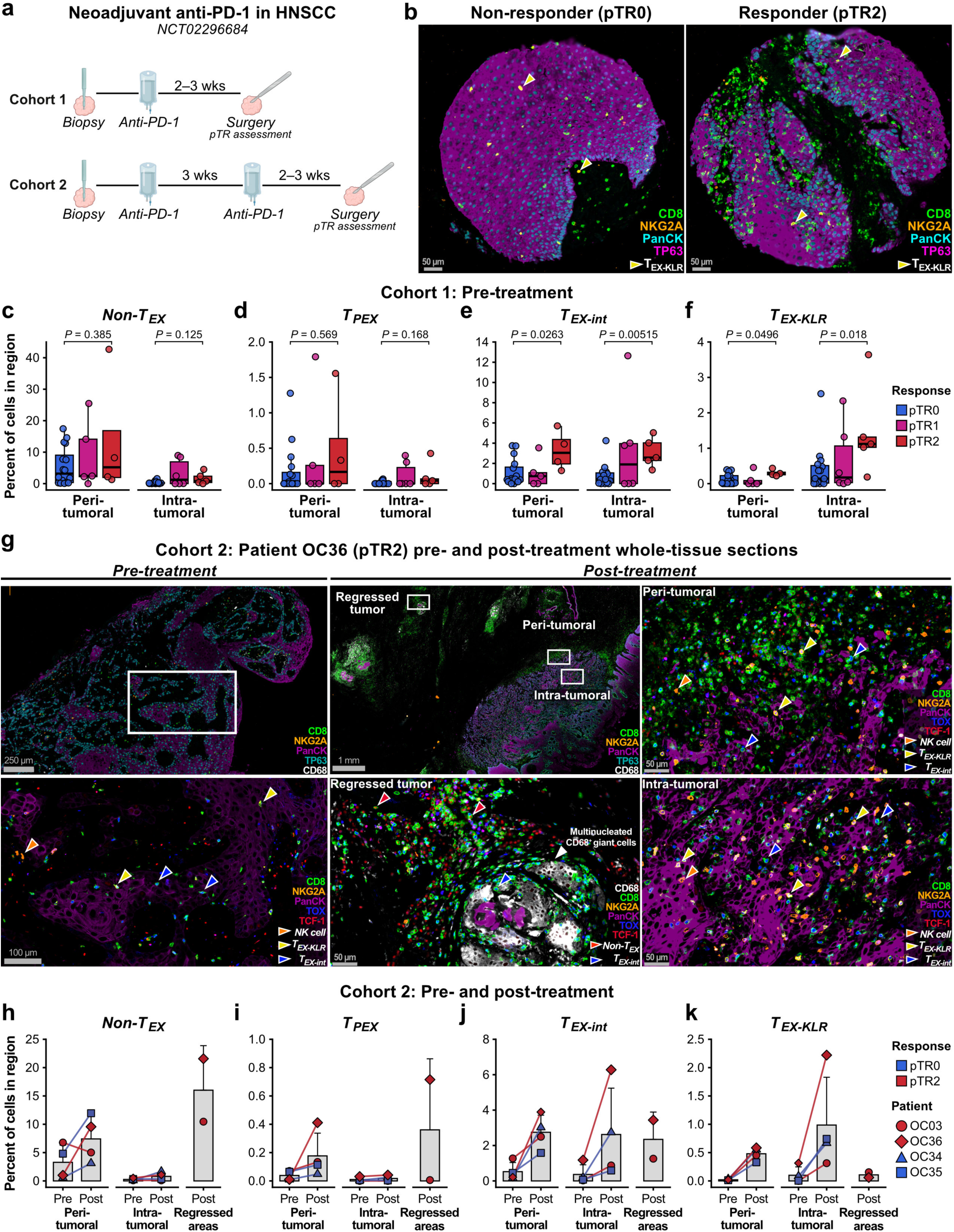
T_EX-int_ and T_EX-KLR_ cells are associated with anti-PD-1 response and increase in areas of viable tumor post-treatment. **a**, Schematic of the clinical trial design. **b**, Representative immunofluorescence images of the Cohort 1 pre-treatment tissue microarray samples of a non-responder (pTR0) and a responder (pTR2). **c–f**, Percentage of (**c**) non-T_EX_, (**d**) T_PEX_, (**e**) T_EX-int_, and (**f**) T_EX-KLR_ cells among all cells within the peri- and intra-tumoral regions in the pre-treatment Cohort 1 samples by response type. Unadjusted *P* values are shown for Wilcoxon rank-sum tests comparing the cell frequencies between the pTR0 and pTR2 samples in the peri-tumoral (*n* = 16 pTR0 and 4 pTR2) and intra-tumoral (*n* = 15 pTR0 and 5 pTR2) regions. **g**, Representative immunofluorescence images of pre- and post-treatment whole-tissue sections of Cohort 2 patient OC36 (pTR2). **h–k**, Percentage of (**h**) non-T_EX_, (**i**) T_PEX_, (**j**) T_EX-int_, and (**k**) T_EX-KLR_ cells among all cells within the peri-tumoral, intra-tumoral, and regression areas at the pre- and post-treatment time points in Cohort 2 samples (*n* = 2 pTR0 and 2 pTR2).

Before treatment in the single-dose Cohort 1, neither the non-T_EX_ nor the T_PEX_ cell frequency among all cells differed across the pTR0, pTR1, and pTR2 groups (**Fig. 6b–d**). By contrast, pTR2 samples had higher frequencies of T_EX-int_ and T_EX-KLR_ cells than pTR0 samples, both *peri*- and *intra*-tumorally (**Fig. 6e, f**). We did not observe a change in the frequency of any subset after anti-PD-1 in the Cohort 1 non-responder samples, and there were insufficient paired samples to analyze this in the responders. As an orthogonal analysis, we applied the avidity-associated T_EX-int_ and T_EX-KLR_ DEG signatures as module scores to T_EX_ TILs in a scRNA-seq dataset of triple-negative breast cancer patients who received neoadjuvant anti-PD-1 therapy (**Extended Data Fig. 11**)^35^. In that dataset, expression of the T_EX-KLR_ signature in T_EX_ TILs at both pre- and post-treatment time points was associated with response, which was assessed after anti-PD-1 followed by radiation and chemotherapy. Together, these data support that T_EX-int_ and T_EX-KLR_ cells are associated with response to neoadjuvant anti-PD-1 therapy.

To better resolve the spatial dynamics of the T_EX_ subsets during anti-PD-1 therapy, we analyzed multiplexed immunofluorescence images from the available whole-tissue sections of two non-responders (pTR0) and two responders (pTR2) in Cohort 2 (**Fig. 6g** and **Extended Data Fig. 12**). Although the limited sample size precluded statistical testing, the frequencies of non-T_EX_, T_PEX_, T_EX-int_, and T_EX-KLR_ cells were similar between response types, a result that differs from that of Cohort 1 and may be attributable to the additional cycle of anti-PD-1 administered in Cohort 2 (**Fig. 6h–k**). Additionally, all four paired tumor samples showed an increase in the *peri*-tumoral frequency of each T_EX_ subset following treatment (**Fig. 6i–k**). Within the *intra*-tumoral region, only T_EX-int_ and T_EX-KLR_ cells substantially increased post-treatment, which is consistent with a role of these cells in tumor elimination (**Fig. 6j, k**). However, in responders, the post-treatment *intra*-tumoral region represents residual viable cancer cells within a tumor that was overall responsive to anti-PD-1 therapy, as indicated by regressed tumor areas containing multinucleated CD68^+^ giant cells, keratinous debris, and a high lymphocyte density (**Fig. 6g** and **Extended Data Fig. 12**). In these regressed areas, non-T_EX_ cells were the majority, and T_EX-int_ and T_EX-KLR_ cells were depleted relative to the remaining *peri*- and *intra*-tumoral regions, suggesting that these T_EX_ cells localize to viable cancer cells during PD-1 blockade (**Fig. 6h, j, k** and **Extended Data Fig. 12**). This distribution of non-T_EX_, T_EX-int_, and T_EX-KLR_ cells is consistent with the higher proportion of non-T_EX_ cells among CD8^+^ TILs reported in the post-treatment scRNA-seq data of responders in this and other HNSCC cohorts^2,14^, which may reflect a relative loss of T_EX-int_ and T_EX-KLR_ cells in exchange for non-T_EX_ cells as viable tumor is eliminated. In sum, these results suggest that T_EX-int_ and T_EX-KLR_ cells accumulate near viable cancer cells after anti-PD-1 therapy, and that the post-treatment increase in T_EX-KLR_ cells may reflect stronger NFAT-dependent TCR signaling upon PD-1 blockade.

In conclusion, our data support that TCR avidity governs reproducible phenotypes of T_EX_ cells through NFAT, with high-avidity TCR signaling driving the T_EX-KLR_ state. The association between TCR avidity and human tumor-infiltrating T_EX_ states warrants further investigation to identify targetable pathways to improve T_EX_ function and cancer immunotherapies.

## DISCUSSION

T_EX_ cells are important to immunotherapy response in solid tumors, where transcriptionally distinct T_EX_ subsets dynamically change upon ICB therapy^2,9,14,15,36–42^. Here, we comprehensively profiled tumor-infiltrating T_EX_ subsets in HNSCC and defined T_PEX_, T_EX-int_, and T_EX-KLR_ as core T_EX_ populations. Additionally, in agreement with Fesneau *et al*.^24^, we found that among CD8 T cells, T_EX-KLR_ cells are enriched in HPV-HNSCC tumors relative to HPV+ tumors, which favor the T_PEX_ and T_EX-int_ subsets. Future work is needed to clarify the association between T_EX_ phenotypes and HPV status in HNSCC, which could be related to the source of tumor-associated antigens, either neoantigen- or HPV-derived. The T_PEX_, T_EX-int_, and T_EX-KLR_ subsets have also been observed in other cancer types and datasets, but cells transcriptionally resembling the T_EX-int_ or T_EX-KLR_ subset have been referred to by several names. For example, the T_EX-int_ subset has been labeled “GZMK^+^ T_EX_” or “cytotoxic T_EX_” based on its expression of *GZMK*, *GZMH*, and *EOMES*^2,16^, and the T_EX-KLR_ subset has been called “dysfunctional,” “terminal,” or “extreme exhausted” based on its elevated expression of CD39 (*ENTPD1*) and inhibitory receptors^2,11,14–16^. We propose the name T_EX-KLR_, as our data suggest that NKG2A (*KLRC1*) distinguishes T_EX_ cells with high functional avidity from their lower avidity counterparts, T_EX-int_ cells. A KLR^+^ T_EX_ subset has also been described in mice, but consistent with prior literature, we find that the murine KLR^+^ T_EX_ signature is most highly expressed in human non-T_EX_ NK-like CD8 T cells, which express *CX3CR1*, a key marker gene for the murine KLR^+^ T_EX_ subset (**Extended Data Fig. 13** and **Supplementary Table 9**)^43^. To capture human T_EX_ subsets in future studies, we propose use of antibodies against CD39, NKG2A, and CCR6 for flow cytometry and PD-1, TOX, NKG2A, and TCF-1 for immunofluorescence, in addition to the antibodies needed to mark CD8 T cells. When combined with Cellular Indexing of Transcriptomes and Epitopes by sequencing (CITE-seq) or spatial transcriptomics, these may improve alignment of transcriptional profiles with reproducible T_EX_ subsets.

In addition to nomenclature, hypotheses on the origin of T_EX-KLR_ cells have differed, with IL-12^24–26^ and TGF-β^44,45^ each proposed to drive the T_EX-KLR_ state because they upregulate NKG2A on activated CD8 T cells *in vitro*. However, we find that the broader *in vivo* T_EX-KLR_ gene program, along with NKG2A, is likely driven by a third mechanism: NFAT-dependent high-avidity TCR signaling. In further support of a TCR-intrinsic mechanism, T_EX-int_ and T_EX-KLR_ cells were near each other in tumors, suggesting that paracrine signals are unlikely to fully account for expression of NKG2A and the T_EX-KLR_ gene program. Together, our data suggest that understanding the determinants of TCR signal strength, such as catch-bond formation^46^, may be particularly insightful when studying the biology of exhaustion in human tumors and its applications to ICB and TCR-T cell therapies.

While our findings support that paracrine signals alone are unlikely to drive the flux of T_EX_ states, investigating how cytokines shape TCR signaling, and in turn T_EX_ phenotypes, could inform cytokine-based immunotherapies, such as IL-2^47,48^, IL-12^49,50^, and anti-TGF-β^51^. For example, we found that IL-12 antagonizes an NFAT-dependent high-avidity TCR signaling gene program while potentiating cytotoxicity, supporting that IL-12 or other mechanisms that modulate TCR signaling may enhance T_EX_ function. However, IL-12 therapies have been limited by systemic toxicity, although some novel therapies have focused on tumor-specific delivery^49,50^. Because unopposed NFAT signaling promotes CD8 T cell dysfunction^52–54^, identifying how IL-12 signaling suppresses high-avidity NFAT signaling may reveal alternative strategies to target this pathway with less toxicity; STAT4, NF-κB, AP-1, BATF3, and MAP3K8 are potential candidates^27,55–57^. Future research on the relationship between cytokines, TCR signal strength, and T_EX_ phenotypes may reveal additional pathways that could be leveraged to improve T_EX_ function.

Here, we defined NFAT signaling as a key driver of the T_EX-KLR_ gene program, but the functional role of these genes remains an open question. We propose that the T_EX-KLR_ state serves to restrain activation-induced cell death (AICD) and preserve high-avidity clonotypes, enforced by upregulation of inhibitory genes beyond *PDCD1*, such as *KLRC1*^58,59^, *SNX9*^60^, and *DUSP2*^61^. Understanding this state, therefore, carries important therapeutic implications. For example, ICB may precipitate AICD in T_EX-KLR_ cells, as shown in mice where anti-PD-1 induces apoptosis in high-avidity, stem-like T_EX_ clonotypes^62^. Anti-PD-1 may also shift low-avidity clonotypes into the T_EX-KLR_ state by increasing TCR signal strength, which may explain the increase in T_EX-KLR_ cells near viable cancer cells that we observed in HNSCC following anti-PD-1. Finally, anti-PD-1 in combination with additional ICB may be ineffective due to the breadth of inhibitory genes upregulated in the T_EX-KLR_ state. Collectively, these possible effects of ICB on T_EX-KLR_ cells may explain the failure of NKG2A blockade with monalizumab, in combination with either cetuximab (anti-EGFR) or durvalumab (anti-PD-L1), to improve survival in platinum- and anti-PD-1-refractory HNSCC^63,64^. As an alternative, adding supportive or stimulatory cytokines, such as IL-2 or IL-12 analogs^49,65,66^, to anti-PD-1, with or without anti-NKG2A, may better exploit the high-avidity TCR signaling of T_EX-KLR_ cells for immunotherapy. In summary, our work provides a mechanistic framework for the balance between T_EX-int_ and T_EX-KLR_ states in tumors, with important implications for improving the effector functions of T_EX_ cells across human cancers.

### Limitations

Our study has several potential limitations. First, the *in vitro* co-culture model using peripheral CD8 T cells that were stimulated, transduced with the E7 TCR, and then re-stimulated upon co-culture may not fully recapitulate the activation of T_EX_ cells *in vivo*. Second, NFAT inhibition was achieved by blocking calcineurin with FK506, and calcineurin inhibition may affect TCR signaling independent of NFAT. Third, the multiplexed immunofluorescence findings from the neoadjuvant immunotherapy HNSCC cohort are based on relatively small tissue samples in the tissue microarray or pre-treatment core biopsies. These smaller tissue samples may not represent the spatial heterogeneity of the whole tumor nor capture a sufficient area to analyze pre- to post-treatment changes of the same tumor. Additionally, the results are limited by modest patient numbers in the Cohort 1 tissue microarray. Although the whole-tissue section analyses on the paired Cohort 2 pre- and post-anti-PD-1 samples captured the diverse spatial patterns of cellular localization, these analyses were limited to a modest sample size of available tissue sections. The association of T_EX-KLR_ cells with immunotherapy response and residual viable tumor requires validation in larger cohorts. Lastly, there may be additional subtleties of the T_EX-KLR_ state and other T_EX_ subtypes among cancer types that we did not assess.

## RESOURCE AVAILABILITY

### Lead Contact

Further information and requests for resources should be directed to the lead contact, Sidharth V. Puram.

## Materials Availability

The plasmids and other resources used in this study are available upon reasonable request to the lead contact.

## Data and Code Availability

Raw and processed immunofluorescence, scRNA-seq, scTCR-seq, and bulk RNA-seq data will be made publicly available upon final publication. This paper does not report original code. The Xue *et al*., 2025 dataset was obtained from 10.5281/zenodo.13382785. The Ibáñez-Molero *et al*., 2026 dataset was downloaded from GSE283942. The Shiao *et al*., 2024 dataset was downloaded from GSE246613. The Daniel *et al*., 2022 dataset was downloaded from GSE188666. The Mints *et al*., 2026 dataset of treatment-naïve HNSCC samples was downloaded as raw FASTQ files from SRA: PRJNA1283925.

## AUTHOR CONTRIBUTIONS

R.D.Z.M., J.M.Z., T.F.B., and S.V.P. conceptualized the study. R.D.Z.M., J.M.Z., T.F.B., and S.V.P. wrote and edited the manuscript with input from all authors. R.D.Z.M., J.M.Z., T.F.B., E.S., P.B., and C.H. performed experiments. R.D.Z.M., J.M.Z., T.F.B., S.V.P., R.D.M., T.E., G.P., and N.S. guided experiments. R.D.Z.M. and J.M.Z. performed the analyses. R.D.Z.M. and J.M.Z. prepared the figures. R.U. and A.M.E. contributed samples from the neoadjuvant anti-PD-1 clinical trial in HNSCC and supported the immunofluorescence staining for the samples. The following authors were responsible for clinical management and collection of patient samples: C.B., A.B., S.R., A.R., J.L., K.S., D.M., J.D.L.S., A.J.A., N.R., W.T., P.O., B.J.K., C.A., J.T.R., R.S.J., P.P., P.A.Z., B.W., R.A.H., D.R.A., S.D.K., R.U., and S.V.P.

## ACKNOWLEDGMENTS AND FUNDING

We are grateful to the Genome Technology Access Center (GTAC) at the McDonnell Genome Institute for sequencing services. This work was supported by K08CA237732 (NIH, S.V.P.), R21DE031072 (NIH, S.V.P.), 1R01DE032371 (NIH, S.V.P.), 1R01DE032865 (NIH, S.V.P.), 5T32CA009547-38 (NIH, J.M.Z.), 5K12CA167540-15 (NIH, J.M.Z.), F31DE032562 (NIH, R.D.Z.M.), R01AI130152 (NIH, T.E.), R01AI176664 (NIH, T.E.), V Foundation (S.V.P.), Cancer Research Foundation (S.V.P.), Barnes-Jewish Hospital Foundation (S.V.P.), and Doris Duke Foundation (S.V.P.). The funding sources were not involved in the design, conduct, or reporting of the research. Schematics were designed with BioRender.

## SUPPLEMENTARY TABLES

Supplementary Table 1. Human sample information.

Supplementary Table 2. Cluster markers of the HNSCC TILs dataset.

Supplementary Table 3. Differential gene expression results for each TEX subset against the pooled non-TEX clusters and each TEX cluster against each other.

Supplementary Table 4. Tumor- and viral-reactive signatures obtained from Supplementary Table 4 of Ibáñez-Molero *et al*., 2026.

**Supplementary** Table 5. Bulk RNA-seq differential gene expression results.

**Supplementary** Table 6. k-means clusters of the bulk RNA-seq activation-induced genes.

**Supplementary** Table 7. Avidity association of the differentially expressed genes between the T_EX-KLR_ and T_EX-int_ subsets.

**Supplementary** Table 8. Avidity-dominant, IL-12-dominant, and shared bulk RNA-seq gene sets.

**Supplementary** Table 9. Murine T_EX_ subset signatures.

**Supplementary** Table 10. CODEX immunofluorescence staining antibodies and cell type assignment strategy.

Extended Data Figure 1. Classification of scRNA-seq clusters

**a**, UMAP plot of the HNSCC TILs dataset.

**b**, Annotation transfer from the huARdb CD8 T cell atlas (*left*; Xue *et al*., 2025) to the HNSCC TILs dataset (*right*); the published UMAP coordinates and clusters of the huARdb dataset are shown.

**c**, Proportions of the transferred annotations within each HNSCC TILs cluster.

**d**, Dot plot of top marker genes per HNSCC TILs cluster.

**e–h**, Per-patient median module scores in each scRNA-seq cluster for tumor-reactive signatures from (**e**) Lowery *et al*., 2022, (**f**) Meng *et al*., 2023, and (**g**) Oliveira *et al*., 2021, and (**h**) for a viral-reactive signature from Oliveira *et al*., 2021.

Extended Data Figure 2. Flow cytometry of HNSCC patient tumors

**a**, Flow cytometry gating strategy for the HNSCC patient tumor samples.

**b–c**, Percentage of (**b**) CD4 T cells and (**c**) CD8 T cells among all T cells in HPV+ and HPV-tumor samples in the flow cytometry data (*n* = 16 HPV+ and 15 HPV-tumors). Unadjusted *P* values from a Wilcoxon rank-sum test are shown.

Extended Data Figure 3. *Ex vivo* stimulation of sorted non-T_EX_, T_EX-int_, and T_EX-KLR_ cells

**a**, Flow cytometry gating strategy for sorting non-T_EX_, T_EX-int_, and T_EX-KLR_ cells from freshly dissociated HNSCC patient tumor samples.

**b**, Sorting purity assessed by analyzing the cells immediately post-sort.

**c–h**, Cells were rested or stimulated with immobilized anti-CD3 for four days *in vitro* and then analyzed by flow cytometry (*n* = 1 tumor sample for rest and 3 tumor samples for anti-CD3 stimulation).

**c–e**, Representative flow cytometry plots of (**c**) CD39 and KLRG1 expression, (**d**) NKG2A and CCR6 expression, and (**e**) CD137 expression and CFSE.

**f–h**, Percentage of (**f**) CD39^+^, (**g**) NKG2A^+^, and (**h**) CD137^+^ CFSE^-^ cells. *P* values from a repeated-measures one-way ANOVA with Tukey’s correction are shown for the anti-CD3 conditions.

Extended Data Figure 4. Phenotypes of expanded CD8 T cell clonotypes

**a**, Number of CD8^+^ TILs with or without a TCR captured by scTCR-seq per patient.

**b**, Number of expanded T_EX_ clonotypes dominant within each T_EX_ cluster per patient.

**c–d**, Cluster composition of up to 10 of the largest clonotypes per patient among (**c**) expanded T_EX_ clonotypes or (**d**) all CD8 T cells. Only patients with at least 600 cells with a TCR are shown.

Extended Data Figure 5. Immunofluorescence data phenotyping

**a–b**, Immunophenotyping marker proteins used in **Fig. 2** for (**a**) broad cell types and (**b**) T cell subsets; (**b**) is also shown as **Fig. 2b**.

**c–d**, Immunophenotyping marker proteins used in **Fig. 6** for (**c**) broad cell types and (**d**) T cell subsets. Total count of cells per type is indicated above as a labeled bar in (**a–d**).

Extended Data Figure 6. *In vitro* model of TCR avidity in the UM-SCC47 cell line

**a**, Flow cytometry gating strategy for analysis of E7 TCR^+^ CD8 T cells following *in vitro* co-culture.

**b**, Flow cytometry histograms of HLA-A2 expression on UM-SCC47 cell lines normalized to the mode (*left*) and quantified as mean fluorescence intensity (MFI; *right*).

**c**, IL-12p70 concentration in the media following culture of E7 TCR^+^ CD8 T cells in the indicated condition for 48 h (*n* = 4 donors; E7 TCR^+^ CD8 T cells generated from donors ID85, ID267, ID481, and ID948 were used).

Extended Data Figure 7. High-avidity TCR signaling promotes NKG2A expression in co-cultures with HEK293T target cells

**a**, Schematic of HEK293T target cell lines, including peptide pulsing and HPV16 E7 overexpression (OE; *left*) and the WT and mutant E7_11–20_ SCT cell lines (*right*).

**b–c**, Representative flow cytometry (**b**) histograms of CD3, CD137, and NKG2A expression normalized to the mode and (**c**) plots of NKG2A and CD137 expression on E7 TCR^+^ CD8 T cells following 48 h co-culture. Data for E7 TCR^+^ CD8 T cells generated from donor LRS071724 are shown.

**d–f**, Quantification of (**d**) CD3 and (**e**) CD137 mean fluorescence intensity (MFI) relative to E7 TCR^+^ CD8 T cells cultured alone and (**f**) percentage of NKG2A^+^ E7 TCR^+^ CD8 T cells (*n* = 3 donors; E7 TCR^+^ CD8 T cells generated from donors LRS072323, LRS050924, and LRS071724 were used). *P* values from a repeated-measures one-way ANOVA with Sidak’s correction are shown.

Extended Data Figure 8. NFAT and IL-12 independently regulate NKG2A expression and cytotoxicity

**a**, Representative flow cytometry plots of NKG2A and CD137 expression on E7 TCR^+^ CD8 T cells following 48 h culture in the indicated condition. Data for E7 TCR^+^ CD8 T cells generated from donor LRS081324 are shown.

**b**, Percentage of CD137^+^ (*left*) and NKG2A^+^ (*right*) E7 TCR^+^ CD8 T cells following rest or anti-CD3 and anti-CD28 stimulation for 48 h (*n* = 3 donors).

**c**, Percentage of UM-SCC47 target cells killed after 48 h co-culture with E7 TCR^+^ CD8 T cells (*n* = 7 replicates from 4 donors).

*P* values from a repeated-measures one-way ANOVA with Tukey’s correction for multiple comparisons are shown in (**b**, **c**).

Extended Data Figure 9. IL-12 signaling and high-avidity TCR signaling trigger shared and distinct gene expression programs

**a**, Differential gene expression of E7 TCR^+^ CD8 T cells co-cultured with UM-SCC47 target cells: HLA-A2 cells plus IL-12 (*y-axis*) or WT SCT cells (*x-axis*), each relative to co-culture with HLA-A2 cells plus vehicle as the baseline-activation reference. Gene data points are colored by their k-means cluster from the **Fig. 4d** heatmap. Diagonal lines indicate a 0.75 absolute log_2_FC difference between the contrasts, and colored overlays indicate the gene sets (see **Methods**).

**b**, Z-scored mean RNA expression of the genes in the IL-12-dominant, avidity-dominant, and shared gene sets for the cells of each patient per cluster. Genes are k-means clustered and ordered by the difference in the mean z-score of the T_EX-KLR_ columns and the T_PEX_ and T_EX-int_ columns.

**c–e**, Per-patient median module scores for the (**c**) avidity-dominant, (**d**) IL-12-dominant, and (**e**) shared gene sets in each scRNA-seq cluster (*n* = 21 T_PEX_, 28 T_EX-int_, and 23 T_EX-KLR_ patients). *P* values from a Wilcoxon rank-sum test with Benjamini–Hochberg correction are shown.

Extended Data Figure 10. Expanded T_EX_ clonotypes in HNSCC occupy a reciprocal axis between T_EX-int_ and T_EX-KLR_ states

**a**, UMAP plot of the HNSCC TILs dataset.

**b**, Module scores for the avidity-associated T_EX-int_ (*left*) and T_EX-KLR_ (*middle*) DEG signatures (from **Fig. 4g**) and the relative module scores represented as the Δz-score (*right*; Δz-score = z-scored T_EX-KLR_ module score minus z-scored T_EX-int_ module score).

**c**, Per-patient mean module scores in each scRNA-seq cluster for the avidity-associated T_EX-int_ (*top*) and T_EX-KLR_ (*middle*) DEG signatures and the Δz-score (*bottom*).

**d**, Relative avidity-associated DEG signature module scores in expanded T_EX_ clonotypes, where cells of each clonotype are arranged into columns and filled by their Δz-score, and clonotypes are ordered by their mean Δz-score. Cells of highlighted clonotypes are shown on a scatter plot. The cluster proportions, size, and dominant T_EX_ cluster of each clonotype are shown below.

Extended Data Figure 11. The avidity-associated T_EX-KLR_ DEG signature is associated with response to anti-PD-1 in triple-negative breast cancer

**a**, Schematic of the clinical trial design.

**b–d**, Per-patient mean module scores in T_EX_ cells for the avidity-associated (**b**) T_EX-KLR_ and (**c**) T_EX-int_ DEG signatures (from **Fig. 4g**) and (**d**) the relative module scores represented as the Δz-score (Δz-score = z-scored T_EX-KLR_ module score minus z-scored T_EX-int_ module score; *n* = 5 non-responders and 15 responders pre-treatment and *n* = 7 non-responders and 17 responders post-cycle 1). *P* values from a Wilcoxon rank-sum test with Benjamini–Hochberg correction are shown.

Extended Data Figure 12. Whole-tissue section immunofluorescence images of Cohort 2 patient OC03 (pTR2) before and after neoadjuvant anti-PD-1 therapy

**a**, Selected region from the pre-treatment whole-tissue section.

**b,** Post-treatment whole-tissue section. The selected peri-tumoral, intra-tumoral, and regression areas indicated by white boxes are shown below.

**c–h,** Views of the selected post-treatment (**c–d**) peri-tumoral, (**e–f**) intra-tumoral, and (**g–h**) regression areas showing T_EX_ subset marker proteins (*left*; **c, e, g**) and single-channel images (*right*; **d, f, h**). Selected cell types are indicated by colored arrows.

Extended Data Figure 13. Human tumor-infiltrating T_EX_ subsets are distinct from murine T_EX_ subsets in the lymphocytic choriomeningitis virus clone 13 chronic infection model

**a**, UMAP plot of the HNSCC TILs dataset with the T_EX-KLR_ and the non-T_EX_ NK-like T cell clusters outlined.

**b**, UMAP plot of murine lymphocytic choriomeningitis virus (LCMV) clone 13-specific CD8 T cells at day 21 post-infection from the Daniel *et al*., 2022 dataset. The published UMAP coordinates and clusters are shown, and the murine KLR^+^ T_EX_ cluster is outlined.

**c–d**, Expression of selected murine T cell subset markers in (**c**) HNSCC TILs and (**d**) murine LCMV clone 13-specific CD8 T cells.

**e**, Mean z-scored module scores for the murine T_EX_ subset signatures in the HNSCC TILs CD8 T cell clusters (see **Methods**).

**f**, Per-cell module scores for the murine T_EX_ subset signatures in the HNSCC TILs dataset.

## METHODS

### Human samples and tissue processing

#### Human samples

This study included new scRNA-seq and immunofluorescence data of treatment-naïve HNSCC tumor samples that were collected in accordance with the Declaration of Helsinki and approved by the IRB of Washington University (IRB# 201102323 and IRB# 202106015). The tissue samples from pre- and post-anti-PD-1-treated HNSCC samples shown in **Fig. 6** originated from the phase 2 clinical trial of neoadjuvant pembrolizumab for patients with HPV-stage III to IVb HNSCC, NCT02296684 (IRB# 201412118), with results previously reported^2^. Information for new human samples in this work is detailed in **Supplementary Table 1**.

#### Peripheral blood samples

Human peripheral blood was obtained from either leukocyte reduction system cones from the Mississippi Valley Regional Blood Center or from buffy coats obtained from the Gulf Coast Regional Blood Center, both of which are classified as non-human research.

#### Human tumor dissociation

Fresh HNSCC tumor samples were minced and washed with phosphate buffered saline (PBS; Sigma-Aldrich, P3813). Minced tumor was dissociated with the Human Tumor Dissociation Kit (Miltenyi Biotec, 130-095-929) according to the manufacturer’s protocol in a gentleMACS C-tube (Miltenyi Biotec, 130-093-237) using the gentleMACS Octo Dissociator with Heaters (Miltenyi Biotec) and the 37C_h_TDK_1 program. Dissociated cells were filtered through a 40-μm strainer (Greiner Bio-One, 542140), pelleted, and resuspended in 1 mL of ammonium-chloride-potassium (ACK) lysis buffer (Gibco, A1049201) for 30–60 s. Lysis was stopped by the addition of 10 mL autoMACS Rinsing Solution (Miltenyi Biotec, 130-091-222) supplemented with 0.5% w/v bovine serum albumin (BSA) from the MACS BSA Stock Solution (Miltenyi Biotec, 130-091-376). Cells not used fresh were cryopreserved in fetal bovine serum (FBS; Peak Serum, PS-FB1) supplemented with 10% dimethyl sulfoxide (DMSO; Sigma-Aldrich, D2650) at −80°C using the Mr. Frosty freezing container (Thermo Scientific, 15-350-50) and then transferred to liquid nitrogen for long-term storage.

### Materials and cell lines for the *in vitro* model of variable TCR avidity

#### Cell line culture

All cells were cultured at 37°C in a humidified chamber with 5% CO₂. UM-SCC47 cells were cultured in Ham’s F-12 Nutrient Mix (Gibco, 11765054) supplemented with 25% high-glucose Dulbecco’s modified Eagle’s medium (DMEM, Gibco, 11965118), 10% FBS (Peak Serum, PS-FB1), and 1% penicillin-streptomycin (P/S, Corning, 30-002-CI). Cells were passaged using Trypsin-EDTA (0.25%) (Gibco, 25200056) upon reaching 80% confluency.

HEK293T cells (Lenti-X HEK293T, Takara, 632180) were cultured in DMEM (Gibco, 11965118) supplemented with 10% FBS (Peak Serum, PS-FB1), 1% P/S (Corning, 30-002-CI), 1 mM sodium pyruvate (Gibco, 11360070), 1× minimal essential medium (MEM) non-essential amino acids (Gibco, 11140050), 1% GlutaMAX (Gibco, 35050061), and 10 mM HEPES (Gibco, 15630080). Cells were passaged every two days upon reaching approximately 70–80% confluence using TrypLE Express Enzyme (Gibco, 12604021) and reseeded at 2 × 10^6^ cells mL^−1^ per tissue-culture treated 15 cm dish.

#### CD8 T cell isolation from peripheral blood

Peripheral blood mononuclear cells (PBMCs) were isolated with Ficoll-Paque density gradient medium (MilliporeSigma, GE17-1440-02) using SepMate-50 tubes (StemCell Technologies, 85460). CD8 T cells were isolated using positive selection with CD8 MicroBeads (Miltenyi Biotec, 130-045-201) for leukocyte reduction system cones or EasySep Human CD8 Positive Selection Kit II (StemCell Technologies, 17853) for the buffy coats. Isolated cells were cryopreserved at a concentration of 1 × 10^7^ cells mL^−1^ in FBS (Peak Serum, PS-FB1) supplemented with 10% DMSO (Sigma-Aldrich, D2650) at −80°C using the Mr. Frosty freezing container (Thermo Scientific, 15-350-50) and then transferred to liquid nitrogen for long-term storage.

#### Plasmid construction

The CloneAmp HiFi PCR Premix (Takara, 639298) and primers obtained from Integrated DNA Technologies were used for PCR. The PCR products were purified using the NucleoSpin Gel and PCR Clean-Up kit (Takara, 740609) and cloned using the In-Fusion Snap Assembly master mix (Takara, 638948). Plasmids were cultivated in Stellar competent cells (Takara, 636763) and isolated using the QIAprep Spin Miniprep kit (QIAGEN, 27104) or the EndoFree Plasmid Maxi kit (QIAGEN, 12362). Full plasmid sequencing was performed using nanopore sequencing through Plasmidsaurus.

The TCR targeting HPV16 E7 was obtained as a gifted plasmid from Christian Hinrichs (Addgene plasmid # 122728; http://n2t.net/addgene:122728; RRID: Addgene_122728) and was subcloned into the N103 lentiviral expression vector. The human elongation factor 1 alpha (EF-1α) promoter was used to express the TCR, and the phosphoglycerate kinase (PGK) promoter was used to express puromycin N-acetyltransferase, which confers puromycin resistance.

The HLA-A*02:01 E7_11–20_ SCT constructs were cloned into the N103 backbone under regulation of the EF-1α promoter, and the PGK promoter was used to express puromycin N-acetyltransferase to confer puromycin resistance. The SCT constructs consisted of, from the N-to C-terminal, the β2 microglobulin (*B2M*) signal peptide, WT or mutant E7_11–20_ peptide nucleotide sequence, a G_4_S_3_ linker, exons 2–4 of *B2M*, a G_4_S_3_ linker, and lastly, the HLA-A*02:01 gene (GenBank AY365426.1). A gene block of the *B2M* signal peptide through the first G_4_S_3_ linker was ordered from Genewiz. Exons 2–4 of *B2M* and the HLA-A*02:01 gene were cloned from cDNA generated from the peripheral blood of a human donor. Amino acid substitutions were introduced in the E7_11–20_ peptide sequence with inverse PCR. The psPAX2 and pMD2.G lentiviral packaging plasmids were used to generate lentivirus. psPAX2 was a gift from Didier Trono (Addgene plasmid #12260; http://n2t.net/addgene:12260; RRID:Addgene_12260). pMD2.G was a gift from Didier Trono (Addgene plasmid #12259; http://n2t.net/addgene:12259 ; RRID:Addgene_12259).

The HPV16 E7 gene was obtained from the pB-actin E6 E7 plasmid as a gift from Karl Munger (Addgene plasmid #13712; http://n2t.net/addgene:13712; RRID: Addgene_13712). The HPV16 E7 gene was subcloned into the N103 lentiviral expression vector under control of the human cytomegalovirus immediate-early (CMV) promoter. The E7 gene was followed by an internal ribosomal entry sequence (IRES) and pMAX GFP with two N-terminal SV40 nuclear localization signals. The PGK promoter was used to express puromycin N-acetyltransferase in this plasmid.

#### Lentivirus production

Lenti-X HEK293T cells (Takara, 632180) were thawed and passaged twice before usage. The cells were then seeded at 3.5 × 10^6^ cells in 5 mL per T25 flask in complete Opti-MEM prepared by combining Opti-MEM (Gibco, 31985070) with 5% FBS (Peak Serum, PS-FB1), 1 mM sodium pyruvate (Gibco, 11360070), 1× MEM non-essential amino acids (Gibco, 11140050), and 1% GlutaMAX (Gibco, 35050061). The following morning, 2.5 mL of media was removed from the Lenti-X HEK293T culture. The cells were then transfected with Lipofectamine 3000 (Thermo Fisher Scientific, L3000008). In one 1.5 mL microcentrifuge tube, 625 μL of Opti-MEM and 20.11 μL of Lipofectamine 3000 reagent were mixed. In a second 1.5 mL microcentrifuge tube, 625 μL of Opti-MEM was mixed with 3.11 μg psPAX2, 1.56 μg pMD2.G, 4.22 μg of the transfer plasmid, and 18.56 μL of the P3000 reagent. The contents of the two tubes were then combined and pipet mixed. After 15 min incubation at room temperature, the transfection mix was added to the Lenti-X HEK293T culture. After 6 h, the media was refreshed with 5 mL complete Opti-MEM containing 1× ViralBoost (ALSTEM, VB100). Lentivirus was collected at 24 h and 48 h following transfection by collecting the culture media and centrifuging it at 500 × g for 5 min. The supernatant was then transferred into a new tube, and one-third of that volume of Lenti-X Concentrator (Takara Bio, 631231) was added. The tube was inverted to mix and incubated overnight at 4°C. After overnight incubation, the tube was centrifuged at 1,500 × g for 45 min at 4°C. The pellet was resuspended in Opti-MEM (Gibco, 31985070) without supplements at one hundredth of the original collected volume. Concentrated lentivirus was immediately snap-frozen in liquid nitrogen and stored at −80°C. The lentivirus was thawed at room temperature immediately before use.

#### Thawing non-transduced and transduced CD8 T cells

Cryopreserved CD8 T cells were thawed in pre-warmed RPMI 1640 (Gibco, 11875119) supplemented with 10% FBS (Peak Serum, PS-FB1), 1% P/S (Corning, 30-002-CI), 55 μM 2-mercaptoethanol (2-ME, Gibco, 21985023), 10 mM HEPES (Gibco, 15630080), 4 mM N-Acetyl-L-cysteine (NAC, Sigma-Aldrich, A9165), and 500 U mL^−1^ IL-2 (StemCell Technologies, 78036). T cells were cultured for 24 h in that media prior to use .

#### Lentiviral transduction of CD8 T cells

After thawing and resting for 24 h in the thawing media, CD8 T cells were collected and activated on plates coated with 5 μg mL^−1^ anti-CD3 (BioLegend, clone OKT3, 317348) for 2 h at 37°C at a density of 1 × 10^6^ cells mL^−1^ in ImmunoCult-XF T cell Expansion medium (StemCell Technologies, 10981) supplemented with 2 μg mL^−1^ anti-CD28 (BioLegend, clone CD28.2, 302943), 1% P/S (Corning, 30-002-CI), 55 μM 2-ME (Gibco, 21985023), and 500 U mL^−1^ IL-2 (StemCell Technologies, 78036). One day after activation, CD8 T cells were transduced with concentrated E7 TCR lentivirus (5% v/v). Two days after activation, puromycin (2.5 μg mL^−1^, PR1MA, KCP33020) was added to the culture media. Three days following activation, one-half volume fresh media was added to the existing media. The CD8 T cells were split two days later and cultured at 0.5 × 10^6^ cells mL^−1^ in media containing 0.25 μg mL^−1^ puromycin and without anti-CD28. The media was refreshed every three days thereafter to maintain a concentration of 0.5 × 10^6^ cells mL^−1^. On day 11 following activation, cells were cryopreserved at a density of 1 × 10^6^ cells mL^−1^ in FBS (Peak Serum, PS-FB1) supplemented with 10% DMSO (Sigma-Aldrich, D2650) at −80°C using the Mr. Frosty freezing container (Thermo Scientific, 15-350-50) and then transferred to liquid nitrogen for long-term storage.

#### Lentiviral transduction of UM-SCC47 and HEK293T cell lines

UM-SCC47 and Lenti-X HEK293T cells (Takara, 632180) were seeded at 50% confluency and transduced with 50% (v/v) of unconcentrated lentivirus. Two days after transduction, 2.5 μg mL^−1^ puromycin (PR1MA, KCP33020) was added to the culture media for three days. Cell lines were maintained in 1 μg mL^−1^ puromycin thereafter. These cell lines were also transduced with a nuclear-localized TdTomato expression vector under blasticidin selection (5 μg mL^−1^). UM-SCC47 cells transduced with the HLA-A*02:01 gene or SCTs were stained with anti-HLA-A2 APC (1:100, clone BB7.2, BioLegend, 343307) to quantify surface expression.

### *In vitro* co-culture experiments

#### Co-culture conditions for flow cytometry of CD8 T cells

For flow cytometry experiments, UM-SCC47 cells or Lenti-X HEK293T cells (Takara, 632180) were seeded at 3–4 × 10^4^ cells per well of a flat-bottom tissue-culture treated 96-well plate in RPMI 1640 (Gibco, 11875119) supplemented with 10% FBS (Peak Serum, PS-FB1) and 1% P/S (Corning, 30-002-CI). After 24 h, existing media was aspirated from each well and E7 TCR^+^ CD8 T cells were added at a 1:1 (target cells:E7 TCR^+^ CD8 T cells) ratio in 250 μL RPMI 1640 (Gibco, 11875119) supplemented with 10% FBS (Peak Serum, PS-FB1), 1% P/S (Corning, 30-002-CI), 300 U mL^−1^ IL-2 (StemCell Technologies, 78036), and any other treatments. Half of the volume was replaced with fresh media one day after adding the CD8 T cells, and the CD8 T cells were collected after 48 h of co-culture. For anti-CD3 and anti-CD28 activation, CD8 T cells were plated onto wells pre-coated with 5 μg mL^−1^ anti-CD3 (BioLegend, clone OKT3, 317348) and 2 μg mL^–1^ anti-CD28 (BioLegend, clone CD28.2, 302943) for 2 h at 37°C.

FK506 (Tacrolimus; Selleck Chemicals, S5003) was added at 5 nM and IL-12p70 (Peprotech, 200-12H) was added at 10 ng mL^−1^ to the co-culture at the same time that the E7 TCR^+^ CD8 T cells were added. FK506 was reconstituted to 10 μM in 100% ethanol (Sigma-Aldrich, E7023), and IL-12p70 was reconstituted to 10 μg mL^–1^ in distilled water supplemented with 0.2% BSA (w/v; Gold Biotechnology, A-420-100). The vehicle control for FK506 was 0.05% ethanol, and that of IL-12p70 was 0.0002% BSA.

#### Peptide pulsing

The HPV16 E7_11–20_ peptide (YMLDLQPETT) was obtained from JPT Peptide Technologies (SP-MHCI-0073-2) and reconstituted in DMSO (Sigma-Aldrich, D2650) at 0.2 mg mL^−1^. The peptide was added to the culture media at a concentration of 500 nM at the same time that E7 TCR^+^ CD8 T cells were added to the co-culture.

#### Flow cytometry compensation

For all flow cytometry experiments, antibody compensation beads were prepared with UltraComp eBeads (Invitrogen, 01-2222-41) stained with the same concentration of each antibody as used for the samples at 4°C for 30 min. Zombie NIR viability dye compensation was performed using ArC amine reactive compensation beads (Invitrogen, A10346) stained at 1:100 for 30 min at room temperature. Compensation beads were washed once in FACS buffer. Flow cytometry results were analyzed using FlowJo Software (v10.10.0, BD Life Sciences).

### Flow cytometry of E7 TCR^+^ CD8 T cells following *in vitro* co-culture

Following 48 h of co-culture, CD8 T cells were collected by washing the wells with PBS (Sigma-Aldrich, P3813). The CD8 T cells were then stained with Zombie NIR viability dye (1:2000, BioLegend, 423105) for 15 min at room temperature. The cells were then washed in FACS buffer, which consisted of PBS supplemented with 2% FBS (Peak Serum, PS-FB1). Subsequently, cells were stained with antibodies in 100 μL FACS buffer for 30 min at room temperature followed by one wash in FACS buffer, and data were immediately acquired. For the experiments of **Fig. 3** and **Extended Data Fig. 7**, data were acquired using the Cytoflex SRT instrument (Beckman Coulter) and CytExpert acquisition software (Beckman Coulter, v1.2.0.10004), and the following antibodies were used: anti-CD8 BV510 (1:50, clone SK1, BioLegend, 344732), anti-mouse TCR β chain BV605 (1:50, clone H57-597, BioLegend, 109241), anti-CD3 BV785 (1:50, clone UCHT1, BioLegend, 300472), anti-NKG2A PE (1:100, clone S19004C, BioLegend, 375104), and anti-CD137 APC (1:100, clone 4B4-1, BioLegend, 309809). For the experiments testing FK506 and IL-12p70, data were acquired using the LSRFortessa X-20 (BD Life Sciences) and FACSDiva software (BD Biosciences, v9.0), and the following antibodies were used: anti-CD8 BV510 (1:100, clone SK1, BioLegend, 344732), anti-mouse TCR β chain BV605 (1:50, clone H57-597, BioLegend, 109241), anti-CD3 BV785 (1:100, clone UCHT1, BioLegend, 300472), anti-NKG2A PE (1:100, clone S19004C, BioLegend, 375104), and anti-CD137 APC (1:100, clone 4B4-1, BioLegend, 309809).

#### IL-12p70 enzyme-linked immunosorbent assay (ELISA)

Cells were cultured under conditions identical to those listed under the methods section, “*Co-culture conditions for flow cytometry of CD8 T cells*.” Following 48 h of co-culture, media was collected, centrifuged at 1,000 × g for 10 min, and snap-frozen in liquid nitrogen. Supernatants were thawed on ice and diluted 1:5 prior to the IL-12p70 ELISA using the human IL-12p70 ELISA kit (MilliporeSigma, RAB0252-1KT) according to the manufacturer’s protocol. Absorbance was measured on the Agilent BioTek Cytation 5 Cell Imaging Multi-Mode Reader (Agilent, CYT5FVSN).

#### T cell cytotoxicity assay

Cancer cell lines expressing nuclear localized TdTomato were plated at 2 × 10^4^ cells per well of a 96-well plate in RPMI 1640 (Gibco, 11875119) supplemented with 10% FBS (Peak Serum, PS-FB1) and 1% P/S (Corning, 30-002-CI). After 24 h, the media was aspirated from the cancer cells, and 2 × 10^4^ E7 TCR^+^ T cells were added in 200 μL fresh media containing 300 U mL^–1^ IL-2 (StemCell Technologies, 78036). Following 24 h of co-culture, the wells were washed twice with FACS buffer supplemented with 1 mM ethylenediaminetetraacetic acid (EDTA; Thermo Scientific, 15575020) and then 50 μL of FACS buffer was added to the well for imaging. The RFP channel was used to image each well on the Agilent BioTek Cytation 5 Cell Imaging Multi-Mode Reader (Agilent, CYT5FVSN). Target cell killing was quantified as the TdTomato^+^ nuclei in each well analyzed within the BioTek Gen5 software (v3.17). For each experiment, wells with UM-SCC47 cells only without co-culture with T cells were included to determine the percentage of target cells killed as defined by the following equation:

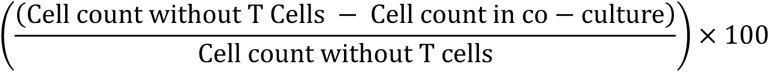

For visualization only, differences in the absolute magnitudes of killing across conditions in separate experiments were normalized: percent killing values were mean-centered by subtracting the experiment-level grand mean across all donors, cell lines, and treatments followed by adding back the grand mean calculated on both experiments. Statistics were performed on the raw percent killing values.

### Flow cytometry of human tumor samples and *ex vivo* stimulation of sorted T_EX_ subsets

#### Flow cytometry of human tumor samples

Samples were thawed and washed three times in magnesium- and calcium-free PBS (Corning, 21-040-CM) supplemented with 0.5% w/v BSA (Gold Biotechnology, A-420-100) and 200 U mL^−1^ DNase I (MilliporeSigma, 10104159001). Cells were incubated with Zombie NIR viability dye (1:4000, BioLegend, 423105) and Human TruStain FcX (1:20, BioLegend, 422301) in PBS (Sigma-Aldrich, P3813) for 15 min at room temperature. Cells were then washed in FACS buffer and stained in 100 μL FACS buffer for 30 min at room temperature with anti-CD39 BV421 (1:200, clone A1, BioLegend, 328214), anti-CD8 BV510 (1:200, clone SK1, BioLegend, 344732), anti-CD45 BV605 (1:200, clone HI30, BioLegend, 304042), anti-CD14 BV650 (1:100, clone M5E2, BioLegend, 301836), anti-CD3 BV785 (1:200, clone UCHT1, BioLegend, 300472), anti-CD137 KIRAVIA Blue 520 (1:40, clone S18014C, BioLegend, 300816), anti-NKG2A PE (1:100, clone S19004C, BioLegend, 375104), anti-CD16 PE/Dazzle 594 (1:100, clone 3G8, BioLegend, 302054), anti-CD4 PE-Cy7 (1:1000, clone SK3, BD Biosciences, 557852), anti-CCR6 APC (1:40, clone G034E3, BioLegend, 353416), and anti-KLRG1 BD Horizon R718 (1:40, clone Z7-205.rMAb, BD Biosciences, 568659). Cells were washed once in FACS buffer and data were acquired immediately using the Cytoflex SRT instrument (Beckman Coulter) and CytExpert acquisition software (Beckman Coulter, v1.2.0.10004).

#### CD8 T cell sorting from tumor samples for *ex vivo* stimulation

To reduce debris prior to flow cytometry sorting, CD8^+^ cells or CD45^+^ cells were positively selected from freshly dissociated HNSCC tumor samples using the REAlease CD8 (TIL) MicroBead Kit (Miltenyi Biotec, 130-121-560) or REAlease CD45 (TIL) MicroBead Kit, human (Miltenyi Biotec, 130-121-563), respectively. One sample was not pre-sorted prior to flow cytometry sorting (see **Supplementary Table 1**). The tumor cell suspension or positively selected cells were washed and stained in PBS (Sigma-Aldrich, P3813) with Zombie NIR viability dye (1:4000, BioLegend, 423105) for 10 min at room temperature. Cells were then washed in FACS buffer and stained in 100 μL FACS buffer for 20 min at room temperature with Human TruStain FcX (1:10, BioLegend, 422301), anti-CD39 BV421 (1:200, clone A1, BioLegend, 328214), anti-CD8 BV510 (1:200, clone SK1, BioLegend, 344732), anti-CD45 BV605 (1:200, clone HI30, BioLegend, 304042), anti-CCR6 BV650 (1:40, clone G034E3, BioLegend, 353425), anti-CD3 BV785 (1:200, clone UCHT1, BioLegend, 300472), anti-CD14 PerCP (1:50, clone HCD14, BioLegend, 325602), anti-CD19 PerCP (1:50, clone HIB19, BioLegend, 302227), anti-NKG2A PE (1:100, clone S19004C, BioLegend, 375104), anti-PD-1 PE/Dazzle 594 (1:50, clone EH12.2H7, BioLegend, 329939), anti-TIM-3 PE/Cy7 (1:50, clone F38-2E2, BioLegend, 345013), anti-CD137 APC (1:100, clone 4B4-1, BioLegend, 309809), and anti-KLRG1 BD Horizon R718 (1:40, clone Z7-205.rMAb, BD Biosciences, 568659). CD14-PerCP and CD19-PerCP were used to exclude monocytes (CD14^+^) and B cells (CD19^+^) by marking both with PerCP.

Cells were washed with FACS buffer and then immediately sorted using the Cytoflex SRT instrument (Beckman Coulter) and CytExpert acquisition software (Beckman Coulter, v1.2.0.10004). Cells were sorted into 1.5 mL tubes containing 500 μL of TIL media: RPMI 1640 (Gibco, 11875119) supplemented with 10% FBS (Peak Serum, PS-FB1), 1% P/S (Corning, 30-002-CI), 1 mM sodium pyruvate (Gibco, 11360070), 1× GlutaMAX (Gibco, 35050061), 1× MEM non-essential amino acids (Gibco, 11140050), 50 μM 2-ME (Fisher Scientific, AC125470100), 0.25 μg mL^−1^ amphotericin B (Fisher Scientific, BP264520), and 300 U mL^−1^ IL-2 (StemCell Technologies, 78036).

### *Ex vivo* stimulation of FACS-sorted CD8^+^ TIL subsets from human HNSCC tumors

After sorting, cells were centrifuged and resuspended in 100 μL PBS (Sigma-Aldrich, P3813) followed by the addition of 100 μL of a 3 μM solution of CellTrace CFSE in PBS (Invitrogen, C34554). Cells were incubated for 4 min at room temperature followed by the addition of 1.3 mL of TIL media. Cells were then centrifuged, resuspended in TIL media, and plated in flat-bottom 96-well plates at 1 × 10^5^ cells per well (or all cells if fewer were recovered) in 100 μL TIL media. After four days of culture, cells were collected and processed for flow cytometry using the same panel and protocol as described for cell sorting.

### Multiplexed immunofluorescence

#### Cyclic multiplexed immunofluorescence using Co-Detection by Indexing (CODEX)

All processing steps were performed in the WashU Immunomonitoring Laboratory (IML) Core. Freshly sectioned human formalin-fixed paraffin-embedded (FFPE) tissue slides were baked for 90 min at 65°C in a HybEZ II Oven (Advanced Cell Diagnostics, USA) immediately prior to dewaxing. Deparaffinization and rehydration were performed on the Parhelia Spatial Station platform with their proprietary dewaxing agent. Antigen retrieval was performed at 108°C for 30 min in 1× Tris-EDTA pH 9.0 (Invitrogen, 00-4956-58). The staining chamber was cooled to 30°C followed by rinsing in distilled water twice and then transferring into the Akoya Hydration Buffer for 2 min. The chamber was then washed again with Hydration buffer. Samples were stained in the Parhelia Spatial Station by first applying the Akoya Staining Buffer without antibodies for 20 min and then incubating in 120 μL of the antibody mix at room temperature in a humidified chamber for 3 h. Slides were then washed and fixed as detailed in the Akoya PhenoCycler-Fusion User Guide_1.0.3_RevD protocol. The slides were photobleached for 45 min in PBS supplemented with 20 mM NaOH and 4.5% (v/v) hydrogen peroxide while positioned between two LED lights. After photobleaching, the flow cells were applied and the slides were imaged on the Akoya PhenoCycler Fusion 2.0 CODEX imaging system (Akoya Biosciences, USA). Images were collected as 8-bit QPTIFF files with embedded metadata. Non-Akoya antibodies were ordered carrier-free from their respective manufacturers and conjugated to the indicated Phenocycler DNA barcodes according to manufacturer instructions. Antibodies and dilutions used are listed in **Supplementary Table 10**.

#### CODEX image tissue and tumor boundary classifications

Tissue boundary objects were created by pixel thresholder using QuPath v0.6.0-rc349 on the DAPI channel (16.23 μm/px resolution, smoothing 2, threshold 5, minimum object size 1000 μm^2^, and minimum hole size 5000 μm^2^). Staining or tissue artifacts (i.e., tissue folds, bubbles, large reporter aggregates, cautery edge artifacts) were annotated manually and excluded from segmentation and further analysis. A high-resolution tumor mask was similarly created within the tissue annotation by a pixel classifier on the pan-cytokeratin (PanCK) channel (2.03 μm/px, smoothing 2, threshold 20, minimum object size 500 μm^2^, and minimum hole size 500 μm^2^). Areas of normal squamous epithelium and secretory glands were manually annotated and subtracted from the mask. Areas of pathologic regression were identified as keratinous debris (PanCK^+^, TP63^-^, DAPI^-^) surrounded by CD68^+^ multi-nucleated giant cells. These areas were manually subtracted from the tumor mask as a separate annotation. The intra-tumoral region was defined as the area within the tumor mask, and the peri-tumoral region was defined as the 200 μm of stroma extending outward from the tumor–stroma interface.

#### CODEX cell segmentation

Nuclear and cell segmentation were performed with the InstanSeg plugin within QuPath, applying the associated pre-trained model (fluorescence_nuclei_and_cells, CPDMI_2023 dataset). The DAPI, PanCK, TP63, CD3E, CD4, CD8, CD45RO, CD14, CD20, and HLA-A channels were used for cell segmentation. Signed minimum distance to the tumor and regression annotation masks for each cell were calculated within QuPath using the ‘Analyze,’ then ‘Spatial Analysis,’ and then ‘Signed distance to annotation 2D’ functions. For each cell, mean pixel fluorescent intensity per marker, a unique cell ID, centroid x and y pixel coordinates, and distance metrics were exported for further analysis in Python (v3.12.3) with Scanpy (v1.10.4).

#### Cell phenotyping of CODEX imaging data

For each image, fluorescent intensity was normalized for each marker by taking the centered log ratio (CLR) of each cell’s mean marker intensity over the geometric mean of all cells in the image, as implemented with pseudocounts in the scverse muon (v0.1) package. CLR is similar to the log transformation and z-score normalization per sample suggested by Hickey *et al*., 2021, which uses the mean of all cells to account for variation in background staining and preserves the true 0 values. A CLR value of 1.1 represents a ∼2-fold difference in intensity over the mean of all cells in the tissue. Histograms of marker CLR values per sample were assessed for marker quality control and outliers. Otsu thresholds for each marker were calculated to guide threshold selection. For cell type assignment, we applied a hierarchical flow cytometry-like hand-gating, according to the schema in **Supplementary Table 10**, which was applied uniformly to all images. Cell classification was applied in two stages, first for broad cell type annotation of T cells (CD8^+^ and CD4^+^), NK cells, B cells, macrophages, and tumor cells. Cells initially typed as CD4 T cells, CD8 T cells, and NK cells were then passed for secondary detailed phenotyping. Cell type classifications were examined for accuracy by visual inspection within QuPath for each image.

#### Cell type frequency analysis of the immunofluorescence data

For each tumor sample, the percentage of each cell subset among the specified denominator was calculated separately within the peri- and intra-tumoral regions. Any region with fewer than 50 nucleated cells or CD8 T cells, depending on which was the denominator in the analysis, was excluded.

#### Odds ratio of intra-tumoral versus peri-tumoral presence in the immunofluorescence data

For each patient and cell type, a two-by-two contingency table was created of region (intra-tumoral versus peri-tumoral) against cell class (the count of the cell type of interest versus all other nucleated cells). Per-patient log odds ratios and their variances were obtained with the Table2x2 and combine_effects functions of statsmodels (v0.14). These were pooled across patients by random-effects meta-analysis using the DerSimonian–Laird estimator of tau-squared and Knapp–Hartung standard errors, which produced the pooled odds ratio and 95% confidence interval per cell type. A pooled odds ratio significantly different from 1 was assessed by a two-tailed Wald test, and *P* values were corrected across all cell types by the Benjamini–Hochberg method.

#### Neighborhood analysis of the immunofluorescence data

The neighbor analysis was performed with the scimap (v2.2.11) package using the spatial_count function with the neighborhood radius set to 20 μm (40 pixels, for images at 0.5 μm per pixel), limiting the analysis to cells within the peri-tumoral and intra-tumoral regions. For each sample, the mean number of each other cell type per index cell type was plotted.

### Bulk RNA-seq

#### Bulk RNA-seq of CD8 T cells following co-culture with UM-SCC47 cell lines

UM-SCC47 cells were seeded at 3 × 10^5^ cells per well of a 24-well plate in RPMI 1640 (Gibco, 11875119) supplemented with 10% FBS (Peak Serum, PS-FB1) and 1% P/S (Corning, 30-002-CI). 24 h after seeding the cancer cell lines, the existing media was removed and 3 × 10^5^ E7 TCR^+^ CD8 T cells were added at a 1:1 ratio in 2 mL of RPMI 1640 (Gibco, 11875119) supplemented with 300 U mL^−1^ IL-2 (StemCell Technologies, 78036), 10% FBS (Peak Serum, PS-FB1), 1% P/S (Corning, 30-002-CI), and the specified treatments. Because non-activated CD8 T cells have low levels of RNA, 6 × 10^5^ E7 TCR^+^ CD8 T cells were added to the following target cell co-culture conditions to obtain sufficient RNA for library preparation: empty vector, HLA-A2 plus FK506, and HLA-A2 plus FK506 and IL-12p70. After 24 h from adding the CD8 T cells, 1 mL of media was removed and 1.4 mL of fresh media was added. The CD8 T cells were collected after 48 h of co-culture by washing the wells with PBS (Sigma-Aldrich, P3813) and then purifying the viable CD8 T cells using Ficoll-Paque density gradient medium (MilliporeSigma, GE17-1440-02) and SepMate-50 tubes (StemCell Technologies, 85460). The CD8 T cells were lysed in DNA/RNA Shield (Zymo Research, R1100) and stored at −80°C until submitting to Plasmidsaurus for bulk RNA-seq.

#### Bulk RNA-seq data processing

Bulk RNA-seq data pre-processing was performed by Plasmidsaurus. FASTQ generation and demultiplexing were performed using BCL Convert (v4.3.6) and fqtk (v0.3.1). Reads were filtered using FastP (v0.24.0) with poly-X tail trimming, 3’ quality-based tail trimming, Phred quality score ≥ 15, and minimum length of 50 bp. Reads were aligned to hg38 with STAR (v2.7.11), removing non-canonical splice junctions and unmapped reads. PCR and optical duplicates were removed using UMICollapse (v1.1.0). Gene expression was quantified with the subread package (v2.1.1) using strand-specific counting, fractional assignment of multi-mapping reads, and exons and three-prime UTRs as feature identifiers.

Additional processing of the gene count matrix was performed to retain only protein-coding genes, lncRNA, mitochondrial rRNA and tRNA, rRNA, and TCR constant- and variable-region genes. TCR variable-region genes were collapsed to the locus level (TRAV, TRBV, TRGV, TRDV) by summing counts across all variable-region genes sharing a locus. Fractional counts arising from multi-mapping reads were rounded to integers. Count matrices from two sequencing runs (LZHV7Z and TVNKCT) were merged on the intersection of gene symbols.

The merged count matrix was input to DESeq2 (v1.46.0) with patient and condition in a paired design (design = ∼Donor + Condition). Genes with at least 10 counts in at least two samples were retained (14,015 genes). Variance stabilizing transformed counts (VST, blind = FALSE) with patient-level batch effects removed by the removeBatchEffect function in limma (v3.62.2) were used for visualization and clustering. Differential gene expression analyses were performed using the DESeq2 Wald test with Benjamini–Hochberg *P* value correction within each contrast.

#### Heatmap of activation-induced genes in the bulk RNA-seq data

Activation-induced genes were defined as those upregulated in either the HLA-A2 UM-SCC47 target cell plus IL-12p70 or the WT SCT UM-SCC47 target cell co-culture conditions, each versus the empty vector control UM-SCC47 cells (log_2_FC > 1, adjusted *P* < 0.01, *n* = 1,777 genes). The batch-corrected and variance-stabilized expression matrix of these genes was z-scored per gene across samples with values capped at ±4. Fill color was capped at ±2 for visualization. k-means clustering (k = 5, 3 random starts) was applied to the z-scored matrix, and within each cluster, genes were sorted by the log_2_FC of HLA-A2 plus IL-12p70 versus WT SCT target cell co-culture conditions. One k-means cluster of 60 genes was enriched for contaminating cancer-cell genes and therefore excluded from downstream analyses, retaining four k-means clusters with a total of 1,717 genes that were plotted as a heatmap with ComplexHeatmap (v2.22.0).

#### Definition of IL-12-dominant, avidity-dominant, and shared gene sets

DEGs of E7 TCR^+^ CD8 T cells co-cultured with HLA-A2 UM-SCC47 target cells plus IL-12p70 or with WT SCT UM-SCC47 target cells, each relative to HLA-A2 plus vehicle target cells, were assessed by DESeq2 (adjusted *P* < 0.05 in at least one of the two contrasts). The difference in the log_2_FC between contrasts (Δlog_2_FC) for each gene was also calculated. The analysis was restricted to the 1,717 activation-induced genes shown in the **Fig. 4d** heatmap, and of those, genes were classified into sets according to the following criteria: *avidity-dominant* (log_2_FC > 0.5 in the WT SCT contrast, log_2_FC < 0.5 in the IL-12 contrast, and |Δlog_2_FC| > 0.75; *n* = 151 genes), *IL-12-dominant* (log_2_FC < 0.5 in the WT SCT contrast, log_2_FC > 0.5 in the IL-12 contrast, and |Δlog_2_FC| > 0.75; *n* = 117 genes), or *shared* (log_2_FC > 0.5 in both contrasts, irrespective of the Δlog_2_FC; *n* = 164 genes).

### Single-cell sequencing data generation and sample processing

#### Isolation of CD8^+^ TILs from HNSCC samples for scRNA-seq

Treatment-naïve samples for P5202, P5055, P5026, P5022, P5144, P5193, P5125, P5172, P5145, P5173, and P5169 were obtained from PRJNA1283925^19^. The scRNA-seq data of the HNSCC TILs from tumor samples spanning WashU01 to WashU20 originated from freshly dissociated tumor samples. CD8^+^ TILs were isolated with the REAlease CD8 (TIL) MicroBead Kit (Miltenyi Biotec, 130-121-560) according to the manufacturer’s protocol. A CD45 sort was then performed on the CD8^-^ TILs with the REAlease CD45 (TIL) MicroBead Kit (Miltenyi Biotec, 130-121-563) according to the manufacturer’s protocol. Separate libraries were prepared for the CD8^+^ sorted fraction (‘cd8p’) and a mixture of 70% CD8^-^ CD45^-^ and 30% CD8^-^ CD45^+^ cells (denoted ‘cd8n’). The CD3^+^ CD8^-^ NK cells and the CD8 T cells present in the CD8^-^ library (‘cd8n’) were included in the analyzed HNSCC TILs scRNA-seq dataset of this study.

The remaining scRNA-seq data of CD8^+^ TILs from tumor samples with the “WashU” prefix were obtained from cryopreserved tumor cell suspensions sorted by flow cytometry. Samples were thawed and washed three times in magnesium- and calcium-free PBS (Corning, 21-040-CM) supplemented with 0.5% w/v BSA (Gold Biotechnology, A-420-100) and 200 U mL^−1^ DNase I (MilliporeSigma, 10104159001). Cells were stained with Zombie UV viability dye (1:40, BioLegend, 423107) in PBS (Sigma-Aldrich, P3813) for 10 min at 4°C. After washing in FACS buffer, cells were then resuspended in FACS buffer containing Human TruStain FcX (1:20, BioLegend, 422301). The following antibodies were then added: anti-CD4 PE-Cy7 (1:50, clone SK3, BioLegend, 344611), anti-CD8 BV510 (1:50, clone SK1, BioLegend, 344731), anti-CD3 BV785 (1:50, clone UCHT1, BioLegend, 300471), and anti-CD45 BUV737 (1:100, clone HI30, BD Biosciences, 568524). Cells were incubated at 4°C for 20 min and washed in PBS supplemented with 2% FBS. Viable (Zombie UV^-^), CD45^+^, CD3^+^, CD8^+^, CD4^-^ cells were sorted on the Cytoflex SRT instrument (Beckman Coulter) using CytExpert acquisition software (Beckman Coulter, v1.2.0.10004). For both modes of isolation, the sorted CD8 T cells were resuspended to a concentration of 800–1,200 cells μL^-1^ and confirmed to have ≥ 90% viability based on exclusion of 0.2% trypan blue (Thermo Fisher Scientific, 15250061).

#### scRNA-seq library preparation and sequencing

Single cell suspensions were processed with the Chromium Next GEM Single Cell 5’ Kit reagents (10x Genomics, 1000265) and Chromium Next GEM Chip K Single Cell Kit (10x Genomics, 1000286) according to the manufacturer’s protocol. Cells were loaded onto the chip to form gel bead-in-emulsions, targeting a recovery of 10,000 cells. Reverse transcription was performed using the resulting emulsion followed by purification and amplification of cDNA. To prepare scRNA-seq gene expression libraries, amplified cDNA was tagmented followed by adapter ligation and PCR amplification using the Library Construction Kit (10x Genomics, 1000190). TCR libraries were prepared with the Chromium Single Cell Human TCR Amplification Kit (10x Genomics, 1000252). scRNA-seq gene expression libraries were sequenced to a target depth of 50,000 paired-end reads per cell, and TCR libraries were sequenced to a target depth of 5,000 paired-end reads per cell. Both library types were sequenced on Illumina NovaSeq instruments at the Genome Technology Access Center (GTAC) at the McDonnell Genome Institute at Washington University School of Medicine.

#### Processing of the Mints *et al*., 2026 pre-treatment HNSCC samples

The FASTQ data for the treatment-naïve samples (available at SRA: PRJNA1283925)^19^ for P5202, P5055, P5026, P5022, P5144, P5193, P5125, P5172, P5145, P5173, and P5169 were aligned with cellranger-6.0.0 count and vdj commands to the joint GRCh38 with high-risk HPV reference and vdj_GRCh38_alts_ensembl-7.1.0 reference respectively.

#### Processing of the HNSCC TILs scRNA-seq dataset of this study

Raw sequencing data were processed with Cell Ranger multi (10x Genomics, v7.0.0) using the GRCh38 reference with high-risk HPV genomes HPV16, 18, 31, 33, and 25 concatenated onto the end as extra chromosomes^67^, and the vdj_GRCh38_alts_ensembl-7.1.0 reference for T cell receptor assembly. The filtered feature-barcode matrices were loaded into Seurat (v5.4.0) in R (v4.4.1). For all newly sequenced samples of this study and the Mints, *et al.*, 2026 samples, which together make up the HNSCC TILs scRNA-seq dataset, cells were filtered to retain those with > 500 detected genes and < 15% mitochondrial reads. Ambient RNA contamination was estimated and corrected with SoupX (v1.6.2). Doublets were identified per sample and removed if identified by 2 of 3 programs with default parameters: scDblFinder (v1.18.0), scds (v1.20.0), and scran (v1.32.0). Following pre-processing, samples were combined into a single Seurat object and counts were log-normalized with the NormalizeData function. TCR genes (TR[ABGD][CDJV]) were removed to avoid clustering cells by TCR usage. Genes with counts in fewer than 300 cells were removed. The top 3,000 variable features were identified with the FindVariableFeatures function using the “vst” method. SketchData was applied with “LeverageScore” method using 80,000 cells. Normalized counts were scaled across the cells of each patient using the ScaleData function. PCA was performed with RunPCA, and the first 50 principal components were used to create the nearest-neighbor graph, perform Louvain clustering (resolution = 1), and compute the UMAP. The PCA embeddings, clusters, and UMAP were then projected to the full dataset with ProjectData. A module score of canonical cell type markers was applied^19^. Cells in “T/NK” annotated clusters were retained and re-processed through the same method. Clusters dominated by CD8 T cells or NK cells were retained, and cells in clusters enriched for both CD8 T cells and CD4 T cells (i.e., naïve and cycling clusters) were filtered for expression of *CD8A* and *CD8B* > 0 and *CD4* = 0. The remaining cells were re-processed through the same method but without sketch/projection. The 26 clusters at “RNA_snn_res.1” were manually merged and annotated into fourteen populations based on canonical marker gene expression and the transferred annotations from the scAtlasVAE processing.

#### Processing of scTCR-seq data

The scTCR-seq filtered_contig_annotations.csv files from Cell Ranger were processed using scirpy (v0.23.0), scanpy (v1.10.4), and mudata (v0.1.6) in Python (v3.12). For each sample, filtered gene expression matrices and VDJ data were loaded with the scanpy read_10x_h5 and scirpy.io.read_10x_vdj functions respectively. The TCR alpha and beta chains were indexed and quality-controlled with the scirpy.pp.index_chains and scirpy.tl.chain_qc functions. The clonotypes were then defined per patient (pre- and post-time points together, if applicable) using scirpy.tl.define_clonotypes with receptor_arms = “any”, dual_ir = “all”, and within_group equal to the patient identifier. Clonotype size was defined as the number of cells sharing an identical clonotype identifier per patient.

### Analysis of single-cell data

#### Assignment of HNSCC TILs expanded T_EX_ clonotypes to a dominant T_EX_ cluster

Expanded T_EX_ clonotypes, defined as ≥ 15 cells in the T_EX_ clusters and ≥ 70% of the clonotype’s cells within the T_EX_ clusters, were assigned a dominant T_EX_ cluster if at least 50% of the clonotype’s T_EX_ cells were within a single T_EX_ cluster. Otherwise, the T_EX_ clonotype was labeled as mixed.

#### Quantification of gene sets as module scores in bulk RNA-seq and scRNA-seq datasets

Gene sets were applied as module scores by inputting normalized counts (log1p counts per 10,000 for scRNA-seq data and log_2_ counts per million for bulk RNA-seq data) into the AddModuleScore function of Seurat (v5.4.0): genes were grouped into 24 bins based on average expression across all cells, and for every gene in the gene set, 100 control genes were drawn at random from the same expression bin without replacement. The module score for each cell, or each sample for the bulk RNA-seq application, was calculated as its mean expression of the genes in the gene set minus the mean expression of the control genes. For only the Xue *et al*., 2025 huARdb scRNA-seq dataset, the module score was applied only to the exhausted subclusters with the score_genes function of scanpy (v1.10.4) using 25 bins of 50 control genes each.

#### Identification of top marker genes across the HNSCC TILs scRNA-seq clusters

The top marker genes across all clusters were identified using the Seurat (v5.4.0) Wilcoxon rank-sum test (FindAllMarkers function, positive markers only) followed by filtering for gene detection in at least 50% of the cells in either the cluster or the remaining cells of all other clusters (min.pct = 0.5), an average log_2_FC > 1, and a Bonferroni method-adjusted *P* < 0.01. Up to five of the top marker genes per cluster, ranked by the product of the average log_2_FC and the difference in the percentage of cells expressing the gene, were plotted in **Extended Data Fig. 1d**, and genes among the top five in more than one cluster were assigned to the first cluster in the plotting order. A separate FindAllMarkers analysis (positive markers only, min.pct = 0.10, average log_2_FC > 0.25, Wilcoxon rank-sum test) was performed to create **Supplementary Table 2**.

#### Identification of DEGs between non-T_EX_ and T_EX_ clusters of the HNSCC TILs scRNA-seq dataset

Pseudo-bulk profiles were generated by summing the raw counts of patient-group combinations from three T_EX_ scRNA-seq clusters – T_PEX_, T_EX-int_, and T_EX-KLR_ – and a single pooled non-T_EX_ group encompassing the T_N_, T_CM_/KLRG1^-^ T_EM_, T_EM_, acute activation T_EM_, HSP^+^ T_EM_, ITGAE^+^ T_RM_, MAIT, and NK-like T cell clusters. The CD56^+^ CD8^-^ NK cell, ISG^+^, and cycling clusters were excluded from the analysis. Differential gene expression was tested using the DESeq2 Wald test with the paired design of ∼Patient + Cluster; patient-cluster combinations with fewer than 50 cells were excluded, and only patients contributing to both groups were retained in each contrast. Only genes detected in at least 10% of the cells in at least one group of each contrast were tested for differential expression; the detection filter for the pooled non-T_EX_ group was the cell-number-weighted mean of the fraction of cells in each of its constituent clusters.

The DEGs (adjusted *P* < 1 × 10^−5^ and |log_2_FC| > 1) were used to define five signatures (**Supplementary Table 3**): *(1)* the common T_EX_ signature – genes upregulated against the non-T_EX_ group in each of the three T_EX_ subsets, excluding any gene differential in all three pairwise T_EX_ comparisons; *(2–4)* the three T_EX_ subset signatures – genes upregulated in one T_EX_ subset against each of the other T_EX_ subsets; and *(5)* the non-T_EX_ signature – genes upregulated in the non-T_EX_ group against each of the three T_EX_ subsets, excluding any gene in a T_EX_ subset signature. Ten genes met the criteria for both the common T_EX_ signature and a T_EX_ subset signature (9 T_EX-KLR_ genes, 1 T_PEX_ gene). These were assigned to the common T_EX_ signature because although they were relatively higher in one T_EX_ subset compared with the other two T_EX_ subsets, they were nonetheless upregulated in all T_EX_ subsets relative to non-T_EX_ cells and therefore better described as common T_EX_ marker genes. For display in **Fig. 1b**, the pseudo-bulk counts of all genes tested were variance-stabilized (DESeq2 vst, ∼Patient + Cluster, blind = FALSE) and z-scored per gene across 79 displayed pseudo-bulk samples of patients present in at least two of the four groups. In total, 233 genes were displayed (*n* = 87 non-T_EX_, 79 common T_EX_, 16 T_PEX_, 16 T_EX-int_, and 35 T_EX-KLR_ genes), and within each signature, genes were arranged in descending order by their smallest log_2_FC across the contrasts involved in the signature. The z-scored expression was plotted with the ComplexHeatmap package (v2.22.0) with the fill color clipped at ±1.5.

#### Analysis of tumor- and viral-reactive signatures in the HNSCC TILs scRNA-seq dataset

Tumor- and viral-reactive signatures were obtained from Supplementary Table 4 of Ibáñez-Molero *et al*., 2026, which contains a list of genes for the tumor- and viral-reactive signatures evaluated in this study. These signatures were applied as module scores to the HNSCC TILs scRNA-seq dataset, with the CD56^+^ CD8^-^ NK cell cluster removed because it was not a CD8 T cell subset. The z-scored means of the per-patient median module scores in each cluster, retaining only patients with at least 20 cells in the cluster, were plotted with the ComplexHeatmap package (v2.22.0), and fill color was capped at ±1.5. The median module scores for each patient with at least 20 cells in the cluster were also visualized with a boxplot.

#### Determination of avidity-associated T_EX-int_ versus T_EX-KLR_ DEG signatures

DEGs were determined between the T_EX-int_ and T_EX-KLR_ subsets using pseudo-bulk profiles, which were generated by summing the raw counts of each patient-cluster combination with at least 50 cells in both clusters, retaining 17 patients and 34 pseudo-bulk profiles. Genes detected in at least 10% of the retained patients’ cells in at least one of the two T_EX_ subsets were retained for analysis (*n* = 6,294 of 18,429 total detected genes). Differential expression was tested using the DESeq2 Wald test with a paired design of ∼Patient + Cluster. Genes with an adjusted *P* < 1 × 10^−5^ and |log_2_FC| > 1 were considered significant. 120 of the 129 genes upregulated in the T_EX-KLR_ subset and 46 of the 50 genes upregulated in the T_EX-int_ subset were present in the filtered bulk RNA-seq gene count matrix and applied as module scores across the bulk RNA-seq samples. The percentage of DEGs associated with avidity is reported using all DEGs as the denominator, not only the genes present among the filtered bulk RNA-seq gene count matrix.

For each DEG, the variance-stabilized expression (DESeq2 vst, blind = FALSE) was regressed on the corresponding module score across the 16 samples of the four avidity-axis conditions (co-culture with the UM-SCC47 empty vector, HLA-A2, Q6E SCT, or WT SCT target cells) using the limma package (v3.62.2) with donor as a fixed covariate and empirical Bayes moderation. Genes with a Benjamini–Hochberg-adjusted *P* < 0.05 within their own signature and a slope > 0.1 against their own module score were defined as avidity-associated (*n* = 57 of 120 T_EX-KLR_ DEGs and 23 of 46 T_EX-int_ DEGs). For displaying expression of these genes in **Fig. 4g**, donor effects were removed with the removeBatchEffect function in limma (v3.62.2), and each gene was z-scored across all 28 samples of all seven co-culture conditions. The genes were ordered by descending slope within each signature, and the fill color was capped at ±1.5.

Because each gene contributes to the module score on which it was regressed, which may introduce false positives, the analysis was repeated with two different methods (**Supplementary Table 7**). First, we performed a leave-one-out analysis by recomputing each module score with one gene excluded followed by recomputing the slope and *P* value of the excluded gene; Spearman ρ was greater than 0.998 between the original and leave-one-out slopes across genes. Second, the avidity-axis conditions were assigned an ordinal rank against which the genes were regressed, a method that is independent of the module score yet imposes an equal separation among conditions that does not strictly reflect the relative avidity. The rank was oriented from empty vector to WT SCT, with conditions assigned 1 through 4. The slopes for the genes in the T_EX-int_ module score were multiplied by –1 such that a positive slope indicated a positive association among both T_EX-int_ and T_EX-KLR_ genes. At adjusted *P* < 0.05 and slope > 0.1, 52 of the 57 T_EX-KLR_ genes (*GRAMD2B*, *KIFC3*, *LINC00299*, *LINC02539*, and *PLPP1* were absent) and 20 of the 23 T_EX-int_ genes (*DNAJB1*, *DRAIC*, and *SERTAD3* were absent) passed the filters. Of the absent genes, all except *SERTAD3* met the adjusted *P* value filter but did not reach the slope threshold; SERTAD3 met neither.

#### Analysis of the IL-12-dominant, avidity-dominant, and shared bulk RNA-seq gene sets in the HNSCC TILs scRNA-seq dataset

The IL-12-dominant, avidity-dominant, and shared gene sets were applied as module scores to the HNSCC TILs scRNA-seq dataset, retaining only genes expressed in at least 5% of cells in at least one CD8 T cell cluster (**Supplementary Table 8**; *n* = 59 IL-12-dominant, 105 avidity-dominant, and 104 shared genes). The CD56^+^ CD8^-^ NK cell cluster was removed prior to applying the module score because it was not a CD8 T cell subset. Additionally, the average log-normalized expression of these genes among the cells of each patient-cluster combination with at least 25 cells was z-scored and k-means clustered (k = 3, 100 random starts) and plotted as a heatmap with the ComplexHeatmap package (v2.22.0); only the T_PEX_, T_EX-int_, T_EX-KLR_, and cycling clusters were used for k-means clustering. The k-means clusters and their constituent genes were plotted in decreasing order of the mean of the z-scored T_EX-KLR_ columns minus the mean of the z-scored T_EX-int_ and T_PEX_ columns, and fill color was capped at ±1.5 for visualization. The median module score of each patient-cluster combination with at least 20 cells was used in the box plots and statistical comparisons between clusters.

#### Analysis of the relative avidity-associated T_EX-int_ and T_EX-KLR_ DEG signatures in expanded T_EX_ clonotypes

For the huARdb CD8 T cell atlas analysis, the object was subset to only cells present in the T_EX_ clusters of the huARdb metadata, and the “ag_spcific” cell subset was removed because these cells included adoptively transferred CD8^+^ TILs. The avidity-associated T_EX-int_ and T_EX-KLR_ DEG signatures were then applied as module scores to all T_EX_ cells. The data were then subset to expanded T_EX_ clonotypes, defined as ≥ 15 T_EX_ cells of the same clonotype (*n* = 58,166 cells from 772 clonotypes), and the module scores were z-scored across these cells. The difference in the z-scored module scores, defined as the z-scored T_EX-KLR_ module score minus the z-scored T_EX-int_ module score, was considered the relative expression of these signatures (Δz-score = z[T_EX-KLR_ module score] – z[T_EX-int_ module score]). For the analysis of the HNSCC TILs dataset, expanded T_EX_ clonotypes were defined as those with at least 15 cells in the T_EX_ clusters and at least 70% of the clonotype’s cells within the T_EX_ clusters (*n* = 24,844 cells from 325 clonotypes). The avidity-associated T_EX-int_ and T_EX-KLR_ DEG signatures were applied as module scores to the cells of those clonotypes, and the resulting scores were used to calculate the Δz-score. For both datasets, expanded T_EX_ clonotypes were plotted in ascending order of the mean Δz-score of the clonotype from left to right, with all cells of a clonotype stacked vertically and filled by their Δz-score. The fill color was capped at ±5 and the y-axis at –5 to 7.5 for visualization. Module scores were also computed on the whole HNSCC TILs dataset and displayed as feature plots and box plots of the per-patient mean scores within each cluster among patient-cluster combinations with at least 20 cells. For the feature plots of module scores, the color scale was capped at the 0.5^th^ and 99.5^th^ percentiles for the huARdb atlas dataset and the 0.1^st^ and 99.9^th^ percentiles for the HNSCC TILs dataset, each rounded to the nearest 0.05. Cells were ordered by descending module score or absolute Δz-score, with the highest cells on top.

### Processing and analysis of public single-cell datasets

#### Processing and analysis of the Xue *et al*., 2025 huARdb CD8 T cell scRNA-seq atlas

The huARdb scRNA-seq atlas raw gene counts matrix, metadata, UMAP coordinates, and cluster assignments were downloaded from Zenodo (10.5281/zenodo.13382785). For the T_EX_ clonotype analysis in **Fig. 5**, only solid tumor CD8^+^ TILs were retained, and the “ag_spcific” cell subset was removed because these cells included adoptively transferred CD8^+^ TILs. The raw counts were normalized to 10,000 counts per cell and log1p-transformed. Otherwise, all data and metadata, including UMAP coordinates and cluster assignments and names, were used as provided without further processing. Clonotypes were defined as cells of the same donor with identical TCR alpha and beta chain CDR3 amino acid sequences and identical TCR V and J genes for both chains.

#### Projection of the Xue *et al*., 2025 huARdb CD8 T cell scRNA-seq atlas cell type annotations onto scRNA-seq datasets

Annotations and UMAP coordinates were transferred from the huARdb v2 CD8 atlas (Xue *et al*., 2025) onto other scRNA-seq datasets using the scAtlasVAE package (v1.0.5.9; https://github.com/WanluLiuLab/scAtlasVAE) in Python (v3.8.20). First, raw counts in AnnData format were subset to the 4,000 highly variable genes of the reference (huARdb_v2_GEX.CD8.hvg4k.h5ad). Missing genes were filled with zeroes using the scatlasvae.pp._preprocess.subset_adata_by_genes_fill_zeros function. Cell type label transfer was performed with the scatlasvae.pipeline.run_transfer function to assign the high-level subtype label (cell_subtype_3). Cells annotated within a T_EX_ cluster were additionally projected onto a T_EX_-specific reference model (huARdb_v2_GEX.CD8.hvg4k.Tex.supervised.model) using the scatlasvae.pipeline.run_transfer function. Query cells were placed in the CD8 T cell atlas UMAP coordinate space using the scatlasvae.tl.umap_alignment function with method = “knn” using the huARdb_v2_GEX.CD8.hvg4k.X_gex.npy reference latent representation. All random seeds were fixed at 42 in this analysis. The “S100A11+ TEX” cluster name in the huARdb atlas was renamed to “XBP1+ TEX,” which is the same population, to be consistent with the Xue *et al*., 2025 manuscript, and the authors’ published cluster identities and UMAP coordinates were used.

#### Processing and analysis of the Ibáñez-Molero *et al*., 2026 scRNA-seq dataset

The scRNA-seq dataset from Ibáñez-Molero *et al*., 2026 was downloaded from GSE283942. The authors’ published cluster identities and UMAP coordinates were used. The TCR chains were indexed and checked for quality with the scirpy.pp.index_chains and scirpy.tl.chain_qc functions, and the filtered 10x VDJ contig annotations were then integrated using scirpy (v0.23.0) using receptor_arms = “any”, dual_ir = “primary_only”, and within_group set to the patient identifier. Only cells in the published metadata were retained. Module scores for the avidity-associated T_EX-KLR_ and T_EX-int_ DEG signatures were calculated on all cells using Seurat (v5.4.0) AddModuleScore. Cells were subset to those with a clonotype in one of the three FACS-sorted subsets – T cell singlets, cancer cell doublets, and APC doublets – and then the module scores were z-scored; the difference in the z-scored module scores, defined as the z-scored T_EX-KLR_ module score minus the z-scored T_EX-int_ module score, was considered the relative expression of these signatures (Δz-score = z[T_EX-KLR_ module score] – z[T_EX-int_ module score]). Clonotypes with at least 3 cells in at least 2 of the FACS-sorted subsets were retained for analysis, together with their cells from only the subsets containing at least 3 cells of that clonotype (12,710 cells from 312 clonotypes). For the per-clonotype analysis, the mean Δz-score was calculated for the cells of each clonotype within each FACS-sorted subset. For the feature plots, expression was scaled between the 0.1^st^ and 99.9^th^ percentiles within each feature, and cells were ordered by descending expression with the highest on top.

#### Processing and analysis of the Shiao *et al*., 2024 scRNA-seq dataset

The scRNA-seq dataset from Shiao *et al*., 2024 was downloaded from GSE246613. The authors’ published cluster identities and UMAP coordinates were used. The downloaded object was filtered to the CD8 T cell subcluster of the published metadata, “Tcell_06,” and then projected onto the CD8 T cell reference atlas with scAtlasVAE as described above. The query cells were placed in the reference UMAP with the scatlasvae.tl.umap_alignment function with method = "knn" and global random seeds set at 42. The avidity-associated T_EX-int_ and T_EX-KLR_ DEG signatures were then applied as module scores using the Seurat (v5.4.0) AddModuleScore function. The analysis was then restricted to T_EX_ and cycling clusters (GZMK+ T_EX_, ITGAE+ T_EX_, XBP1+ T_EX_, and cycling T cells). Patient03 had two tumor samples that were combined and treated as a single sample. The module scores were z-scored across the cells, where Δz-score = z(T_EX-KLR_ module score) − z(T_EX-int_ module score), and the mean module scores and mean Δz-score were calculated per patient at each time point. Only patients with at least 20 cells across all retained clusters in a given time point were included.

#### Processing and analysis of the Daniel *et al*., 2022 scRNA-seq dataset

The counts, features, barcodes, and metadata of the Daniel *et al*., 2022 LCMV dataset were downloaded from GSE188666. The authors’ published cluster identities and UMAP coordinates were used. We restricted analyses to cells from day 21-post inoculation of LCMV clone 13 (chronic infection model), and counts were log-normalized with the Seurat (v5.4.0) NormalizeData function. The murine T_EX_ subset signatures were defined as the upregulated genes of each of the six T_EX_ clusters (Progenitor T_EX_, Intermediate T_EX_, Early Effector T_EX_, T_EX-KLR_, T_EX_, and Lung T_EX_) against all other clusters using the Wilcoxon rank-sum test implemented through the Seurat FindAllMarkers function (positive markers only, average log_2_FC > 1, gene expressed in at least 10% of cells of the cluster in which the gene was upregulated). Each gene was assigned to the single T_EX_ cluster with the highest log_2_FC, and the 50 highest-ranked genes per cluster were defined as the T_EX_ subset signatures (**Supplementary Table 9**). The mouse gene symbols were mapped to human orthologs with babelgene (v22.9), and the resulting T_EX_ subset signature gene sets were applied as module scores to the HNSCC TILs dataset of this study. For the module score summary heatmap, the mean module score per murine T_EX_ subset signature was z-scored across the HNSCC TILs clusters and plotted with the ComplexHeatmap (v2.22.0) package, with the fill color capped at ±1.5. The CD56^+^ CD8^−^ NK cluster was excluded prior to z-scoring because it is not a CD8 T cell subset. For feature plots, expression was scaled between the 0.1^st^ and 99.9^th^ percentiles within each gene or module score, and cells were ordered by descending expression with the highest on top.

## Statistics

GraphPad Prism (v11.0.0) was used to perform the statistical analyses presented in **Fig. 3**, **Extended Data Fig. 3**, and **Extended Data Fig. 7**. The analyses presented in **Fig. 2** were performed in Python (v3.12.3). All other statistical analyses were performed in R (v4.4.1) using rstatix (v0.7.3). Sample numbers are indicated in the text or figure legends. Unless otherwise specified, all tests were two-sided, error bars represent the mean ± standard deviation, and box plots show the median and interquartile range with the whiskers extending to the minimum or maximum values within 1.5× of the interquartile range. Asterisks denote the following *P* values: \**P* < 0.05, \*\**P* < 0.01, \*\*\**P* < 0.001, \*\*\*\**P* < 0.0001.

