## Extended Data Figures for "High-avidity TCR signaling drives an NKG2A-positive CD8 T cell exhaustion state in human tumors"

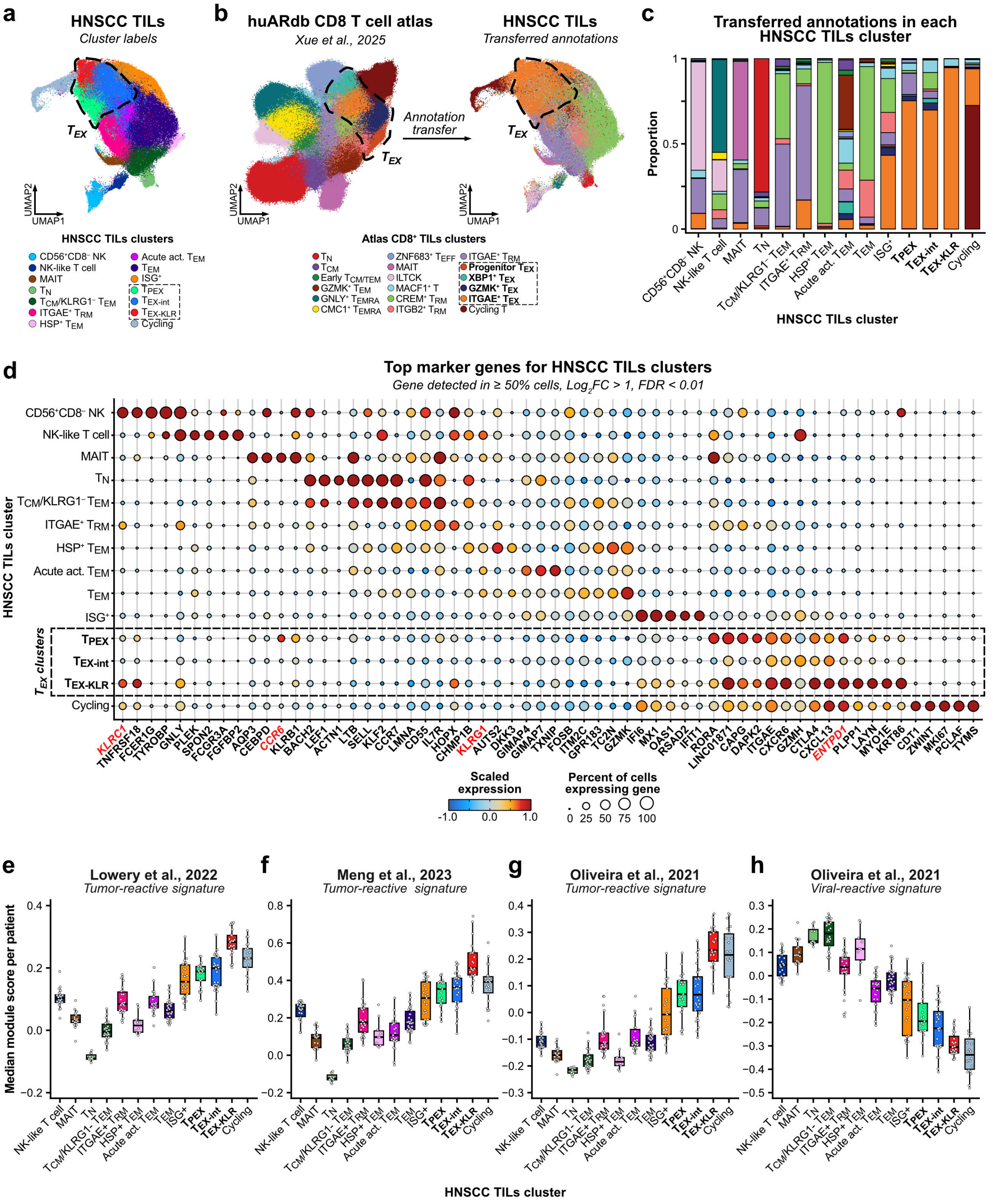
a
HNSCC TILs
Cluster labels
TEX
UMAP1
UMAP2
HNSCC TILs clusters
CD56+CD8− NK
NK-like T cell
MAIT
TN
TCM/KLRG1− TEM
ITGAE+ TRM
HSP+ TEM
Acute act. TEM
TEM
ISG+
TPEX
TEX-int
TEX-KLR
Cycling
b
huARdb CD8 T cell atlas
Xue et al., 2025
Annotation transfer
HNSCC TILs
Transferred annotations
Atlas CD8+ TILs clusters
TN
TCM
Early TCM/TEM
GZMK+ TEM
GNLY+ TEMRA
CMC1+ TEMRA
ZNF683+ TEFF
MAIT
ILTCK
MACF1+ T
CREM+ TRM
ITGB2+ TRM
ITGAE+ TRM
Progenitor TEX
XBP1+ TEX
GZMK+ TEX
ITGAE+ TEX
Cycling T
c
Transferred annotations in each HNSCC TILs cluster
Proportion
HNSCC TILs cluster
d
Top marker genes for HNSCC TILs clusters
Gene detected in ≥ 50% cells, Log2FC > 1, FDR < 0.01
HNSCC TILs cluster
TEX clusters
KLRC1 TNFRSF18 FCER1G TYROBP GNLY PLEK SPON2 FCGR3A FGFBP2 AQP3 CEBPD CCR6 KLRB1 BACH2 LEF1 ACTN1 LTB SELL KLF2 CCR7 LMNA CD55 IL7R HOPX CHMP1B KLRG1 AUTS2 DKK3 GIMAP4 GIMAP7 TXNIP FOSB ITM2C GPR183 TC2N GZMK IFI6 MX1 OAS1 RSAD2 IFIT1 RORA LINC01871 CAPG DAPK2 ITGAE CXCR6 GZMH CTLA4 CXCL13 ENTPD1 PLPP1 LAYN MYO1E KRT86 CDT1 ZWINT MKI67 PCLAF TYMS
Scaled expression
Percent of cells expressing gene
e
Lowery et al., 2022
Tumor-reactive signature
Median module score per patient
f
Meng et al., 2023
Tumor-reactive signature
g
Oliveira et al., 2021
Tumor-reactive signature
h
Oliveira et al., 2021
Viral-reactive signature
HNSCC TILs cluster

**Extended Data Figure 1. Classification of scRNA-seq clusters**

**a**, UMAP plot of the HNSCC TILs dataset.

**b**, Annotation transfer from the huARdb CD8 T cell atlas (*left*; Xue *et al.*, 2025) to the HNSCC TILs dataset (*right*); the published UMAP coordinates and clusters of the huARdb dataset are shown.

**c**, Proportions of the transferred annotations within each HNSCC TILs cluster.

**d**, Dot plot of top marker genes per HNSCC TILs cluster.

**e–h**, Per-patient median module scores in each scRNA-seq cluster for tumor-reactive signatures from (**e**) Lowery *et al.*, 2022, (**f**) Meng *et al.*, 2023, and (**g**) Oliveira *et al.*, 2021, and (**h**) for a viral-reactive signature from Oliveira *et al.*, 2021.

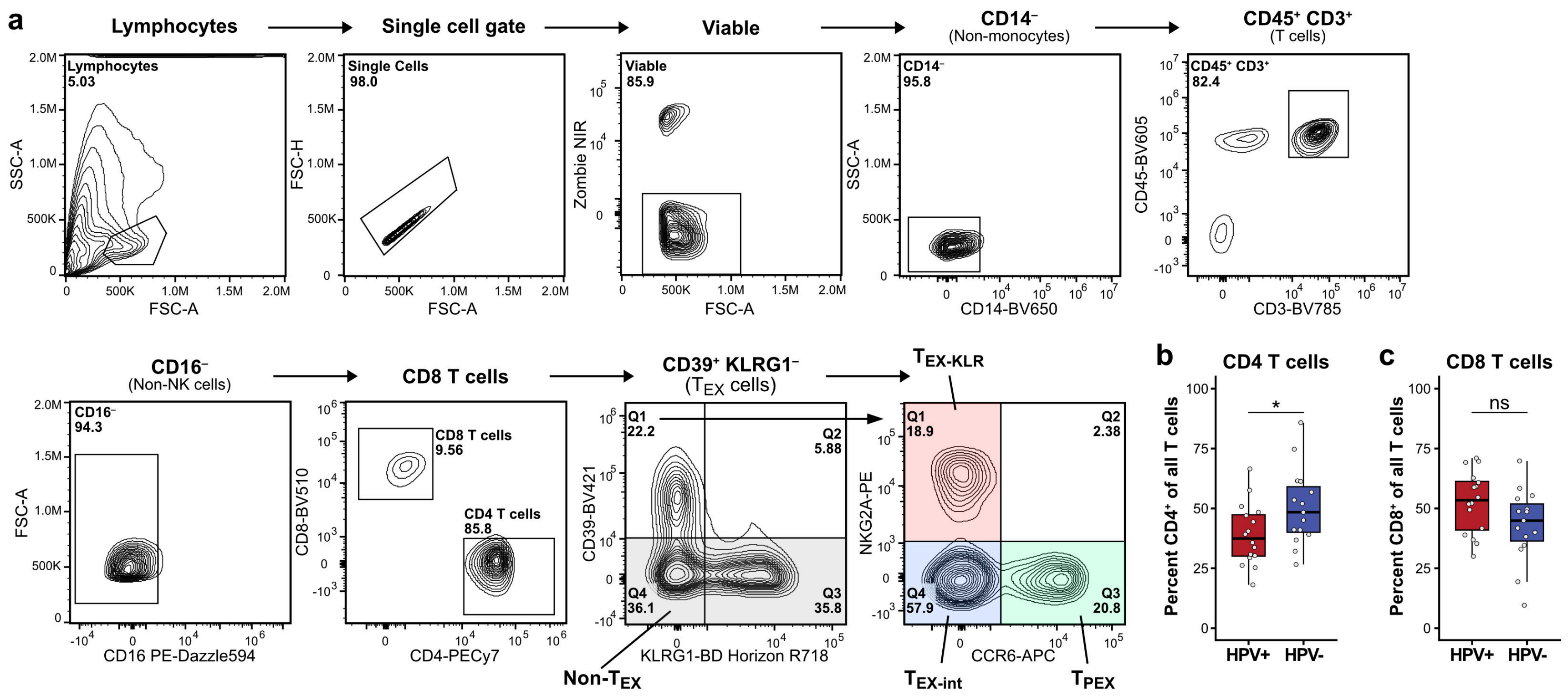

a
Lymphocytes
Single cell gate
Viable
CD14⁻ (Non-monocytes)
CD45⁺ CD3⁺ (T cells)
Lymphocytes 5.03
Single Cells 98.0
Viable 85.9
CD14⁻ 95.8
CD45⁺ CD3⁺ 82.4
SSC-A
FSC-A
FSC-H
Zombie NIR
CD14-BV650
CD45-BV605
CD3-BV785
CD16⁻ (Non-NK cells)
CD8 T cells
CD39⁺ KLRG1⁻ (TEX cells)
TEX-KLR
CD16⁻ 94.3
CD8 T cells 9.56
CD4 T cells 85.8
Q1 22.2
Q2 5.88
Q3 35.8
Q4 36.1
Q1 18.9
Q2 2.38
Q3 20.8
Q4 57.9
FSC-A
CD16 PE-Dazzle594
CD8-BV510
CD4-PECy7
CD39-BV421
KLRG1-BD Horizon R718
NKG2A-PE
CCR6-APC
Non-TEX
TEX-int
TPEX
b
CD4 T cells
Percent CD4⁺ of all T cells
*
HPV+
HPV-
c
CD8 T cells
Percent CD8⁺ of all T cells
ns
HPV+
HPV-

**Extended Data Figure 2. Flow cytometry of HNSCC patient tumors**

**a**, Flow cytometry gating strategy for the HNSCC patient tumor samples.

**b–c**, Percentage of (**b**) CD4 T cells and (**c**) CD8 T cells among all T cells in HPV+ and HPV- tumor samples in the flow cytometry data (*n* = 16 HPV+ and 15 HPV- tumors). Unadjusted *P* values from a Wilcoxon rank-sum test are shown.

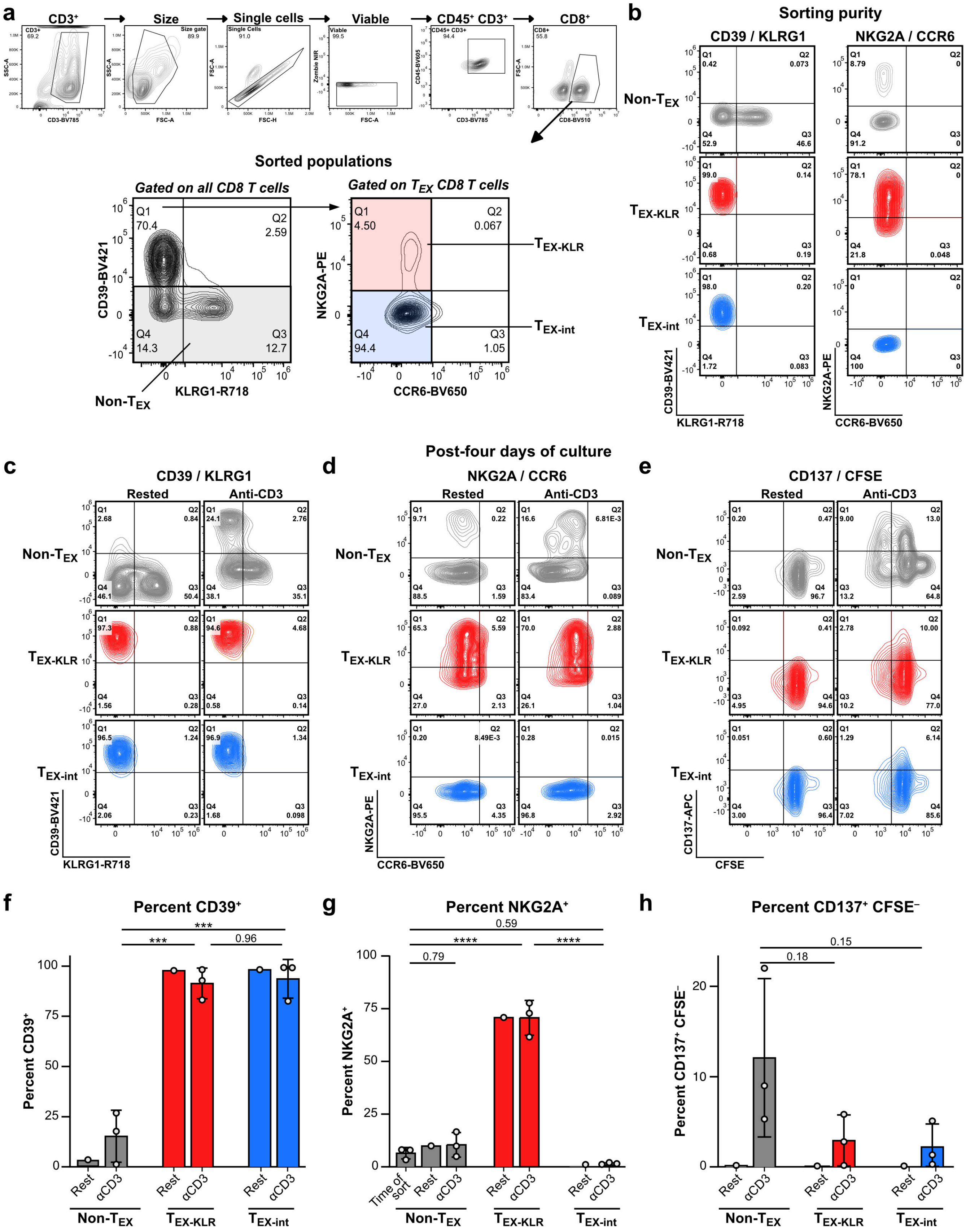

a
CD3⁺ → Size → Single cells → Viable → CD45⁺ CD3⁺ → CD8⁺
Sorted populations
Gated on all CD8 T cells
Gated on TEX CD8 T cells
CD39-BV421
KLRG1-R718
NKG2A-PE
CCR6-BV650
Non-TEX
TEX-KLR
TEX-int
b
Sorting purity
CD39 / KLRG1
NKG2A / CCR6
Post-four days of culture
c
CD39 / KLRG1
d
NKG2A / CCR6
e
CD137 / CFSE
Rested
Anti-CD3
CD137-APC
CFSE
f
Percent CD39⁺
g
Percent NKG2A⁺
h
Percent CD137⁺ CFSE⁻
Time of sort
Rest
αCD3

**Extended Data Figure 3. *Ex vivo* stimulation of sorted non-$T_{EX}$, $T_{EX\text{-}int}$, and $T_{EX\text{-}KLR}$ cells**

**a**, Flow cytometry gating strategy for sorting non-$T_{EX}$, $T_{EX\text{-}int}$, and $T_{EX\text{-}KLR}$ cells from freshly dissociated HNSCC patient tumor samples.

**b**, Sorting purity assessed by analyzing the cells immediately post-sort.

**c–h**, Cells were rested or stimulated with immobilized anti-CD3 for four days *in vitro* and then analyzed by flow cytometry ($n$ = 1 tumor sample for rest and 3 tumor samples for anti-CD3 stimulation).

**c–e**, Representative flow cytometry plots of (**c**) CD39 and KLRG1 expression, (**d**) NKG2A and CCR6 expression, and (**e**) CD137 expression and CFSE.

**f–h**, Percentage of (**f**) $CD39^+$, (**g**) $NKG2A^+$, and (**h**) $CD137^+$ $CFSE^-$ cells. *P* values from a repeated-measures one-way ANOVA with Tukey's correction are shown for the anti-CD3 conditions.

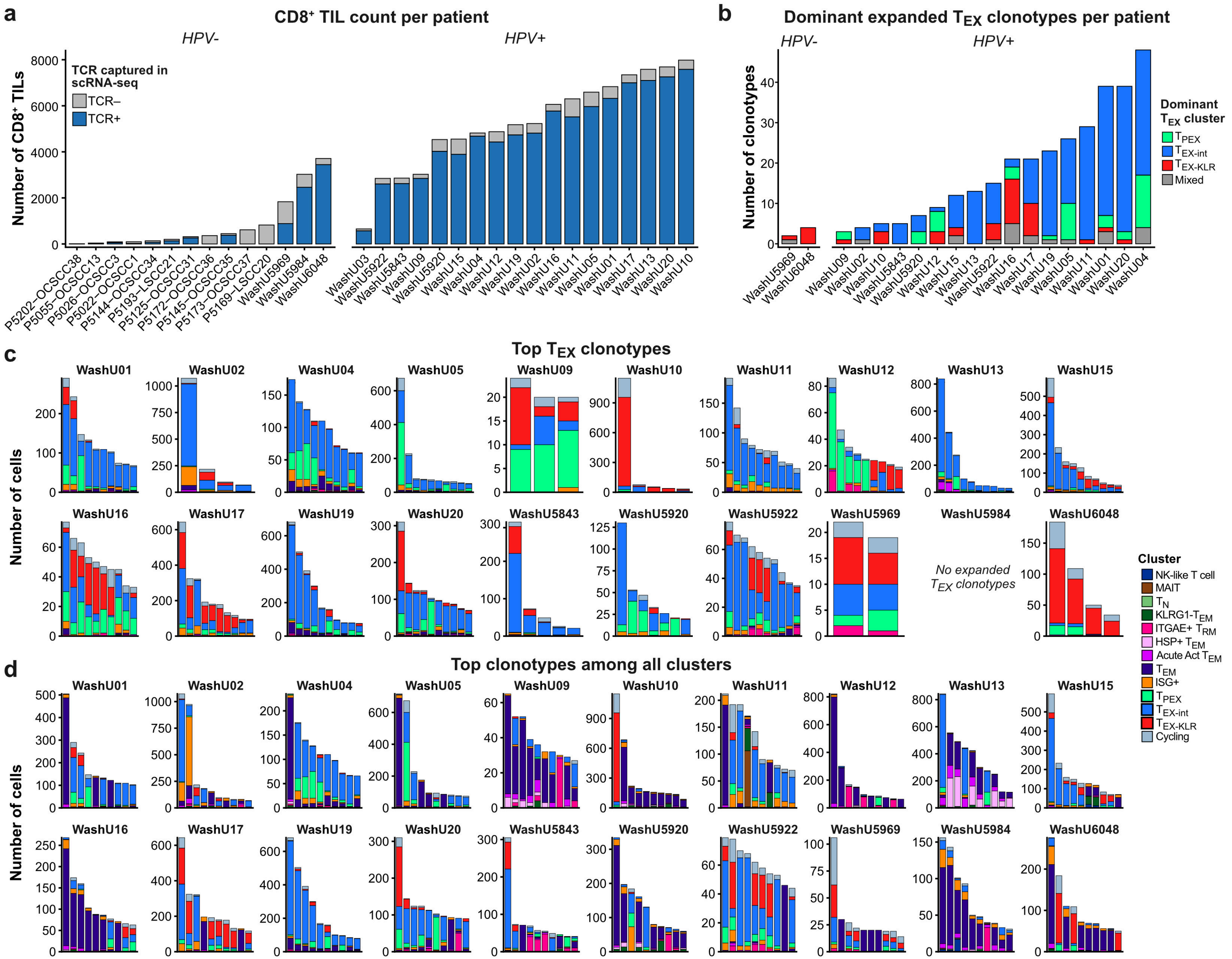
a
CD8+ TIL count per patient
HPV-
HPV+
Number of CD8+ TILs
TCR captured in scRNA-seq
TCR–
TCR+
b
Dominant expanded TEX clonotypes per patient
HPV-
HPV+
Number of clonotypes
Dominant TEX cluster
TPEX
TEX-int
TEX-KLR
Mixed
c
Top TEX clonotypes
Number of cells
No expanded TEX clonotypes
Cluster
NK-like T cell
MAIT
TN
KLRG1-TEM
ITGAE+ TRM
HSP+ TEM
Acute Act TEM
TEM
ISG+
TPEX
TEX-int
TEX-KLR
Cycling
d
Top clonotypes among all clusters
Number of cells

**Extended Data Figure 4. Phenotypes of expanded CD8 T cell clonotypes**

**a**, Number of $CD8^+$ TILs with or without a TCR captured by scTCR-seq per patient.

**b**, Number of expanded $T_{EX}$ clonotypes dominant within each $T_{EX}$ cluster per patient.

**c–d**, Cluster composition of up to 10 of the largest clonotypes per patient among (**c**) expanded $T_{EX}$ clonotypes or (**d**) all CD8 T cells. Only patients with at least 600 cells with a TCR are shown.

### Figure 2 immunophenotyping

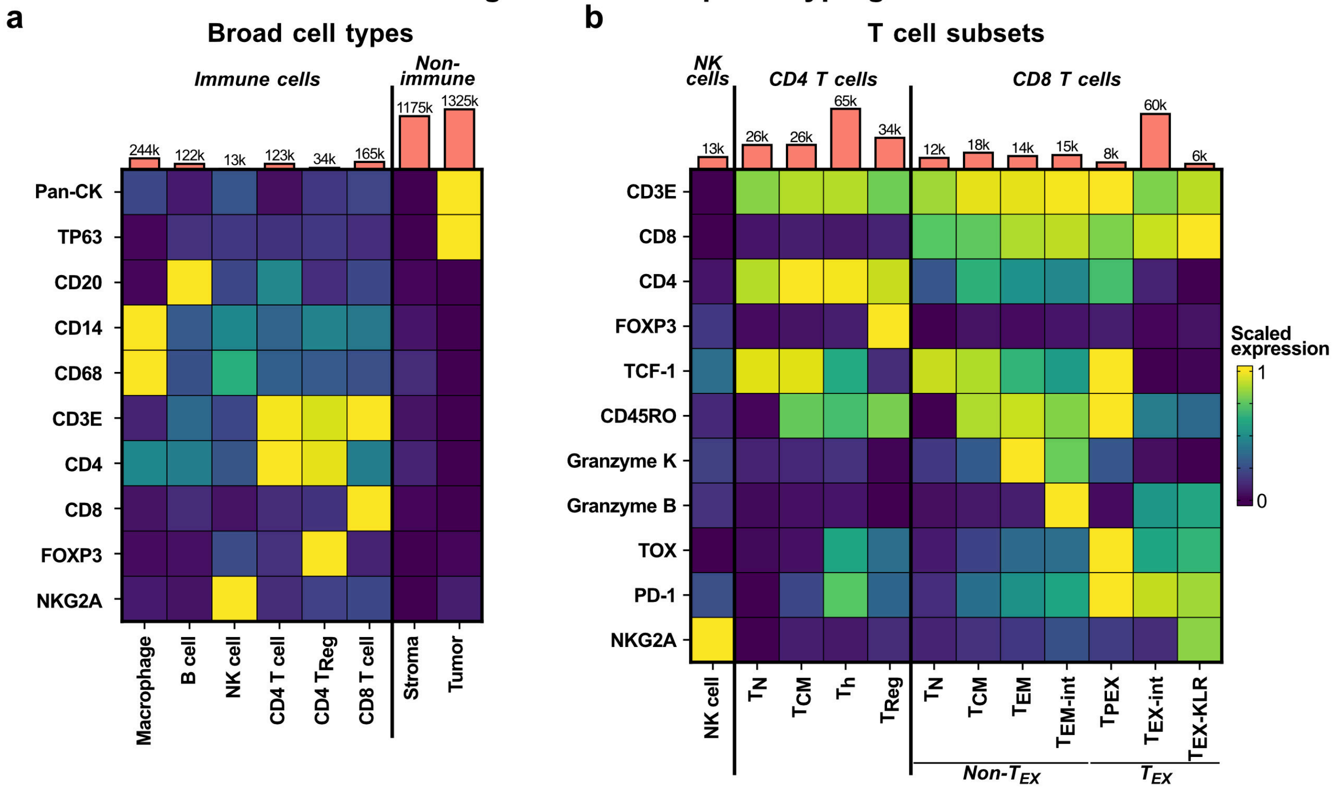


### Figure 6 immunophenotyping

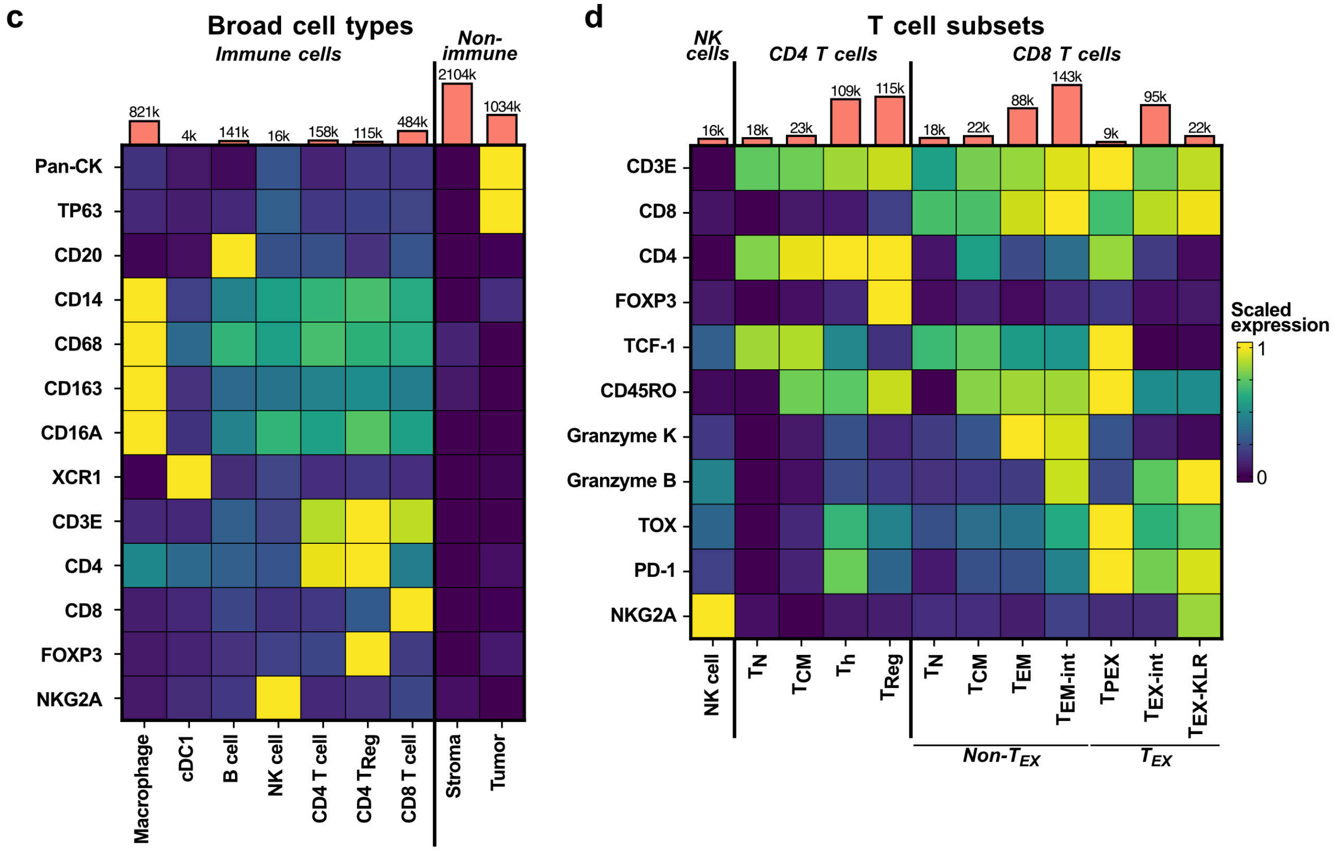

**Extended Data Figure 5. Immunofluorescence data phenotyping**

**a–b**, Immunophenotyping marker proteins used in **Fig. 2** for (**a**) broad cell types and (**b**) T cell subsets; (**b**) is also shown as **Fig. 2b**.

**c–d**, Immunophenotyping marker proteins used in **Fig. 6** for (**c**) broad cell types and (**d**) T cell subsets. Total count of cells per type is indicated above as a labeled bar in (**a–d**).

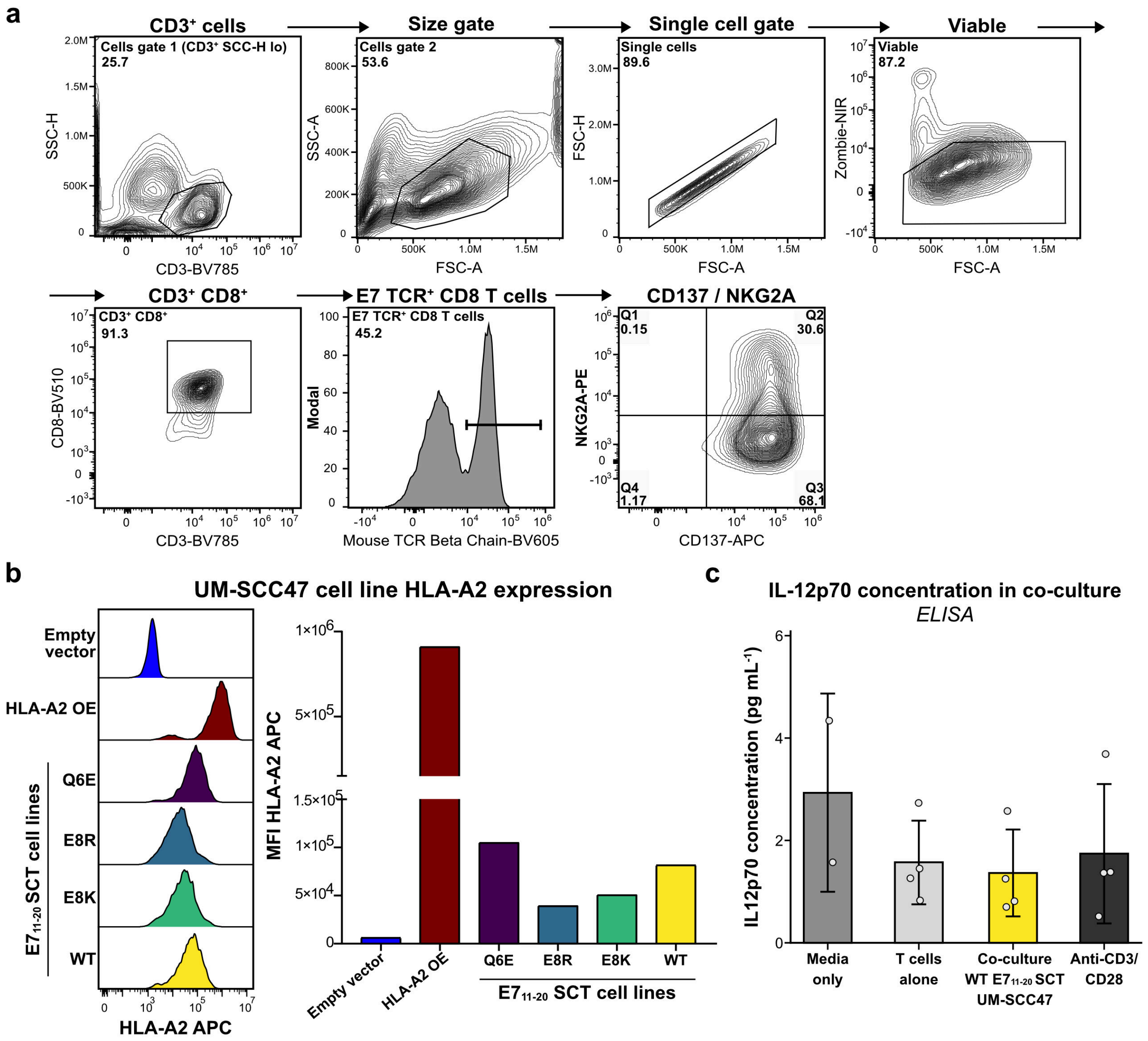
a
CD3⁺ cells
Cells gate 1 (CD3⁺ SCC-H lo)
25.7
SSC-H
CD3-BV785
Size gate
Cells gate 2
53.6
SSC-A
FSC-A
Single cell gate
Single cells
89.6
FSC-H
FSC-A
Viable
Viable
87.2
Zombie-NIR
FSC-A
CD3⁺ CD8⁺
CD3⁺ CD8⁺
91.3
CD8-BV510
CD3-BV785
E7 TCR⁺ CD8 T cells
E7 TCR⁺ CD8 T cells
45.2
Modal
Mouse TCR Beta Chain-BV605
CD137 / NKG2A
Q1
0.15
Q2
30.6
Q3
68.1
Q4
1.17
NKG2A-PE
CD137-APC
b
UM-SCC47 cell line HLA-A2 expression
Empty vector
HLA-A2 OE
Q6E
E8R
E8K
WT
E7 11-20 SCT cell lines
HLA-A2 APC
MFI HLA-A2 APC
Empty vector
HLA-A2 OE
Q6E
E8R
E8K
WT
E7 11-20 SCT cell lines
c
IL-12p70 concentration in co-culture
ELISA
IL12p70 concentration (pg mL⁻¹)
Media only
T cells alone
Co-culture WT E7 11-20 SCT UM-SCC47
Anti-CD3/ CD28

**Extended Data Figure 6. *In vitro* model of TCR avidity in the UM-SCC47 cell line**

**a**, Flow cytometry gating strategy for analysis of E7 $TCR^+$ CD8 T cells following *in vitro* co-culture.

**b**, Flow cytometry histograms of HLA-A2 expression on UM-SCC47 cell lines normalized to the mode (*left*) and quantified as mean fluorescence intensity (MFI; *right*).

**c**, IL-12p70 concentration in the media following culture of E7 $TCR^+$ CD8 T cells in the indicated condition for 48 h (*n* = 4 donors; E7 $TCR^+$ CD8 T cells generated from donors ID85, ID267, ID481, and ID948 were used).

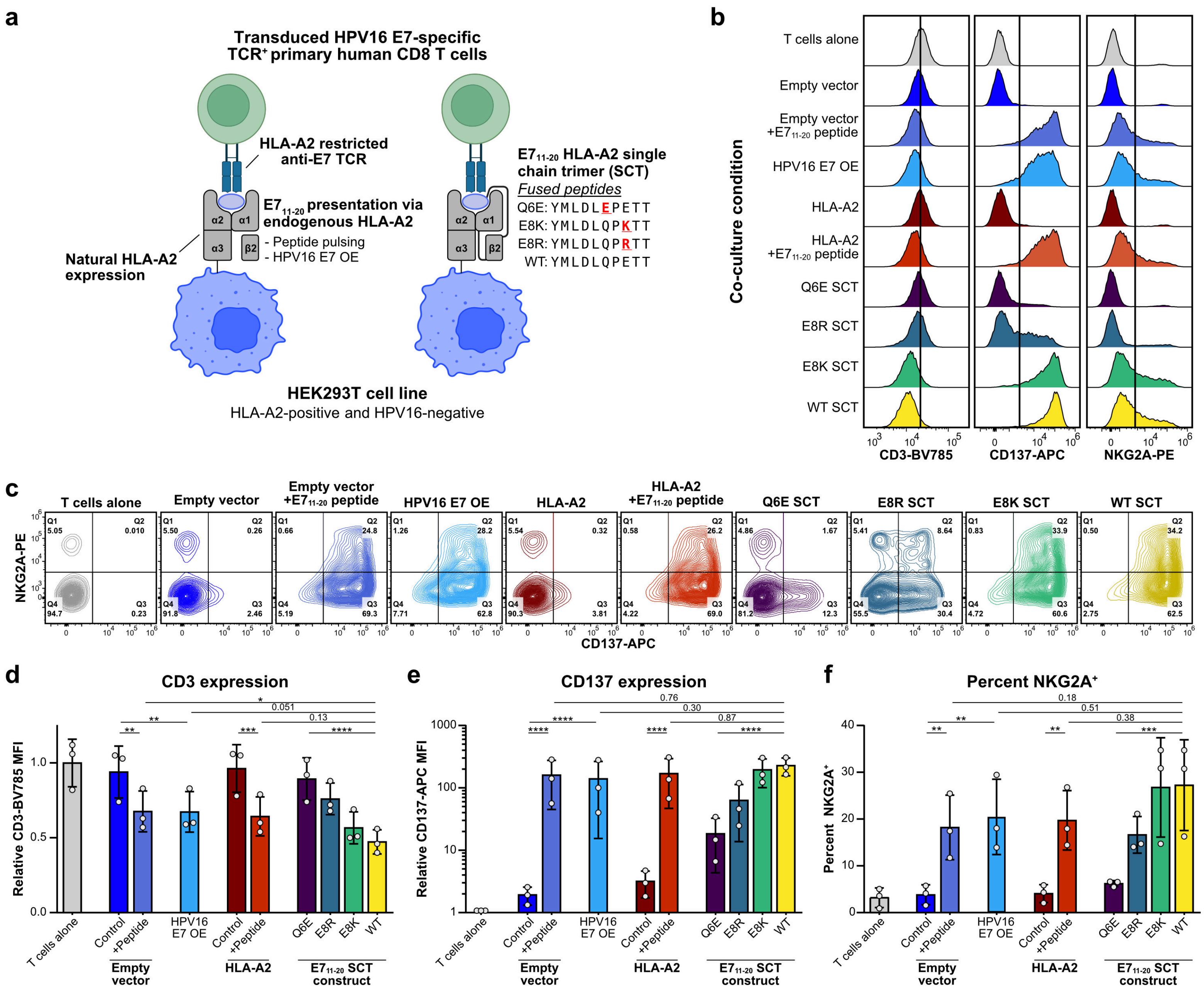

a
Transduced HPV16 E7-specific TCR+ primary human CD8 T cells
HLA-A2 restricted anti-E7 TCR
E7 11-20 presentation via endogenous HLA-A2
- Peptide pulsing
- HPV16 E7 OE
Natural HLA-A2 expression
E7 11-20 HLA-A2 single chain trimer (SCT)
Fused peptides
Q6E: YMLDLEPETT
E8K: YMLDLQPKTT
E8R: YMLDLQPRTT
WT: YMLDLQPETT
HEK293T cell line
HLA-A2-positive and HPV16-negative
b
Co-culture condition
T cells alone
Empty vector
Empty vector +E7 11-20 peptide
HPV16 E7 OE
HLA-A2
HLA-A2 +E7 11-20 peptide
Q6E SCT
E8R SCT
E8K SCT
WT SCT
CD3-BV785
CD137-APC
NKG2A-PE
c
NKG2A-PE
CD137-APC
d
CD3 expression
Relative CD3-BV785 MFI
e
CD137 expression
Relative CD137-APC MFI
f
Percent NKG2A+
Percent NKG2A+
T cells alone
Control
+Peptide
HPV16 E7 OE
Empty vector
HLA-A2
Q6E
E8R
E8K
WT
E7 11-20 SCT construct

**Extended Data Figure 7. High-avidity TCR signaling promotes NKG2A expression in co-cultures with HEK293T target cells**

**a**, Schematic of HEK293T target cell lines, including peptide pulsing and HPV16 E7 overexpression (OE; *left*) and the WT and mutant $E7_{11–20}$ SCT cell lines (*right*).

**b–c**, Representative flow cytometry (**b**) histograms of CD3, CD137, and NKG2A expression normalized to the mode and (**c**) plots of NKG2A and CD137 expression on E7 $TCR^+$ CD8 T cells following 48 h co-culture. Data for E7 $TCR^+$ CD8 T cells generated from donor LRS071724 are shown.

**d–f**, Quantification of (**d**) CD3 and (**e**) CD137 mean fluorescence intensity (MFI) relative to E7 $TCR^+$ CD8 T cells cultured alone and (**f**) percentage of $NKG2A^+$ E7 $TCR^+$ CD8 T cells ($n$ = 3 donors; E7 $TCR^+$ CD8 T cells generated from donors LRS072323, LRS050924, and LRS071724 were used). *P* values from a repeated-measures one-way ANOVA with Sidak's correction are shown.

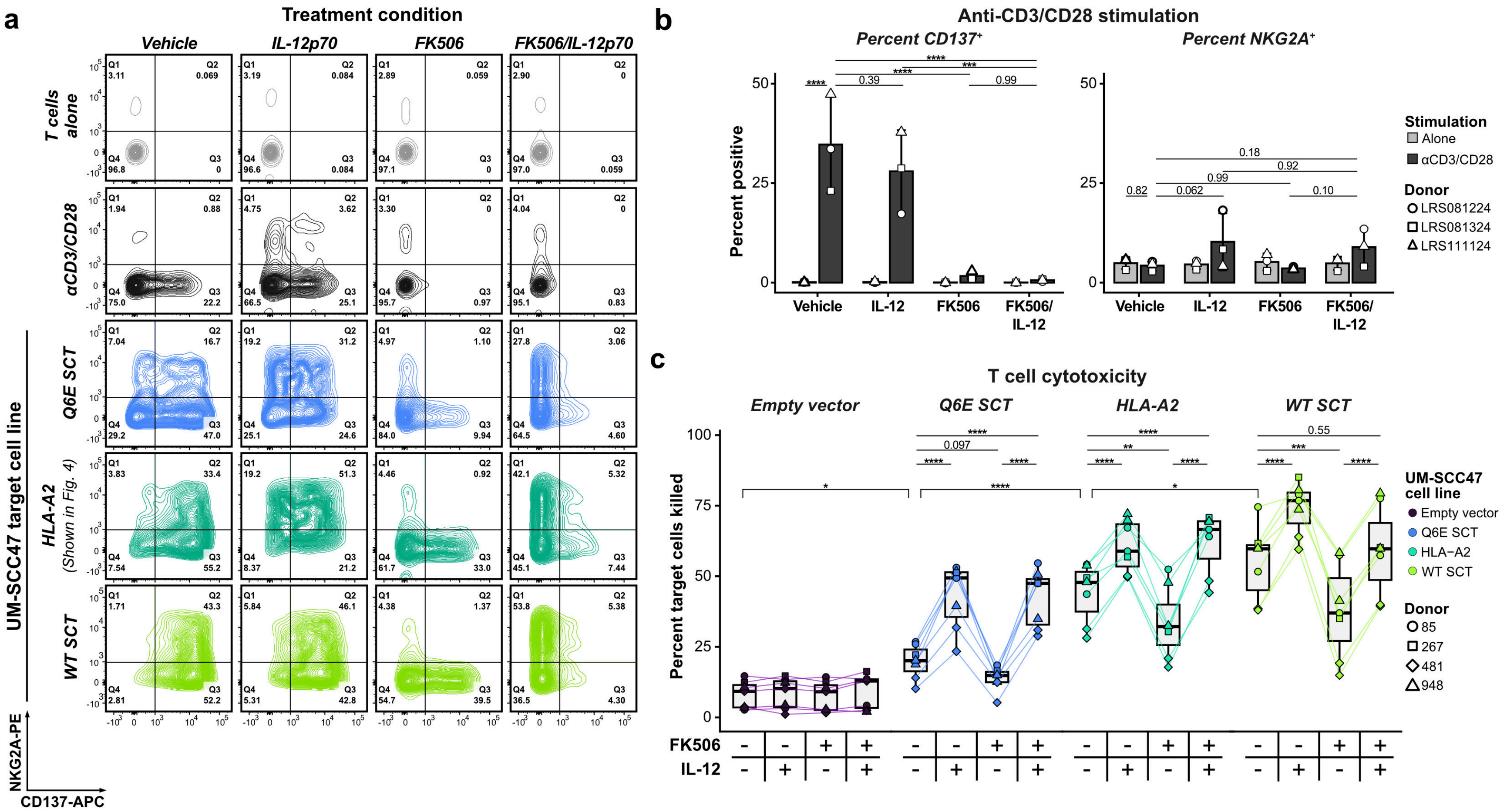

a
Treatment condition
Vehicle
IL-12p70
FK506
FK506/IL-12p70
T cells alone
αCD3/CD28
UM-SCC47 target cell line
Q6E SCT
HLA-A2
(Shown in Fig. 4)
WT SCT
NKG2A-PE
CD137-APC
b
Anti-CD3/CD28 stimulation
Percent CD137+
Percent NKG2A+
Percent positive
Vehicle
IL-12
FK506
FK506/IL-12
Stimulation
Alone
αCD3/CD28
Donor
LRS081224
LRS081324
LRS111124
c
T cell cytotoxicity
Empty vector
Q6E SCT
HLA-A2
WT SCT
Percent target cells killed
UM-SCC47 cell line
Empty vector
Q6E SCT
HLA−A2
WT SCT
Donor
85
267
481
948
FK506
IL-12

**Extended Data Figure 8. NFAT and IL-12 independently regulate NKG2A expression and cytotoxicity**

**a**, Representative flow cytometry plots of NKG2A and CD137 expression on E7 TCR$^+$ CD8 T cells following 48 h culture in the indicated condition. Data for E7 TCR$^+$ CD8 T cells generated from donor LRS081324 are shown.

**b**, Percentage of CD137$^+$ (*left*) and NKG2A$^+$ (*right*) E7 TCR$^+$ CD8 T cells following rest or anti-CD3 and anti-CD28 stimulation for 48 h ($n$ = 3 donors).

**c**, Percentage of UM-SCC47 target cells killed after 48 h co-culture with E7 TCR$^+$ CD8 T cells ($n$ = 7 replicates from 4 donors).

*P* values from a repeated-measures one-way ANOVA with Tukey's correction for multiple comparisons are shown in (**b**, **c**).

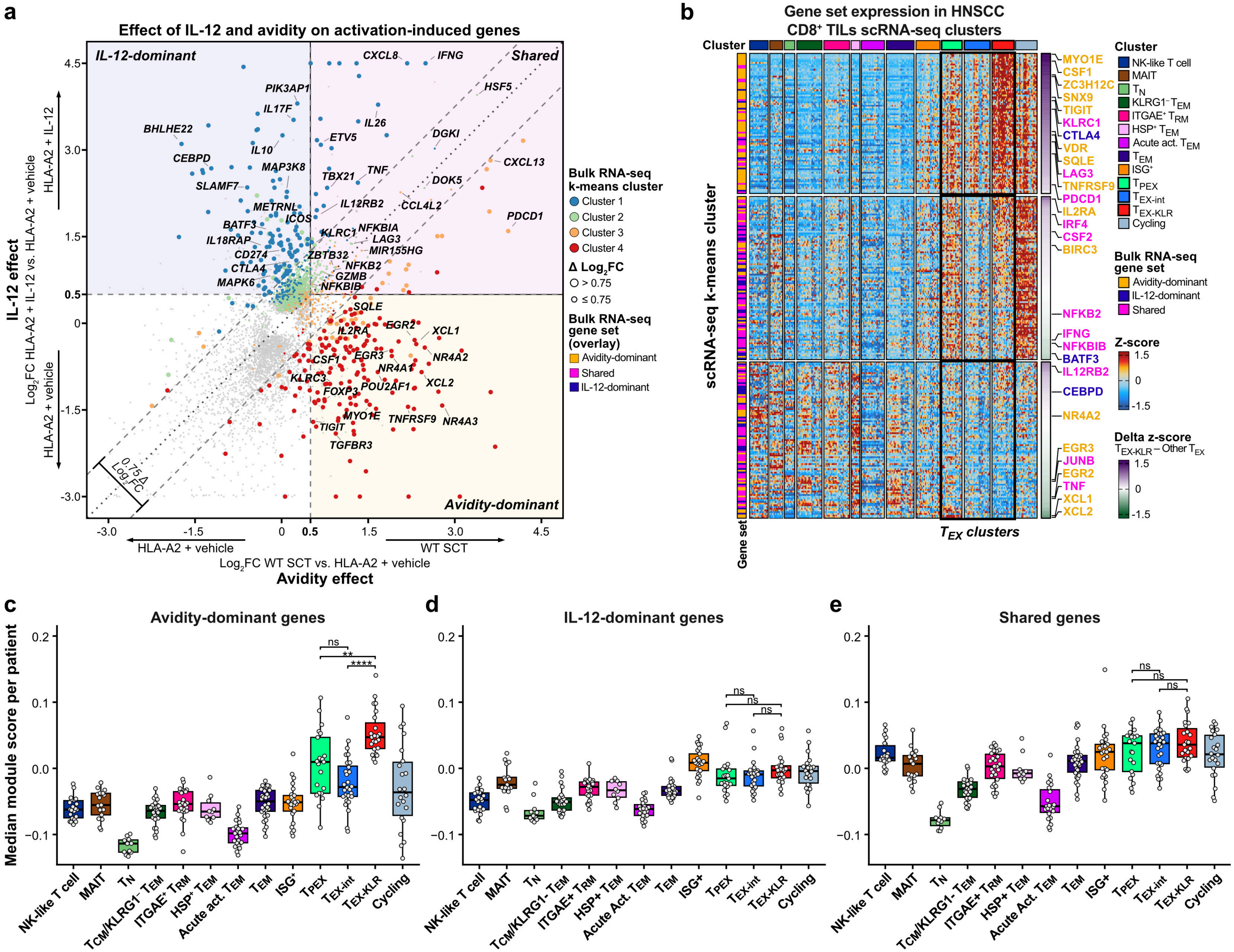
a
Effect of IL-12 and avidity on activation-induced genes
IL-12-dominant
Shared
Avidity-dominant
IL-12 effect
Log2FC HLA-A2 + IL-12 vs. HLA-A2 + vehicle
HLA-A2 + IL-12
HLA-A2 + vehicle
Avidity effect
Log2FC WT SCT vs. HLA-A2 + vehicle
HLA-A2 + vehicle
WT SCT
0.75 Δ Log2FC
Bulk RNA-seq k-means cluster
Cluster 1
Cluster 2
Cluster 3
Cluster 4
Δ Log2FC
> 0.75
≤ 0.75
Bulk RNA-seq gene set (overlay)
Avidity-dominant
Shared
IL-12-dominant
b
Gene set expression in HNSCC CD8+ TILs scRNA-seq clusters
Cluster
scRNA-seq k-means cluster
Gene set
TEX clusters
NK-like T cell
MAIT
TN
KLRG1− TEM
ITGAE+ TRM
HSP+ TEM
Acute act. TEM
TEM
ISG+
TPEX
TEX-int
TEX-KLR
Cycling
Bulk RNA-seq gene set
Avidity-dominant
IL-12-dominant
Shared
Z-score
Delta z-score
TEX-KLR − Other TEX
c
Avidity-dominant genes
Median module score per patient
d
IL-12-dominant genes
e
Shared genes
NK-like T cell
MAIT
TN
TCM/KLRG1- TEM
ITGAE+ TRM
HSP+ TEM
Acute act. TEM
TEM
ISG+
TPEX
TEX-int
TEX-KLR
Cycling
ns
**
****

**Extended Data Figure 9. IL-12 signaling and high-avidity TCR signaling trigger shared and distinct gene expression programs**

**a**, Differential gene expression of E7 TCR$^+$ CD8 T cells co-cultured with UM-SCC47 target cells: HLA-A2 cells plus IL-12 (*y-axis*) or WT SCT cells (*x-axis*), each relative to co-culture with HLA-A2 cells plus vehicle as the baseline-activation reference. Gene data points are colored by their k-means cluster from the **Fig. 4d** heatmap. Diagonal lines indicate a 0.75 absolute $\log_2$FC difference between the contrasts, and colored overlays indicate the gene sets (see **Methods**).

**b**, Z-scored mean RNA expression of the genes in the IL-12-dominant, avidity-dominant, and shared gene sets for the cells of each patient per cluster. Genes are k-means clustered and ordered by the difference in the mean z-score of the $T_{EX\text{-}KLR}$ columns and the $T_{PEX}$ and $T_{EX\text{-}int}$ columns.

**c–e**, Per-patient median module scores for the (**c**) avidity-dominant, (**d**) IL-12-dominant, and (**e**) shared gene sets in each scRNA-seq cluster ($n$ = 21 $T_{PEX}$, 28 $T_{EX\text{-}int}$, and 23 $T_{EX\text{-}KLR}$ patients). *P* values from a Wilcoxon rank-sum test with Benjamini–Hochberg correction are shown.

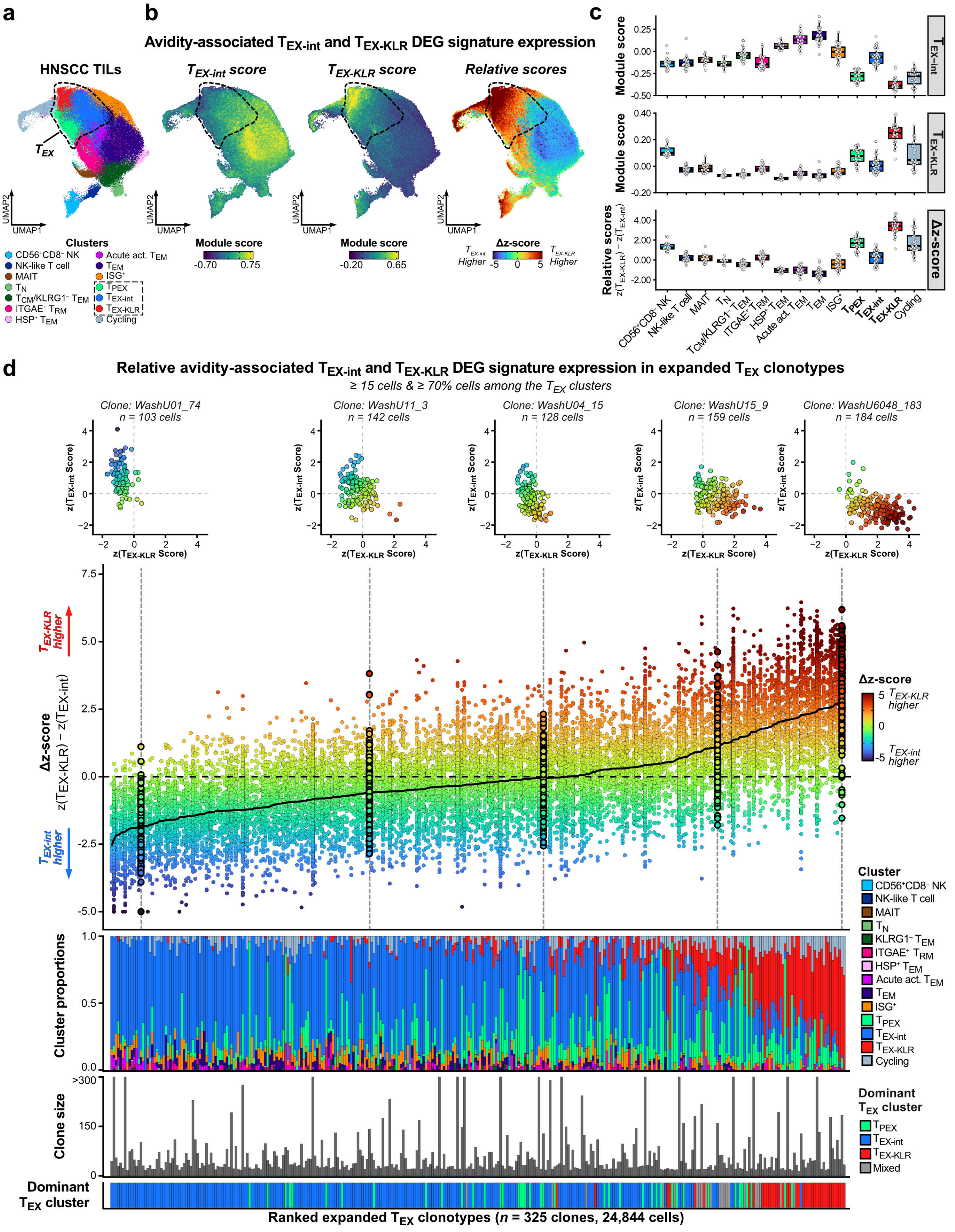
a
b
c
Avidity-associated $T_{EX-int}$ and $T_{EX-KLR}$ DEG signature expression
HNSCC TILs
$T_{EX}$
$T_{EX-int}$ score
$T_{EX-KLR}$ score
Relative scores
UMAP1
UMAP2
Clusters
CD56⁺CD8⁻ NK
NK-like T cell
MAIT
$T_N$
$T_{CM}$/KLRG1⁻ $T_{EM}$
ITGAE⁺ $T_{RM}$
HSP⁺ $T_{EM}$
Acute act. $T_{EM}$
$T_{EM}$
ISG⁺
$T_{PEX}$
$T_{EX-int}$
$T_{EX-KLR}$
Cycling
Module score
-0.70 0.75
Module score
-0.20 0.65
Δz-score
$T_{EX-int}$ Higher
-5 0 5
$T_{EX-KLR}$ Higher
Module score
Module score
Relative scores $z(T_{EX-KLR}) - z(T_{EX-int})$
$T_{EX-int}$
$T_{EX-KLR}$
Δz-score
d
Relative avidity-associated $T_{EX-int}$ and $T_{EX-KLR}$ DEG signature expression in expanded $T_{EX}$ clonotypes
≥ 15 cells & ≥ 70% cells among the $T_{EX}$ clusters
Clone: WashU01_74 n = 103 cells
Clone: WashU11_3 n = 142 cells
Clone: WashU04_15 n = 128 cells
Clone: WashU15_9 n = 159 cells
Clone: WashU6048_183 n = 184 cells
$z(T_{EX-int}$ Score)
$z(T_{EX-KLR}$ Score)
$T_{EX-KLR}$ higher
$T_{EX-int}$ higher
Δz-score $z(T_{EX-KLR}) - z(T_{EX-int})$
Δz-score
5 $T_{EX-KLR}$ higher
0
-5 $T_{EX-int}$ higher
Cluster proportions
Cluster
CD56⁺CD8⁻ NK
NK-like T cell
MAIT
$T_N$
KLRG1⁻ $T_{EM}$
ITGAE⁺ $T_{RM}$
HSP⁺ $T_{EM}$
Acute act. $T_{EM}$
$T_{EM}$
ISG⁺
$T_{PEX}$
$T_{EX-int}$
$T_{EX-KLR}$
Cycling
Clone size
>300
150
Dominant $T_{EX}$ cluster
$T_{PEX}$
$T_{EX-int}$
$T_{EX-KLR}$
Mixed
Dominant $T_{EX}$ cluster
Ranked expanded $T_{EX}$ clonotypes (n = 325 clones, 24,844 cells)

**Extended Data Figure 10. Expanded $T_{EX}$ clonotypes in HNSCC occupy a reciprocal axis between $T_{EX\text{-}int}$ and $T_{EX\text{-}KLR}$ states**

**a**, UMAP plot of the HNSCC TILs dataset.

**b**, Module scores for the avidity-associated $T_{EX\text{-}int}$ (*left*) and $T_{EX\text{-}KLR}$ (*middle*) DEG signatures (from **Fig. 4g**) and the relative module scores represented as the Δz-score (*right*; Δz-score = z-scored $T_{EX\text{-}KLR}$ module score minus z-scored $T_{EX\text{-}int}$ module score).

**c**, Per-patient mean module scores in each scRNA-seq cluster for the avidity-associated $T_{EX\text{-}int}$ (*top*) and $T_{EX\text{-}KLR}$ (*middle*) DEG signatures and the Δz-score (*bottom*).

**d**, Relative avidity-associated DEG signature module scores in expanded $T_{EX}$ clonotypes, where cells of each clonotype are arranged into columns and filled by their Δz-score, and clonotypes are ordered by their mean Δz-score. Cells of highlighted clonotypes are shown on a scatter plot. The cluster proportions, size, and dominant $T_{EX}$ cluster of each clonotype are shown below.

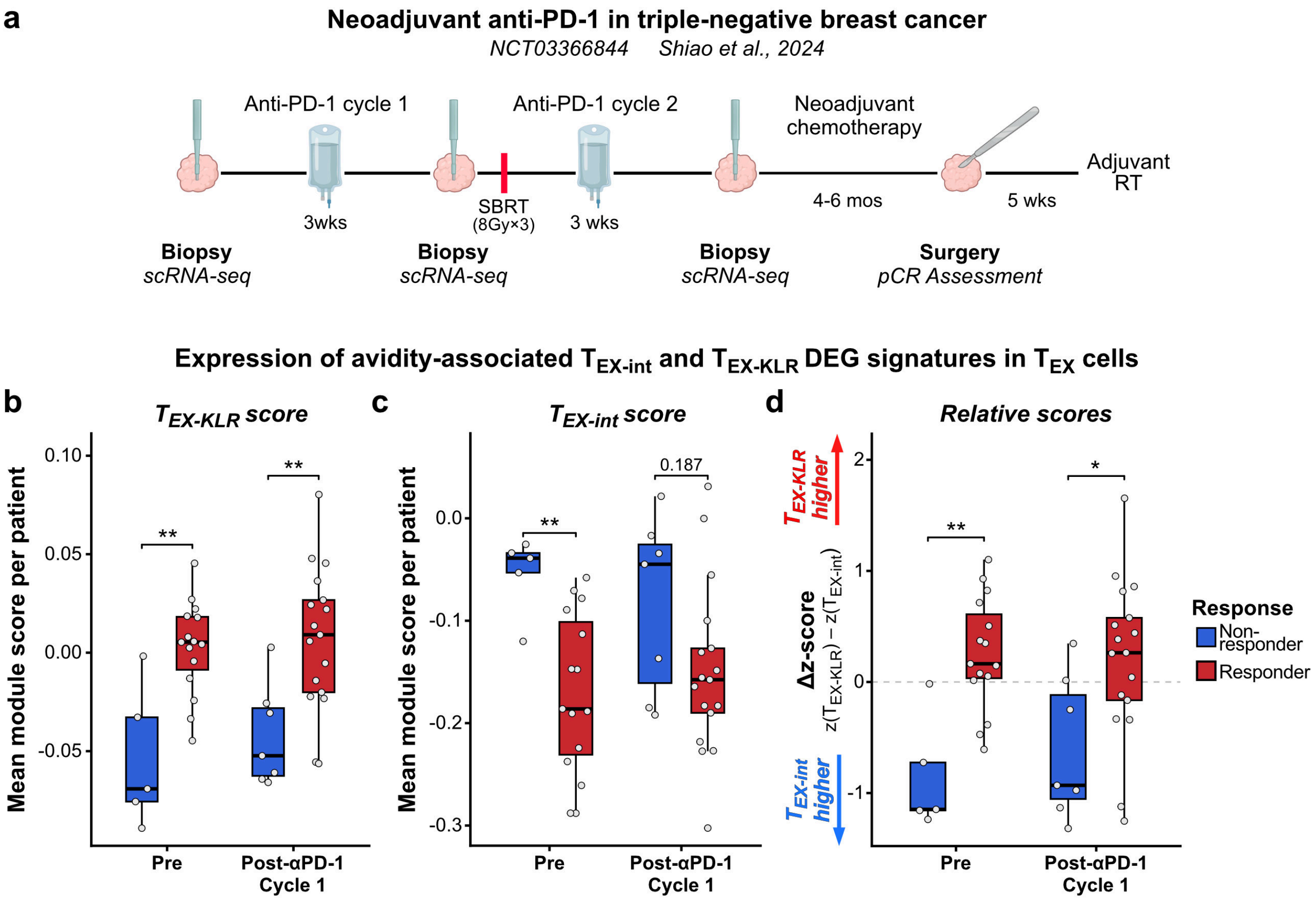

a
Neoadjuvant anti-PD-1 in triple-negative breast cancer
NCT03366844 Shiao et al., 2024
Anti-PD-1 cycle 1
Anti-PD-1 cycle 2
Neoadjuvant chemotherapy
Adjuvant RT
3wks
SBRT (8Gy×3)
3 wks
4-6 mos
5 wks
Biopsy scRNA-seq
Biopsy scRNA-seq
Biopsy scRNA-seq
Surgery pCR Assessment
Expression of avidity-associated TEX-int and TEX-KLR DEG signatures in TEX cells
b
TEX-KLR score
Mean module score per patient
**
**
Pre
Post-αPD-1 Cycle 1
c
TEX-int score
Mean module score per patient
**
0.187
Pre
Post-αPD-1 Cycle 1
d
Relative scores
TEX-KLR higher
TEX-int higher
Δz-score
z(TEX-KLR) − z(TEX-int)
**
*
Pre
Post-αPD-1 Cycle 1
Response
Non-responder
Responder

**Extended Data Figure 11. The avidity-associated $T_{EX\text{-}KLR}$ DEG signature is associated with response to anti-PD-1 in triple-negative breast cancer**

**a**, Schematic of the clinical trial design.

**b–d**, Per-patient mean module scores in $T_{EX}$ cells for the avidity-associated (**b**) $T_{EX\text{-}KLR}$ and (**c**) $T_{EX\text{-}int}$ DEG signatures (from **Fig. 4g**) and (**d**) the relative module scores represented as the Δz-score (Δz-score = z-scored $T_{EX\text{-}KLR}$ module score minus z-scored $T_{EX\text{-}int}$ module score; $n$ = 5 non-responders and 15 responders pre-treatment and $n$ = 7 non-responders and 17 responders post-cycle 1). $P$ values from a Wilcoxon rank-sum test with Benjamini–Hochberg correction are shown.

### Cohort 2: Patient OC03 (pTR2) pre- and post-treatment whole-tissue sections

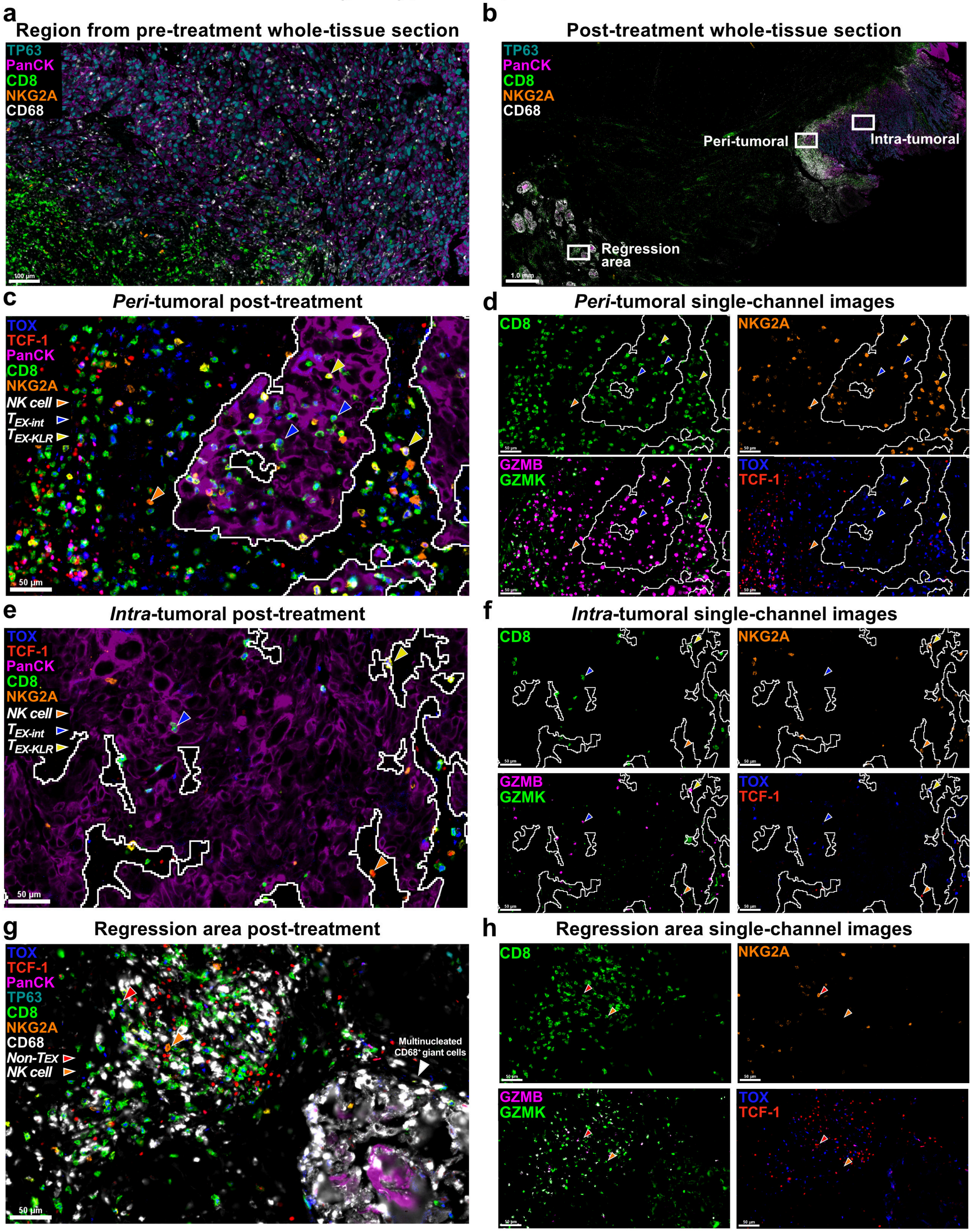

**Extended Data Figure 12. Whole-tissue section immunofluorescence images of Cohort 2 patient OC03 (pTR2) before and after neoadjuvant anti-PD-1 therapy**

**a**, Selected region from the pre-treatment whole-tissue section.

**b,** Post-treatment whole-tissue section. The selected peri-tumoral, intra-tumoral, and regression areas indicated by white boxes are shown below.

**c–h,** Views of the selected post-treatment (**c–d**) peri-tumoral, (**e–f**) intra-tumoral, and (**g–h**) regression areas showing $T_{EX}$ subset marker proteins (*left*; **c, e, g**) and single-channel images (*right*; **d, f, h**). Selected cell types are indicated by colored arrows.

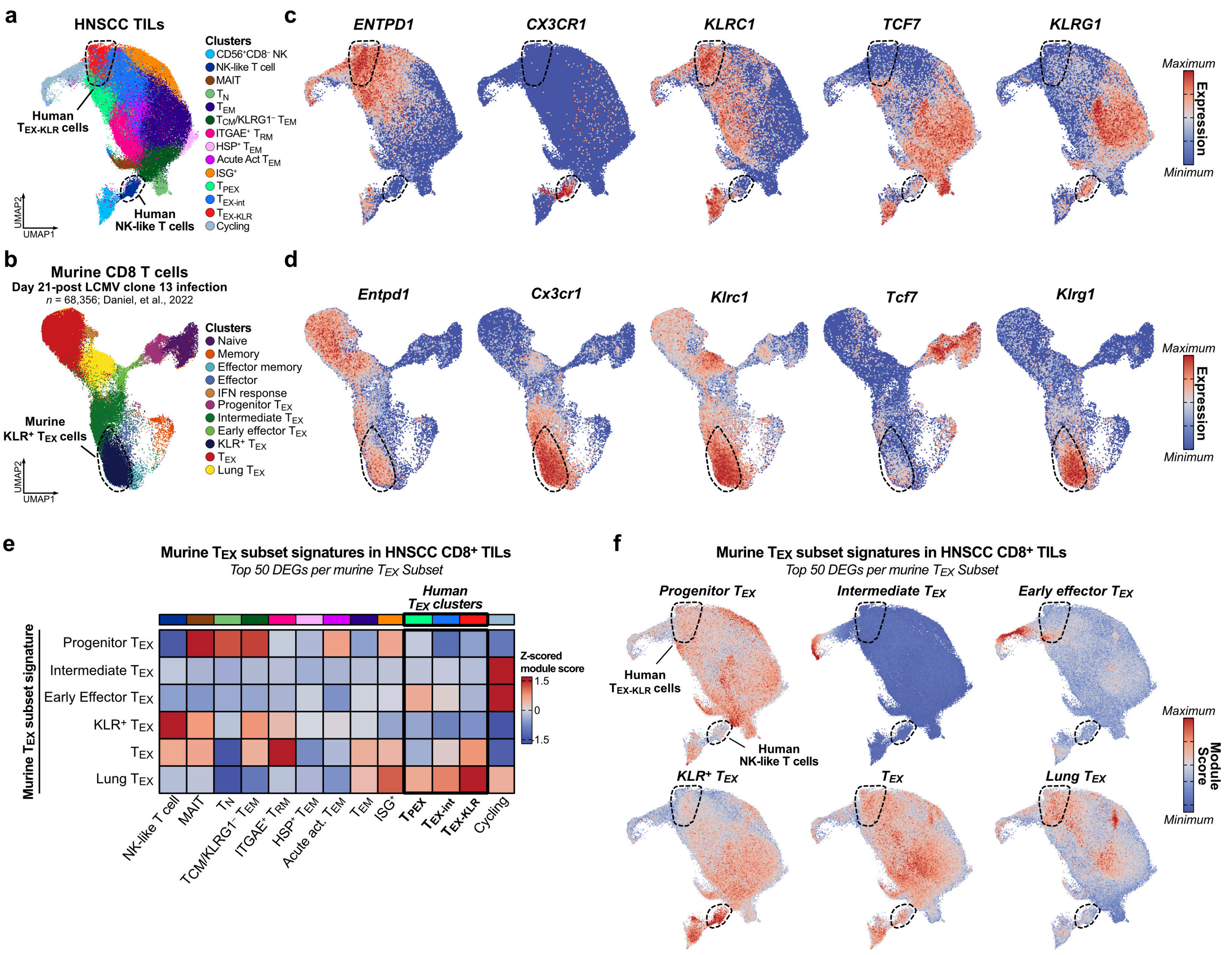
a
HNSCC TILs
Clusters
CD56+CD8− NK
NK-like T cell
MAIT
TN
TEM
TCM/KLRG1− TEM
ITGAE+ TRM
HSP+ TEM
Acute Act TEM
ISG+
TPEX
TEX-int
TEX-KLR
Cycling
Human TEX-KLR cells
Human NK-like T cells
UMAP1
UMAP2
b
Murine CD8 T cells
Day 21-post LCMV clone 13 infection
n = 68,356; Daniel, et al., 2022
Clusters
Naive
Memory
Effector memory
Effector
IFN response
Progenitor TEX
Intermediate TEX
Early effector TEX
KLR+ TEX
TEX
Lung TEX
Murine KLR+ TEX cells
c
ENTPD1
CX3CR1
KLRC1
TCF7
KLRG1
Maximum
Expression
Minimum
d
Entpd1
Cx3cr1
Klrc1
Tcf7
Klrg1
e
Murine TEX subset signatures in HNSCC CD8+ TILs
Top 50 DEGs per murine TEX Subset
Human TEX clusters
Murine TEX subset signature
Progenitor TEX
Intermediate TEX
Early Effector TEX
KLR+ TEX
TEX
Lung TEX
Z-scored module score
1.5
0
-1.5
NK-like T cell
MAIT
TN
TCM/KLRG1− TEM
ITGAE+ TRM
HSP+ TEM
Acute act. TEM
TEM
ISG+
TPEX
TEX-int
TEX-KLR
Cycling
f
Murine TEX subset signatures in HNSCC CD8+ TILs
Top 50 DEGs per murine TEX Subset
Progenitor TEX
Intermediate TEX
Early effector TEX
KLR+ TEX
TEX
Lung TEX
Human TEX-KLR cells
Human NK-like T cells
Maximum
Module Score
Minimum

**Extended Data Figure 13. Human tumor-infiltrating $T_{EX}$ subsets are distinct from murine $T_{EX}$ subsets in the lymphocytic choriomeningitis virus clone 13 chronic infection model**

**a**, UMAP plot of the HNSCC TILs dataset with the $T_{EX\text{-}KLR}$ and the non-$T_{EX}$ NK-like T cell clusters outlined.

**b**, UMAP plot of murine lymphocytic choriomeningitis virus (LCMV) clone 13-specific CD8 T cells at day 21 post-infection from the Daniel *et al*., 2022 dataset. The published UMAP coordinates and clusters are shown, and the murine $KLR^{+}$ $T_{EX}$ cluster is outlined.

**c–d**, Expression of selected murine T cell subset markers in (**c**) HNSCC TILs and (**d**) murine LCMV clone 13-specific CD8 T cells.

**e**, Mean z-scored module scores for the murine $T_{EX}$ subset signatures in the HNSCC TILs CD8 T cell clusters (see **Methods**).

**f**, Per-cell module scores for the murine $T_{EX}$ subset signatures in the HNSCC TILs dataset.
